# A developmental atlas of the mouse brain by single-cell mass cytometry

**DOI:** 10.1101/2022.07.27.501794

**Authors:** Amy L. Van Deusen, Sarah M. Goggin, Corey M. Williams, Austin B. Keeler, Kristen I. Fread, Irene Cheng, Christopher D. Deppmann, Eli R. Zunder

## Abstract

Development of the mammalian brain requires precisely controlled differentiation of neurons, glia, and nonneural cells. To investigate protein-level changes in these diverse cell types and their progenitors, we performed single-cell mass cytometry on whole brain (E11.5/E12.5) and microdissected telencephalon, diencephalon, mesencephalon, and rhombencephalon (E13.5–P4) collected at daily timepoints from C57/BL6 mice. Measuring 24,290,787 cells from 112 sample replicates with a 40-antibody panel, we quantified 85 molecularly distinct cell populations across embryonic and postnatal development, including microglia putatively phagocytosing neurites, neural cells, and myelin. Differentiation trajectory analysis also identified two separate pathways for producing oligodendrocyte precursor cells. Comparison with previous studies revealed considerable discrepancies between protein and mRNA abundances in the developing brain, demonstrating the value of protein-level measurements for identifying functional cell states. Overall, our findings demonstrate the utility of mass cytometry as a high-throughput, scalable platform for single-cell profiling of brain tissue.

## Main

While numerous studies have cataloged cells present in the brain at maturity^1^, many fundamental questions about their development remain unresolved. In particular, the molecular profiles, timing of appearance, and cell-lineage relationships of neural stem cells (NSCs) and intermediate progenitors remain poorly characterized. Mapping the molecular trajectories and cell fate decisions underlying brain development promises to enhance our understanding of developmental disorders such as autism spectrum disorder^2^ and epilepsy^3, 4^, as well as mature-onset diseases that may originate during development, such as schizophrenia^5, 6^ and Alzheimer’s disease^7, 8^.

Previous efforts to identify and characterize cell populations in the central nervous system (CNS) have primarily relied on either immunofluorescence microscopy, which detects a small number of proteins simultaneously, or single-cell RNA sequencing (scRNA-seq), which detects a large number of transcripts simultaneously. scRNA-seq and the related technique single-nuclei RNA sequencing have been applied to characterize the molecular diversity of cells in a wide range of embryonic and postnatal mouse brain tissues^9–25^. However, no study to date has molecularly profiled single cells from every brain region at daily time points across embryonic and postnatal development, and no single-cell study to date has quantified protein instead of mRNA abundance levels in the developing brain.

To address these deficits, we leveraged the high throughput (1 × 10^6^ cells/hour) of single-cell mass cytometry to profile four regions of the developing brain: telencephalon diencephalon, mesencephalon, and rhombencephalon, with daily time points from embryonic day 11.5 (E11.5) to postnatal day (P4). Mass cytometry is a variation of flow cytometry in which abundances of proteins and other biomolecules are quantified at the single-cell level using rare earth metal isotope-labeled antibodies and other affinity reagents^26, 27^. Commercially available metal isotope reagents permit over 40 molecular markers to be measured and quantified simultaneously in each cell by mass cytometry, including cell surface receptors, transcription factors, cytoskeletal proteins, and markers of cell cycle status and viability.

The antibody-based measurements from mass cytometry can directly read out functional biomolecules, unlike mRNA transcripts that do not necessarily correlate with protein abundance – a disconnect most pronounced during dynamic cell transitions^28, 29^ like those occurring in early brain development. Mass cytometry was previously employed to investigate glioma^30–37^, microglia^38–45^, and dorsal root ganglia^46^, but this approach has not yet been applied to study neural cell types in the brain, except for one limited analysis with seven neural-specific markers in a study of obesity-inhibited adult neurogenesis^47^.

To adapt mass cytometry for neural cells from embryonic and postnatal brain tissue, we developed a 40-antibody panel of CNS development-specific markers and optimized cell dissociation techniques for brain tissue. Once this iterative optimization and development process was complete, we profiled brains of C57/BL6 mouse embryos and pups from E11.5, shortly after the secondary vesicles have formed and cortical neurogenesis begins, to P4, when the brain regions have assumed their final morphology, mature neural cell types are present, and synaptic connections are beginning to form^48^.

Using this neural mass cytometry approach, we identified and quantified 85 molecularly distinct cell populations across embryonic and postnatal development in the telencephalon, diencephalon, mesencephalon, and rhombencephalon. These cell populations generally show complementary overlap with those previously identified by scRNA-seq studies^9–11, 13–15, 16, 17–24, 25, 49, 50, 51–55^, although our time-course comparison reveals that mRNA transcript levels do not accurately predict protein abundance during early brain development. Application of URD pseudotime analysis^56^ to map cell differentiation captured classical excitatory and inhibitory neuronal trajectories, and revealed two distinct cell-lineage hierarchies for producing embryonic OPCs. Additionally, our measurements detected putative phagocytic cargo within individual microglia, highlighting their dynamic functions during early brain development. Collectively, our findings, methods, and analytical strategies establish mass cytometry as a platform to identify cell types in the developing brain, as well as the transition states and molecular trajectories underlying their specification.

## Results

### Classification of cells in the developing mouse brain by mass cytometry

To characterize single cells in the developing brain by their protein expression signatures, we first adapted mass cytometry for brain tissue by optimizing dissection and cell dissociation techniques for single-cell analysis (**Methods**), and developed a 40-antibody staining panel for specific cell types in the brain (**Extended Data Table 1)**. The antibody panel included general markers of neural identity (CD24, N-cadherin, Cux1), neuronal development [doublecortin (DCX), TuJ1, NeuN, MAP2, GAD65, Tbr2, NeuroD1, Ctip2, Tbr1], glial development (A2B5, BLBP, GFAP, GLAST, Olig2, OligoO4, Sox10, PDGFRα), leukocytes and microglia (CD45, CD11b, Ly6C, F4/80), and vascular cells (CD31, VCAM, MCAM, PDGFRβ). We also included markers of NSCs and progenitor cells (Sox1, Sox2, nestin, Pax6, PSA-NCAM, CD133, SSEA1), cell signaling and proliferation (Ki67, TrkB, p75NTR), and apoptosis (cleaved caspase 3). As outlined in **Extended Data Table 1**, many of these markers are not restricted to a single cell type, such as BLBP, which is expressed in radial glial cells (RGCs) as well as astrocytes. Antibodies for each marker were conjugated to unique rare earth metal isotopes with Maxpar® X8 Antibody Labeling Kits (Fluidigm, South San Francisco, CA), and then titrated to identify their optimal staining concentrations using known-positive and known-negative control cells on a CyTOF® Helios^TM^ mass cytometer (Fluidigm) (**Extended Data Fig. 1**). Additional details are provided in the Methods section.

Brain samples from C57/BL6 mouse litters were collected at daily timepoints from E11.5 to P4 by dissection of timed-pregnant females (embryonic) and newborn pups (postnatal). Because E11.5 and E12.5 brains are difficult to reliably microdissect, we chose to analyze whole brain samples for these early ages. Brains aged E13.5 and older were microdissected into the telencephalon, diencephalon, mesencephalon, and rhombencephalon (**Fig. 1a**). Each dissected or microdissected tissue was pooled by litter before cell dissociation, and at least two biological replicates (i.e. pooled litters) were analyzed for each age and brain region (**Supplementary Table 1**). After dissection, microdissection, and pooling by litter, each sample was dissociated into a single-cell suspension (**Methods**), briefly incubated with cisplatin as a non-cell-permeant viability stain^57^, fixed with 1.6% paraformaldehyde, and stored at -80°C.

**Fig. 1.**
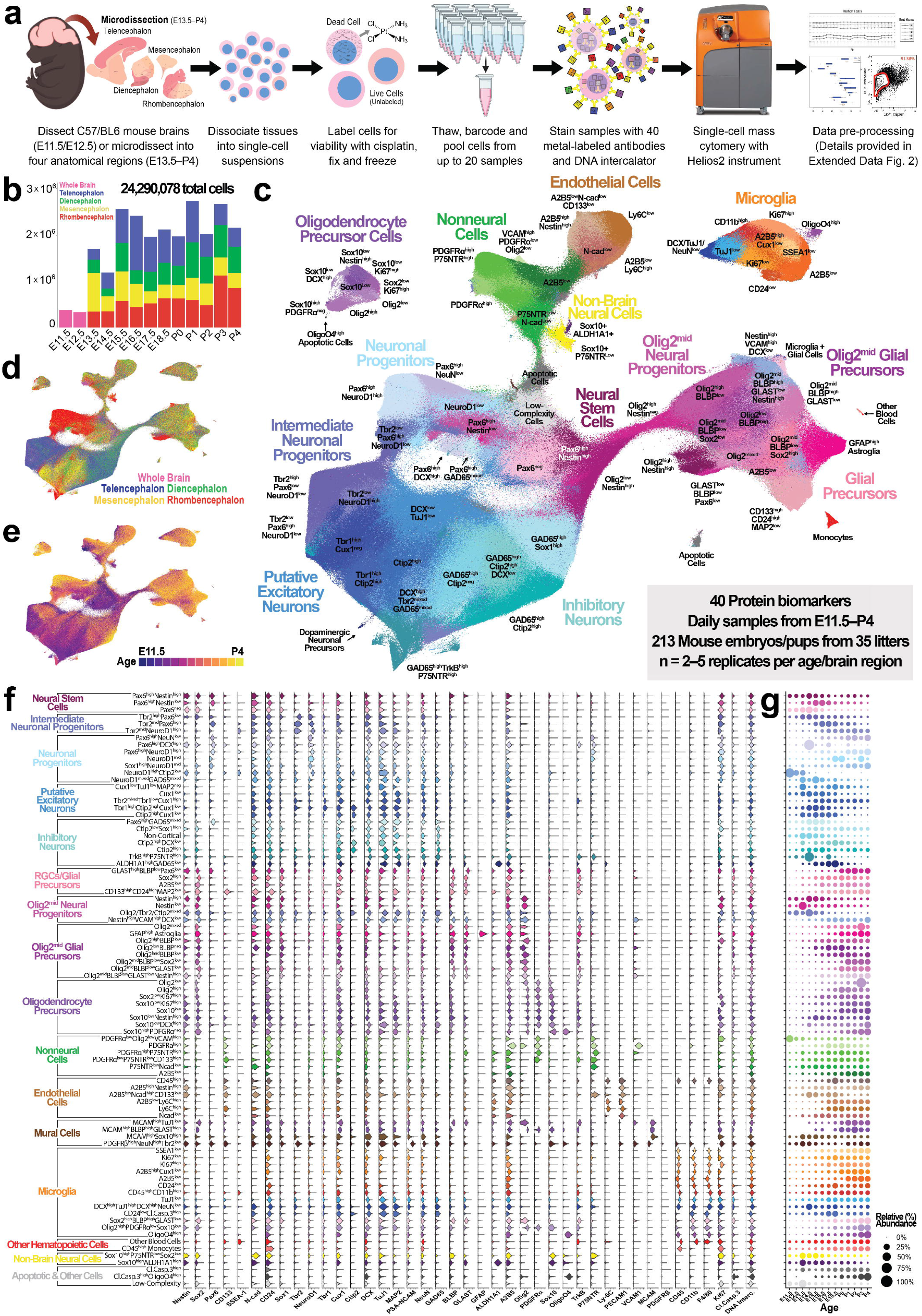
Classification of cells in the developing mouse brain by mass cytometry. **a,** Overview of workflow for isolating and processing single cells from developing mouse telencephalon, diencephalon, mesencephalon, and rhombencephalon for mass cytometry and high-dimensional analysis. **b,** Bar graph showing numbers of cells analyzed for each age (El 1.5-P4) and brain region. **c-e,** UMAP of Leiden clustering all samples colored by cluster (**c**), tissue origin (**d),** and developmental age (**e**). **f,** Violin plot showing expression of all 40 markers in the 85 identified clusters. Clusters are arranged according to class and molecular profile. g, Dot plot showing relative cluster abundances at each developmental age. The data used to generate this plot is available in .csv format in the Source Data for this figure. For **c-g,** n = 2-5 litters per age for a total of 5,750,000 cells from 112 samples.

For mass cytometry analysis, cell samples were thawed and labeled with palladium barcodes^58^, followed by pooling into barcode sets for uniform antibody staining. After mass cytometry measurement, the resulting 37,913,425 cell events were pre-processed to isolate viable single cells. Briefly, pre-processing steps included: 1) bead normalization^59^ (**Extended Data Fig. 2a**); 2) debarcoding^58, 60^ (**Extended Data Fig. 2b**); 3) clean-up gating to remove cell doublets, aggregates, dead cells, debris, calibration beads, and other metal contaminants (**Extended Data Fig. 2c–h**; www.cytobank.org); 4) batch correction^61^ (**Extended Data Fig. 2i**); and 5) marker scaling (**Extended Data Fig. 2j**). These preprocessing steps resulted in 24,290,787 high-quality, viable, singlet cells from 112 samples of C57/BL6 mice (n = 2–5 litters per age) (**Fig. 1b** and **Supplementary Table 1**).

To identify and categorize cell populations in the mouse brain throughout development and across brain regions, we performed Leiden clustering^62^ on all samples and visualized the results on a 2D Uniform Manifold Approximation and Projection (UMAP) layout^63^. An initial round of Leiden clustering yielded 22 clusters (**Extended Data Fig. 3a–b**), which were manually grouped by protein expression profile into six subsets: NSCs and progenitors, neurons and neuronal progenitors, glial progenitors and precursors, oligodendroglia and contaminating non-brain neural cells, endothelial and other nonneural cells, and hematopoietic cells (**Extended Data Fig. 3c**). A second round of Leiden clustering on these six subsets yielded 85 distinct clusters in total (**Fig. 1c**, **Extended Data Fig. 3d,e)**, each with a unique regional identity (**Fig. 1d**, **Extended Data Fig. 3f**), age profile (**Fig. 1e**, **Extended Data Fig. 3g**), and protein expression pattern (**Fig. 1f**). These properties were used to identify the following major cell classes: NSCs (4.99%), intermediate neuronal progenitors (INPs, 4.45%), neuronal progenitors (8.56%), putative excitatory neurons (14.45%), inhibitory neurons (17.45%), RGCs/glial precursors (5.65%), Olig2^mid^ neural progenitor cells (Olig2^mid^ NPCs, 5.01%), Olig2^mid^ glial precursors (13.19%), OPCs (2.61%), nonneural cells (5.96%), endothelial cells (5.18%), mural cells (0.03%), microglia (8.37%), other hematopoietic cells (0.56%), non-brain-derived neural cells (0.39%), apoptotic cells (0.99%). As 2.16% of all cells fell into clusters categorized as low-complexity, 97.84% of all cells examined were classified as specific cell types (**Methods**). Changes in cell abundances across development indicated the following general trends: large proportions of neuronal cells during embryonic ages, vast proliferation of glial cells perinatally, and steady expansion of microglia and nonneural cells across embryonic and postnatal development (**Fig. 1g**).

### Comparison of cell abundances and proliferation across brain regions

To compare developmental trends in the telencephalon, diencephalon, mesencephalon, and rhombencephalon, their relative cell abundances (**Fig. 2; Extended Data Fig. 4a–e**) were plotted for comparison. Similar dynamics were observed across the forebrain (telencephalon and diencephalon), although glial and nonneural cell populations began to increase in relative proportion and diversity two days earlier in the diencephalon (E15.5) compared with the telencephalon (E17.5) (**Fig. 2a,b**). The mesencephalon initially exhibited distributions of inhibitory and excitatory neuronal cells similar to the forebrain, but displayed a more rapid change in the proportions of neuronal and glial cells commencing around E17.5. In contrast to the rest of the brain, the rhombencephalon contained a large complement of glial progenitors by E13.5 and exhibited prolonged periods of both GABAergic and glutamatergic neurogenesis, consistent with previous studies^18, 50^. Although some intralitter and interlitter variation is expected, especially when evaluating developmental phenotypes, analysis of variance for biological replicates indicated high consistency of results for individual clusters (**Extended Data Fig. 4c–e**). Generally, the highest variances were observed in neuronal populations from E11.5–P0 and glial populations from P0– P4, corresponding with periods during which these cells vastly increase in number; in contrast, low variances were observed for OPC populations at all ages.

**Fig. 2.**
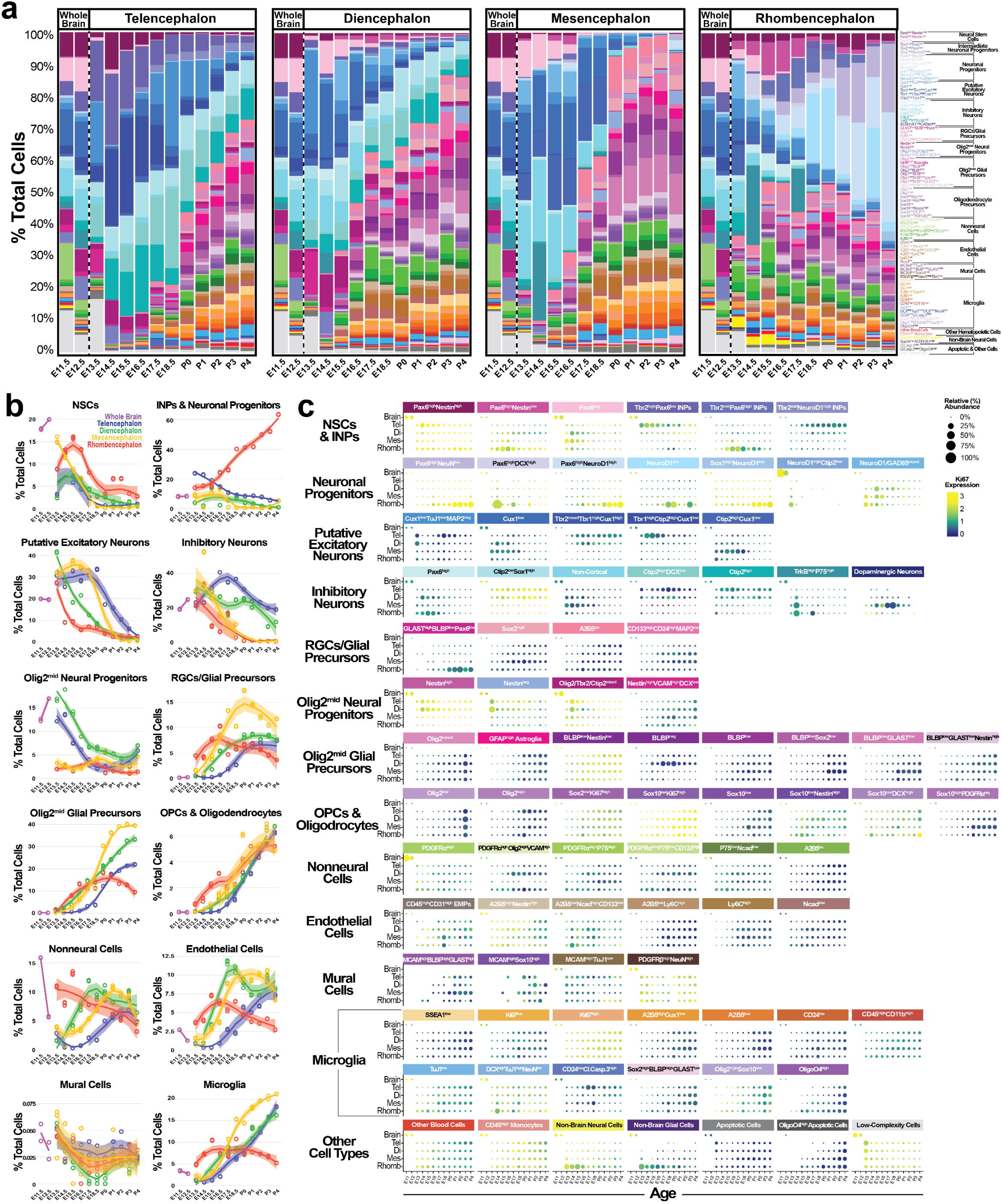
Spatiotemporal profile of cell abundances across the developing mouse telencephalon, diencephalon, mesencephalon, and rhombencephalon. **a**, Relative cluster abundances in the four examined regions of the developing mouse brain. Whole brain samples (El 1.5 and E12.5) were included on each plot for clarity. Note that results for Ell.5 and E12.5 are replicated in each plot to facilitate comparison with that region. **b,** Relative abundances of major cell classes (as shown in **a)** in each brain region from El 1.5-P4. Individual replicates are shown along with Loess curve fitting of the data. The data used to generate the plots in **a** and **b** are available in .csv format in the Source Data for Figs. 1 and 2, respectively. **c,** Relative cluster abundances in each brain region from E11.5-P4 grouped according to major cell class. Dots are colored according to the relative expression level of Ki67, a marker of cell proliferation. n = 2-5 litters per age for a total of 5,750,000 cells from 112 samples.

To directly compare the population dynamics and proliferative status of individual cell populations from **Figure 1** between microdissected brain regions, their relative abundances across development were plotted along with Ki67, a non-binary indicator of cell proliferation^64^ (**Fig. 2c**). For clarity, the term “precursor” has been applied to nonproliferative (Ki67^low^) cell populations, while the term “progenitor” has been reserved for proliferative (Ki67^high^) cell populations. Among NSCs and INPs, we observed a Pax6^neg^ subset in the embryonic diencephalon and mesencephalon, while populations expressing Pax6 and Tbr2 were enriched in the telencephalon and rhombencephalon, where they ultimately produce glutamatergic neuronal lineages^65–67^. Notably, a small fraction of Tbr2^mid^Pax6^high^ INPs in the rhombencephalon was positive for Olig2 (**Fig. 1f**), consistent with previous descriptions of Olig2-positive cells in the embryonic cerebellar ventricular zone and rhombic lip^68, 69^.

In line with the cerebellum containing as many as 70% of neurons in the mouse brain^70^, six of the seven neuronal progenitor populations were observed almost exclusively in the rhombencephalon (**Fig. 2c**). Although the majority presumably go on to produce cerebellar granule cells^71^, the rare Ctip2^high^NeuroD1^high^Olig2^low^ population observed at the earliest ages likely corresponds to early Purkinje cell progenitors^72^. Other putative glutamatergic and inhibitory neuron populations were distinguished by positive or negative GAD65 expression, respectively, although it is worth noting that GAD65 is expressed by GABAergic^73^, dopaminergic^74^, and cholinergic neurons^75^. The remaining non-cortical population displayed widely varying expression (mixed) levels of NeuroD1 and GAD65 (NeuroD1^mixed^GAD65^mixed^) (**Fig. 1f**), indicating this cluster contained a mixture of cells with glutamatergic and GABAergic fates. The two waves of Cux1^low^TuJ1^low^MAP2^neg^ neurons in the telencephalon suggest that this is a transitional population (presumably derived from waves of direct and indirect neurogenesis around E13.5 and E18.5, respectively^76^) that ultimately matures into the observed Cux2^low^, Cux1^high^, and Ctip2^high^ populations.

Three of the six GABAergic inhibitory neuron populations were predominantly observed in the forebrain and expressed Ctip2 (**Fig. 2c**), a classical marker of deep layer subcerebral projection neurons that is also expressed by parvalbumin, somatostatin, and 5-HT3 inhibitory neurons in layers I–V of the neocortex^77^ and Purkinje cells^72^. Of the four Ctip2^neg^ GABAergic subtypes, Pax6^high^GAD65^mixed^ cells in the embryonic diencephalon and rhombencephalon likely correspond to inhibitory neurons in the developing dorsal white matter^78^, while the ALDH1A1^high^GAD65^low^ population likely corresponds to ventral mesencephalic dopaminergic neurons^9, 17, 79^.

RGCs/glial precursors (Olig2^neg^ cells expressing stemness markers such as Sox2, BLBP, and GLAST, but not mature neuronal or glial markers; **Fig. 1f**) were generally nonproliferative (Ki67^low^) (**Fig. 2c**). Sox2^high^ and A2B5^low^ RGCs/glial progenitors exhibited caudal-to-rostral expansion, while a GLAST^high^BLBP^low^Pax6^low^ population was observed almost exclusively in the rhombencephalon and a CD133^high^CD24^high^MAP2^low^ population (likely containing ependymal cells^80^) was primarily observed in the diencephalon and mesencephalon.

The four Olig2^mid^ neural progenitor clusters likely represent cells that can give rise to neuronal and glial lineages. Two of these populations, Olig2^mid^Nestin^high^ and Olig2^mid^Nestin^low^ NSCs, expressed no mature neural markers (**Fig. 1f**) and were principally observed in the forebrain. An Olig2/Tbr2/Ctip2^mixed^ population most abundant at E14.5 likely corresponds to early basal forebrain cholinergic neuronal progenitors^81^. Finally, a Nestin^high^VCAM^high^DCX^low^ population exhibiting caudal-to-rostral expansion corresponds to uncommitted Olig2-expressing neural progenitors (**Fig. 2c**).

Eight molecularly similar glial precursor clusters expressing Olig2 were designated Olig2^mid^ glial precursors, including a GFAP^high^ population that expanded from E16.5 in all brain regions consistent with previous descriptions of “Olig2-lineage” or “OPC-derived” astrocytes^82–85^ (**Fig. 2c**). Thus, these cells likely represent various states of intermediate glial cells (iGCs) with distinct astrocytic or oligogenic potentials^86^. Notably, only BLBP^low^Nestin^low^ Olig2^mid^ glial precursors were proliferative.

OPCs and oligodendrocytes were characterized by high Olig2 and PDGFRα expression, although one PDGFRα^neg^ OPC population expressing Sox10 was also identified (**Fig. 1f**), in agreement with previous reports^87, 88^. Consistent with multiple waves of OPC generation in distinct ventricular zones during embryonic development^89^, expansion of OPC populations varied spatiotemporally (**Fig. 2c**). However, the eventual presence of OPCs with similar protein expression profiles by P4 in all microdissected brain regions is consistent with previously reported transcriptional uniformity of postnatal OPCs^90^.

Nonneural cells (generally defined by a lack of Sox2 and neural cell markers) could not be precisely identified by our CNS-focused antibody panel, but likely contain brain fibroblasts, vascular smooth muscle cells, and other cerebrovascular cells. All six endothelial cell populations generally exhibited caudal-to-rostral expansion, including a rare subset of CD45^low^PECAM^high^ cells (**Fig. 2c**) resembling erythromyeloid progenitors (EMPs) capable of producing erythroid, myeloid, and endothelial cell lineages^91^. Mural cells were characterized by MCAM and PDGFRβ expression, but also contained neural markers, likely due to incomplete removal of neural cell debris from their surface (**Fig. 2c**). Microglia were characterized by CD45 and CD11b expression, although many subsets also contained various neural markers (**Fig. 1f**), likely due to the established phenomena of microglial phagocytosis^92^, although we cannot rule out cell surface-associated debris. Finally, two Sox10^high^ populations observed exclusively in whole brain and rhombencephalon tissues likely correspond to contaminating spinal cord and neural crest-derived cells (**Fig. 2c**).

### Comparison of protein and mRNA expression patterns in the developing mouse brain

To investigate how well mRNA expression predicts protein abundance in the developing brain, we compared our antibody-based mass cytometry measurements with two sets of age-matched and tissue-matched mRNA measurements: *in situ* hybridization (ISH) from the Allen Developing Mouse Brain Atlas^93^ (https://developingmouse.brain-map.org, **Extended Data Fig. 5a**) and scRNA-seq from Linnarsson and colleagues^53^ (**Extended Data** **Fig. 5b**). Our antibody panel has 34 proteins with directly comparable cognate mRNAs (25 of which are included in the Allen Brain Atlas), but other markers are not directly comparable, such as cleaved caspase 3 and the ganglioside A2B5 (**Extended Data Fig. 5c**).

Because protein and mRNA are each subject to numerous regulatory mechanisms that control their synthesis and degradation, there is no universal method to accurately predict protein abundances from mRNA expression levels, or vice versa^28^. To investigate the relationship between protein and mRNA levels in the developing brain, we compared percentages of cells expressing each of these protein-mRNA cognate pairs (34 for scRNA-seq, 25 for ISH) and their normalized mean abundance levels (**Fig. 3a**). In many cases, the highest protein levels observed were delayed compared with the highest mRNA levels, such as for Ctip2, nestin, DCX, MAP2, PDGFRɑ, TrkB, and N-cadherin. In other cases, the highest protein levels actually preceded the highest mRNA levels, such as for Olig2, PDGFRβ, GLAST, VCAM, and MCAM. Inconsistent results between scRNA-seq and *in situ* hybridization further complicate the conclusions drawn from this protein-mRNA comparison, although the single-cell dissociation step shared by mass cytometry and scRNA-seq means they are subject to similar experimental constraints, and may therefore be more directly comparable.

**Fig. 3.**
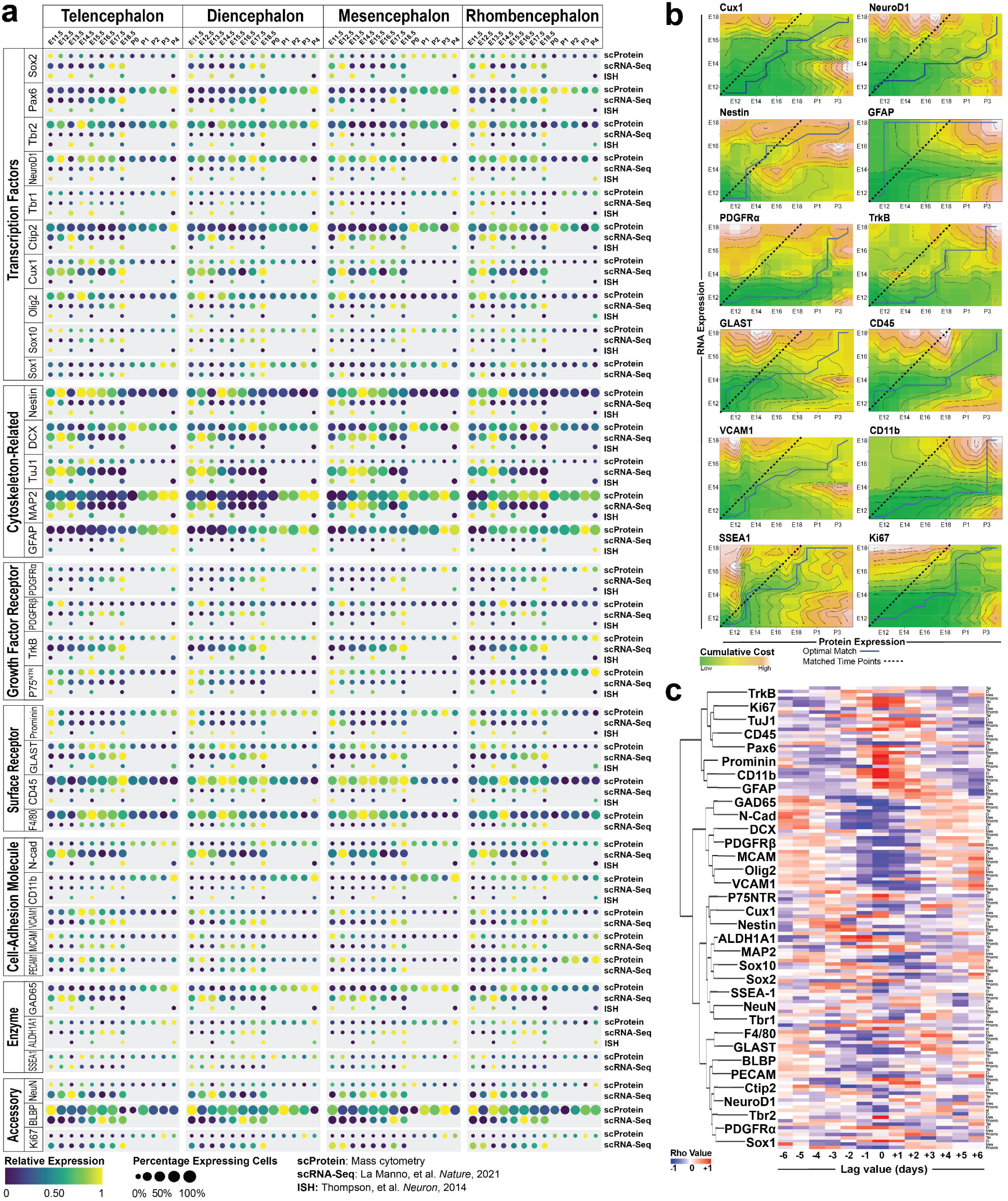
Comparison of protein and mRNA expression profiles in the developing mouse brain. **a,** Dot plot comparing protein/RNA expression from Allen Brain Atlas and La Manno *(Nature,* 2021), grouped according to the general class of protein encoded by each gene. Data for genes or ages not shown were unavailable at the time of publication. Note that results for El 1.5 and E12.5 are replicated in each plot because they were normalized separately for comparison with each region. **b,** Dynamic time-warp analysis comparing relative RNA and protein expression levels in expressing cells. Comparisons with low cumulative costs (indicating similarity) are colored green, while those with high cumulative costs (indicating dissimilarity) are colored orange. Solid blue lines indicate the optimal alignment path calculated by the algorithm, while dashed lines indicate matched time points (i.e., null hypothesis of RNA= protein expression with a lag of 0 days). **c,** Heatmap showing cross correlations of differenced mean expression measurements for each overlapping gene in each brain region. n = 2-5 litters per age for a total of 5,750,000 cells from 112 samples.

To investigate relationships between RNA and protein abundance during embryonic and early postnatal brain development, we performed dynamic time warp analysis^94^ on data for each protein-RNA pair. The resulting optimal alignments between relative protein and RNA expression suggested three general trends: consistent lag (e.g., Cux1, nestin, TrkB, and SSEA-1), increasing lag (e.g., NeuroD1, GLAST, CD45, VCAM, and Ki67), or unpatterned lag (e.g., GFAP, PDGFR⍺, and CD11b) (**Fig. 3b**). It should be noted that because scRNA-seq is prone to dropout of low-abundance RNA transcripts, evaluation of protein/RNA relationships for certain markers (e.g., transcription factors) may be less reliable. To quantify the kinetics of expression lag and determine whether they are shared between specific protein types, we calculated the cross correlation of relative expression changes across overlapping timepoints (**Fig. 3c**). The largest absolute correlation values were generally observed between 0 and 3 days, with PDGFRβ, MCAM, and Olig2 showing strong negative Rho values, and prominin, CD11b, and GFAP showing strong positive Rho values centered around day 0.

### NSC/RGC and intermediate progenitor populations in the Sox2^+^Nestin^+^ compartment

We next investigated how the differentiation potential of NSCs/RGCs and intermediate progenitors changes over the course of development and between brain regions. Because cells positive for stem cell markers were distributed across a large number of clusters (**Extended Data Fig. 3h**), this subset of cells was isolated by gating on two canonical NSC markers: Sox2^95–100^ and nestin^101–103^ (**Fig. 4a**). Sox2^+^Nestin^+^ cells displayed the highest abundance at E14.5, reached a nadir around E18.5/P0, and increased postnatally (P1–P4) in all brain regions, although relative abundances of postnatal NSC/RGCs varied between regions (**Fig. 4b**). The first embryonic wave of NSCs expressed neurogenic markers such as Tbr2, NeuroD1, DCX, and TuJ1, while the second postnatal wave expressed glial markers such as BLBP, GLAST, and GFAP (**Fig. 4c**), consistent with a neurogenic to gliogenic switch of NSC/RGC fates in the developing brain before _birth104,105_.

**Fig. 4.**
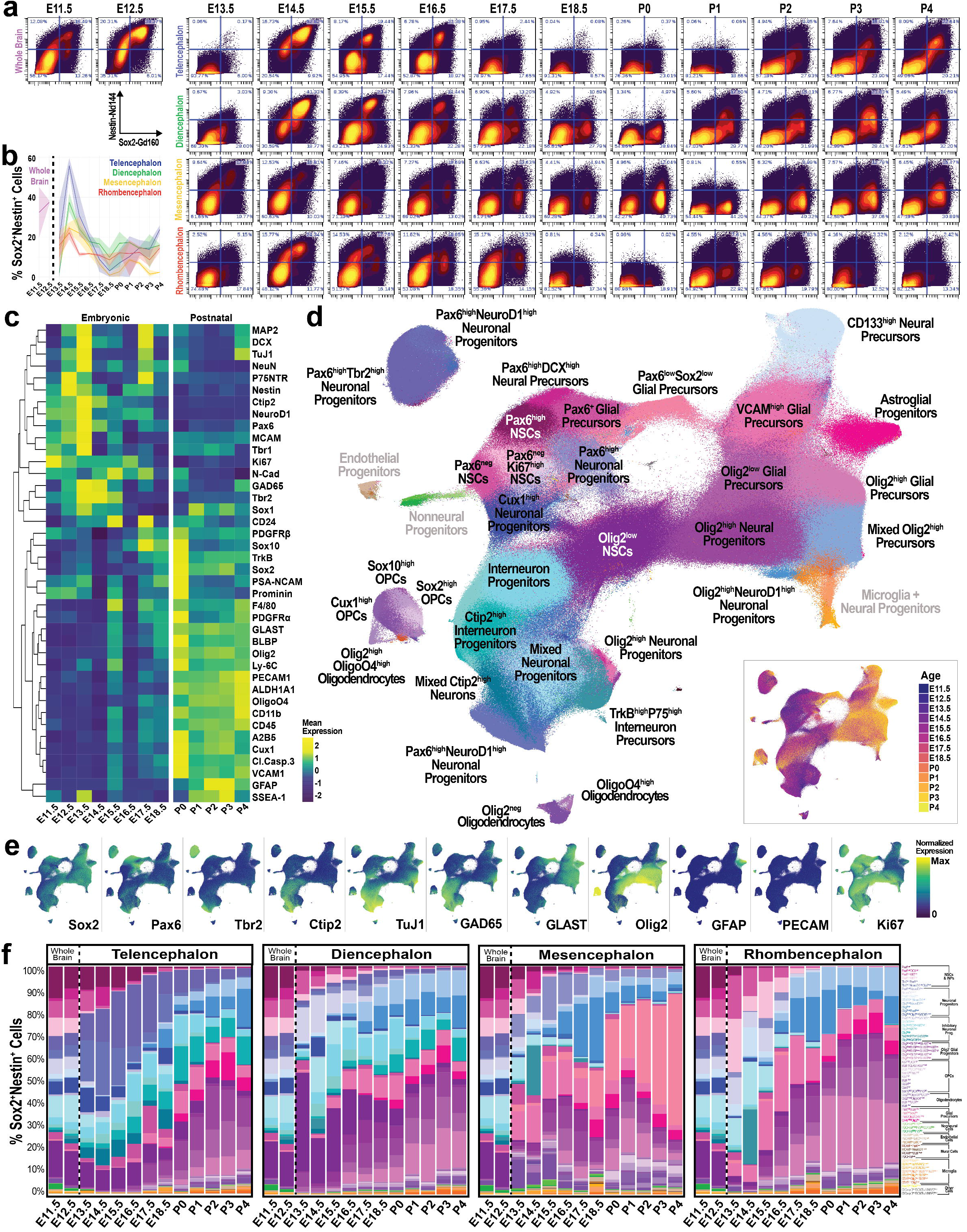
Survey of cells with stem-like properties in the developing mouse brain. **a**, Biaxial plots showing gating of Sox2+Nestin+ cells with CytoBank. Percentages of cells in each quadrant are shown. Cells in the upper right quadrant were included in analyses. **b,** Relative abundances of Sox2+Nestin+ cells in each brain region across time. Mean (solid line) shown along with range. **c,** Heatmap showing mean expression levels of all 40 markers in the antibody panel for each developmental age. **d,** UMAP of Leiden clustering of Sox2+Nestin+ cells colored according to cluster identity. Inset shows UMAP colored according to developmental age. **e,** UMAPs colored according to expression levels of key markers. **f,** Stacked bar graphs showing relative abundances of Sox2+Nestin+ clusters in each brain region from El 1.5-P4. Note that results for El 1.5 and E12.5 are replicated in each plot to facilitate comparison with that region. The data used to generate these plots is available in .csv format in the Source Data for this figure. n = 2-5 litters per age for a total of 3,253,024 cells in 112 samples.

To identify cell populations in the NSC/RGC compartment, we performed Leiden clustering on Sox2^+^Nestin^+^ cells (**Extended Data Fig. 6**). As a result, 43 neural cell populations were identified, including three distinct clusters of NSCs (Pax6^high^, Olig2^low^, and Pax6^neg^), two INP clusters, 17 clusters of neuronal progenitors, and 22 clusters representing various glial cell populations (**Fig. 4d**; **Extended Data Fig. 6c–e**). Generally, these clusters molecularly resembled those described in Figure 1. In addition, three PECAM^high^ populations were identified (**Fig. 4d,e, Extended Data Fig. 6c–d)**, two of which had low Sox2 expression corresponding to nestin-expressing endothelial progenitor cells^106^. The third PECAM^high^ population expressed high levels of Sox2, as well as Olig2, BLBP, and GLAST, potentially representing tissue-resident endothelial cell progenitors^107^. Although these cells have not previously been reported to express glial markers, if endothelial cells and neural cells can arise from a common neural progenitor^107, 108^, it is plausible that PECAM expression overlaps with RGC/glial markers during development. Alternatively, these cells could represent incompletely dissociated endothelial cells and/or glial cells, which interact to form the blood-brain barrier^109^. In addition, small populations of mural cells, microglia, apoptotic cells, and contaminating non-brain cells were observed (**Fig. 4d,f**; **Extended Data Fig. 6a–d**).

Comparison of relative Sox2^+^Nestin^+^ cluster abundances across brain regions revealed that uncommitted Pax6^high^, Pax6^low^, and Olig2^mid^ NSC/RGC-like cell populations were maintained in all brain regions from E11.5 to P4, although at relatively low abundances postnatally (**Fig. 4f**, **Extended Data Fig. 6d**). The forebrain exhibited similar cell population dynamics with two major exceptions: 1) expansion of Olig2-expressing neuronal and glial progenitors began two days earlier in the diencephalon, around E15.5; and 2) more than 30% of cells in the telencephalon from E13.5– E18.5 were Tbr2^high^ INPs, a cell population not observed in other brain regions (**Fig. 4f**). Compared with the forebrain, GAD65-expressing neuronal progenitors were far less prevalent in the mesencephalon and rhombencephalon during late embryonic and postnatal ages. Moreover, in the mesencephalon, a VCAM^high^Pax6^mixed^Sox2^mid^ population that was not observed in other brain regions rapidly expanded from E17.5 to comprise over 50% of all Sox2^+^Nestin^+^ cells by P4. Similar VCAM^high^ cells reported at this stage have been described as a quiescent NSC/RGC population that persists until adulthood in the lateral ventricles^110^ and neocortex^14^. At the cell-subtype level, our clustering analysis generally recapitulates the progressive switch from neurogenic to gliogenic NSCs/RGCs^111^. However, our results also demonstrate that an array of glial progenitors are already present at E13.5, while neuronal progenitors persist until after P0, just at much lower relative abundances in the forebrain and mesencephalon (**Fig. 4f**). These findings are consistent with previous reports^9, 14, 16, 18, 20, 21, 50, 53, 54^, including clonal neuron, astrocyte, and oligodendrocyte progeny arising from individual RGCs genetically labeled between E10 and E13^105^.

### Differentiation trajectories of Sox2^+^Nestin^+^ NSC/RGCs and intermediate progenitors

To investigate neurogenic and gliogenic differentiation trajectories of NSC/RGCs and intermediate progenitors, we applied URD pseudotime analysis^56^ to analyze Sox2^+^Nestin^+^ cells sampled and pooled from all brain regions and ages (**Extended Data Fig. 7a–d**). We chose Pax6^high^ NSCs from E11.5 whole brain as the root cells because Pax6 expression commences at the earliest point of CNS development (∼E8) and acts upstream of many other factors defining neural fates (including Tbr2^66^, Olig2^112^, and BLBP^113^), while neural cell populations with mature expression profiles at P0–P4 were chosen as the tip cells (**Fig. 5a**). In the resulting URD dendrogram, the first branchpoint separates two neuronal trajectories (cortical INPs destined to generate pyramidal neurons and Pax6^high^NeuroD1^high^ neuronal progenitors destined to produce cerebellar granule cells) from all other neural progenitors according to varying expression levels of Pax6, Tbr2, and DCX (**Fig. 5b–d**). The second branchpoint separates the remaining neuronal progenitors from glial progenitors by expression of CD24 and DCX, both of which were highly expressed in the former and low in the latter. The neuronal branch (segment 3) splits further into Olig2^high^Ctip2^high^ neuronal precursors and various inhibitory progenitor/precursor populations according to expression of Olig2, GAD65, MAP2, NeuN, and Ki67. Paradoxically, GAD65^high^Ctip2^low^ and GAD65^high^Ctip2^high^ inhibitory neuron precursors (segments 7 and 8) were NeuN^high^ embryonically and NeuN^low^ postnatally (**Extended Data Fig. 7e**). One possible explanation is that phosphorylation or protein-protein interactions of NeuN masked the epitope targeted by our NeuN antibody^114^. Analysis of the contribution of each brain region to URD branches (**Fig. 5e**) revealed distinct regional biases of these neuronal progenitors: INPs (segment 26) were primarily located in the telencephalon, Pax6^high^NeuroD1^high^ INPs (segment 25) were primarily located in the rhombencephalon, and all GAD65^high^ inhibitory neurons (segments 3–11) were primarily located in the forebrain.

**Fig. 5.**
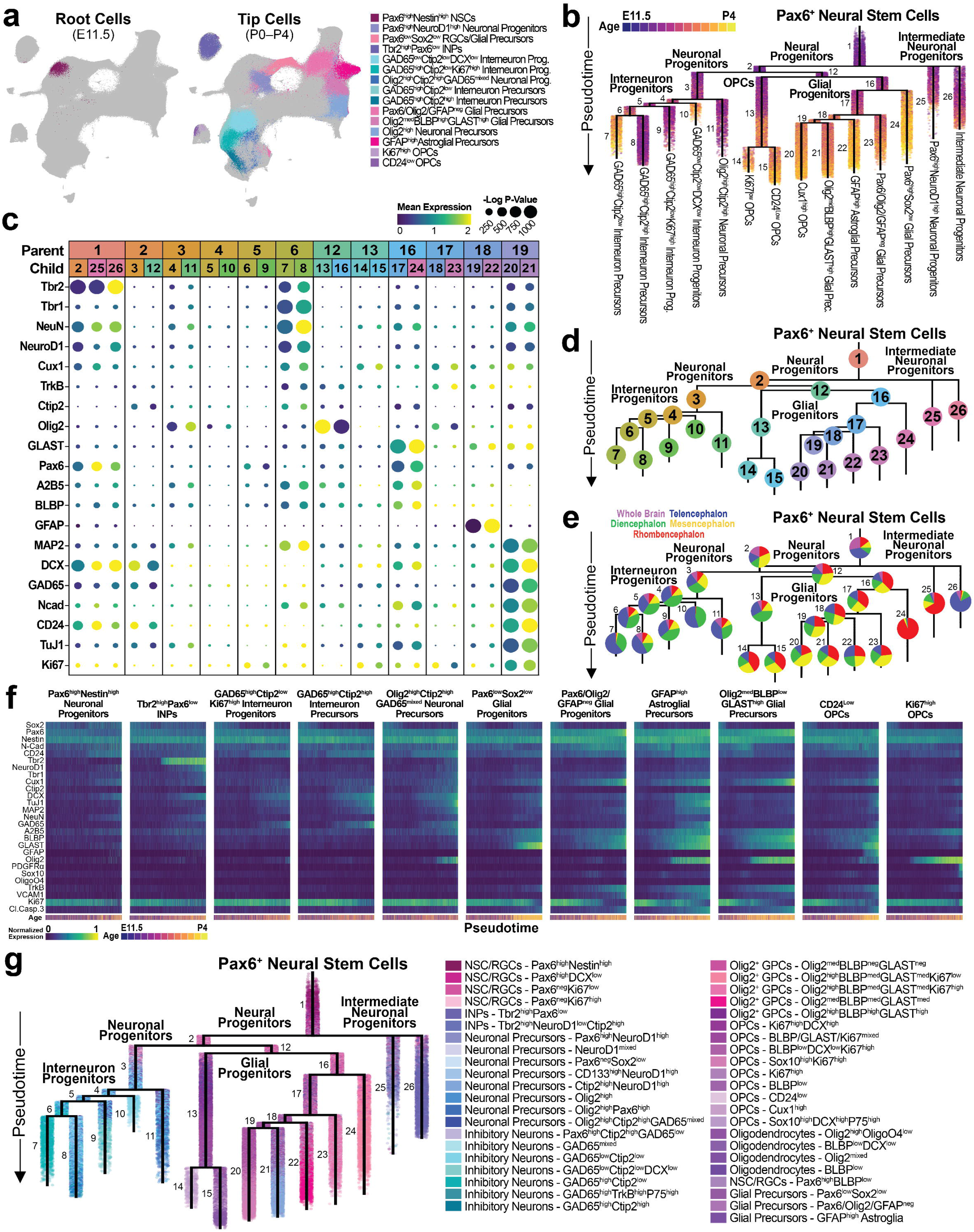
Exploring neural cell fates in the developing mouse brain with pseudotime-based analysis of molecular trajectories. **a**, UMAP of Sox2+Nestin+ cells chosen at the root and tips for URD analysis. **b,** URD dendrogram colored by developmental age. Numbers indicate branch numbers. **c,** Dot plot showing expression of markers key for the division of each branchpoint in URD analysis. **d,** URD dendrogram colored by branchpoint. **e,** URD dendrogram with pie charts showing relative abundance of cells from each brain region for that branch. **f,** Heatmaps colored by marker expression for select trajectories identified by URD analysis. **g,** URD dendrogram colored according to Leiden clustering (shown in Fig. 4). n = 2-5 litters per age for a total of 60,955 cells from 112 samples.

The glial branch (segment 12) of the URD lineage hierarchy splits into two sub-branches: unipotent Olig2^high^ cells that only give rise to OPCs, and bipotent Olig2^low^ cells that differentiate into OPC and astroglial lineages (**Fig. 5b**). Olig2^high^ committed OPC progenitors (segment 13) only branch again much later in pseudotime, while Olig2^low^ glial cells (segment 16) branch early in pseudotime to separate Pax6^high^Sox2^high^ glial precursors from the remaining glial cells. These remaining glial cells subsequently branch into Pax6/Olig2/GFAP^neg^ glial progenitors, GFAP^high^ astroglial precursors, Olig2^mid^BLBP^high^GLAST^high^ glial precursors, and Cux1^high^ OPCs separated by expression of GLAST, BLBP, TrkB, GFAP, DCX, and Ki67 (**Fig. 5c,d**). By this analysis, segments 16–18 represent bipotent iGCs capable of producing both astrocytes and oligodendrocytes^111, 115^. In contrast to neuronal progenitors, cells in glial trajectories (segments 12– 23) more evenly represented the four microdissected brain regions, with the exception of BLBP^high^GLAST^high^ RGCs (segment 24), which were observed almost exclusively in the rhombencephalon (**Fig. 5e**) and likely represent Bergmann glia^116^. The abundances of OPCs in three of the four terminal OPC branches (segments 14, 15, and 21, but not 20) displayed a pattern of rhombencephalon > mesencephalon > diencephalon > telencephalon, indicating a generally caudal-to-rostral pattern of OPC gliogenesis.

Analysis of protein levels along URD pseudotime trajectories revealed the relative timing and sequence of molecular transitions contributing to cell specification (**Fig. 5f**, **Extended Data Fig. 7e**). As expected, the INP trajectory was defined by Tbr2 expression; inhibitory neuron trajectories were distinguished by GAD65 and neural filaments (commencing with DCX); and glial precursors sequentially expressed A2B5, BLBP, and GLAST, followed by GFAP in astroglial precursors and Olig2 in Olig2^med^BLBP^high^GLAST^high^ glial precursors. Although all OPCs expressed low levels of A2B5, the two populations of direct-differentiating OPCs did not proceed through a BLBP^high^GLAST^high^ intermediate state. Overlaying all Sox2^+^Nestin^+^ Leiden clusters from **Fig. 4d** onto the URD dendrogram further demonstrates the segregation of glial progenitors in direct-differentiating and iGC-dependent OPC trajectories (**Fig. 5g**). Collectively, these findings indicate two distinct differentiation trajectories for generation of OPCs in the brain: 1) a direct pathway by which Olig2-expressing NSCs increase levels of OPC/oligodendrocyte markers, but not generic glial proteins; and 2) an indirect pathway by which Olig2-negative NSCs/RGCs (e.g. Pax6 cortical NSCs) first give rise to BLBP- and GLAST-expressing iGCs that subsequently produce Olig2-expressing OPCs. Because the URD algorithm has no mechanism to describe convergent cell trajectories, application of other single-cell trajectory analysis methods^117^ may be necessary to determine if these two trajectories ultimately produce molecularly distinct OPCs and oligodendrocytes later in development – especially because postnatal OPCs have been reported to be transcriptionally convergent^90^.

### Differentiation trajectories and molecular dynamics in the telencephalon

Expanding our trajectory analysis beyond progenitors to include the maturation of neuronal and glial cells, we performed URD pseudotime analysis on all non-hematopoietic cells from E11.5– E12.5 whole brain and E13.5–P4 telencephalon samples using Pax6^high^ RGCs from E11.5 as the root, and twelve terminal populations from the P4 telencephalon as tips (**Fig. 6a,b, Extended Data Fig. 8a–f**). The resulting cell lineage hierarchy first separated nonneural cells and neural cells before further branching into major cell classes (**Fig. 6c**), with neuronal trajectories enriched during embryogenesis, and glial and nonneural trajectories enriched postnatally (**Fig. 6d**).

**Fig. 6.**
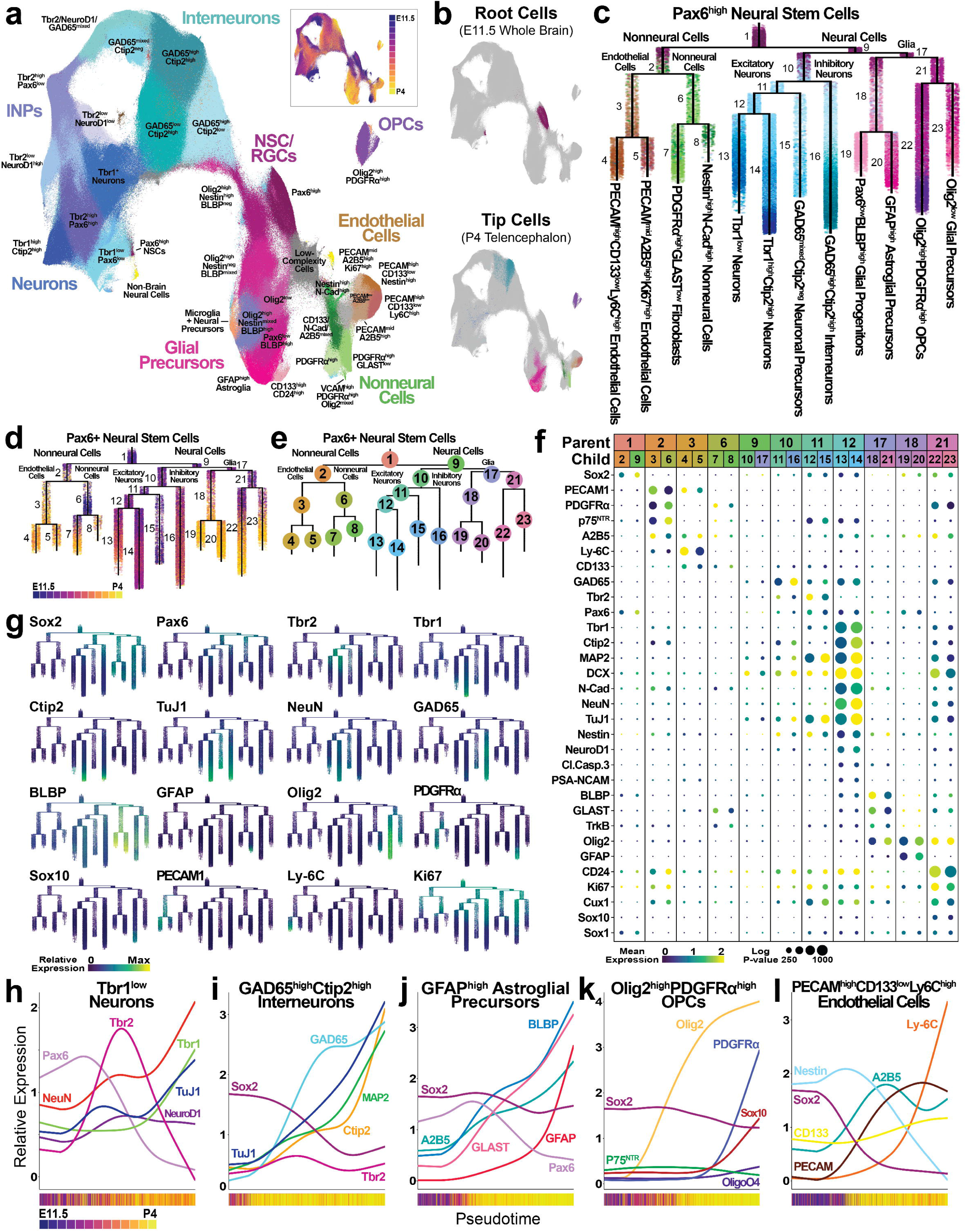
Identifying molecular trajectories underlying cell specification in the developing mouse telencephalon. **a**, UMAP of Leiden clustering of cells in the mouse telencephalon from El 1.5-P4 colored according to cluster identity. Inset shows UMAP colored according to developmental age. n = 2 litters per age for a total of 3,048,448 cells from 26 samples. **b,** UMAP oftelencephalon cells chosen as the root and tips for URD analysis. **c,** URD dendrogram colored according to Leiden cluster identity. **d,** URD dendrogram colored by developmental age. **e,** URD dendrogram with branch numbers indicated. **f,** Dot plot showing expression of markers key for the division of each branchpoint in URD analysis. **g,** URD dendrograms colored according to expression levels of select markers. **h-1,** Relative expression levels of proteins key for defining select molecular trajectories with URD analysis. Bars underneath graphs show median developmental ages of cells within each corresponding pseudotime bin. n = 2 litters per age for a total of 61,006 cells from 26 samples.

The neural branch initially split into DCX^high^ neuronal progenitors (segment 10) and DCX^low^ glial progenitors (segment 17), which were observed primarily from E11.5–E18.5 and P0–P4, respectively (**Fig. 6d–f**). Neuronal trajectories first separated GAD65^high^Ctip2^high^ inhibitory neurons (segment 16) from Nestin^high^ progenitors (segment 11), which split further into GAD65^mixed^Ctip2^neg^ neuronal precursors (segment 15) and Tbr2^high^ INPs (segment 12). The two populations of pyramidal neurons included Ctip2^high^ deep-layer neurons expressing high levels of DCX, TuJ1, and MAP2 (segment 14), as well as Tbr1^low^ upper-layer neurons (segment 13) that appeared less mature, consistent with inside-out formation of the cerebral cortex^118^. As observed for Sox2^+^Nestin^+^ cells, glial trajectories first separated into BLBP^high^GLAST^high^ glial progenitors and Olig2^mid^ glial precursors (segments 18 and 21, respectively). The former trajectory branched into GFAP^high^ astroglial precursors and Pax6^low^BLBP^high^ glial precursors, while the latter split into Olig2^high^PDGFRɑ^high^ OPCs and Olig2^mid^ glial precursors (segments 22 and 23, respectively). Global analysis of marker expression on the URD lineage hierarchy (**Fig. 6g**) revealed additional molecular transitions, including a general shift from Ki67^high^ to Ki67^low^ populations that coincided with increasing expression of mature neuronal or glial markers.

Inspection of key proteins across the URD pseudotime trajectories provides improved resolution on their dynamics and order of expression. For example, we observed sequential expression of Pax6, Tbr2, NeuroD1, and Tbr1 in the Tbr1^low^ neuron trajectory (**Fig. 6h**), similar to previous reports on glutamatergic neuron development in the dorsal telencephalon^66, 119^. On closer inspection, we observed that Pax6 does not decrease until after Tbr2 is elevated, resulting in a subset of Pax6^high^Tbr2^high^ cells. Moreover, NeuN and TuJ1 began to rise with Tbr2, but then paused and did not increase further until Tbr2 decreases, consistent with Tbr2 temporarily inhibiting neuronal maturation by transcriptionally repressing genes associated with axonal growth and dendritic complexity^120^. A similar pause in neuronal maturation was observed in the development of GAD65^high^Ctip2^high^ GABAergic neurons (**Fig. 6i**). Produced by RGCs in the ventral telencephalon, these cells do not traverse through Pax6, Tbr2, and Tbr1 stages; however, their trends for GAD65, Ctip2, and MAP2 expression suggest that a similar mechanism of temporary transcriptional repression may occur during maturation of inhibitory neurons.

Sox2 expression decreases to undetectable levels during neuronal development and maturation, but expression of this stem cell-associated transcription factor was maintained throughout astroglial and OPC differentiation trajectories (**Fig. 6g,j,k**). In the astroglial trajectory, A2B5, BLBP, and GLAST all increased with similar kinetics, while GFAP increased at a slower rate until Pax6 levels dropped, at which point GFAP rapidly increased (**Fig. 6j**). While previous studies indicate that OPCs sequentially express Olig2, Sox10, PDGFRɑ, and OligoO4 as they mature^121–,123^, we observed simultaneous increases in Sox10 and PDGFRɑ protein levels (**Fig. 6k**), perhaps due to translational control or protein degradation preventing earlier accumulation of Sox10.

Whether endothelial cells making up the brain vasculature are solely derived from sprouting of existing endothelial cells^124^ or can also arise locally from EMPs^91, 125, 126^ and/or cells expressing Pax6^127^ or Sox2^107^ is a disputed^128^ and active area of research. To explore the possibility that nonneural cells in the telencephalon arise from Pax6^high^ NSCs/RGCs, we included PECAM^high^CD133^low^Ly6C^high^ and PECAM^mid^A2B5^high^Ki67^high^ endothelial cells (segments 4 and 5), as well as PDGFRɑ^high^GLAST^low^ and Nestin^high^Ncad^high^ nonneural cells (segments 7 and 8), as tips in our URD analysis. It should be noted that the URD algorithm will attempt to generate a path for each user-defined tip, but excludes cells for which it finds no reliable molecular path.

Although some nonneural cells were excluded from analysis (“NA” in **Extended Data Fig. 8j**), most were assigned to a branch that split into endothelial (segments 3–5) and nonneural (segments 6–8) lineages based on expression of PECAM and A2B5 in the former, and PDGFRα and P75NTR in the latter (**Fig. 6e–g**). URD trajectory analysis predicted increases in A2B5 and PECAM coinciding with the loss of Sox2 and nestin in cells differentiating into PECAM^high^CD133^low^Ly6C^high^ endothelial cells (**Fig. 6l**), consistent with a previous description of nestin downregulation during endothelial cell differentiation in the brain^129^.

### Microglial expansion and putative phagocytic cargoes in the developing mouse brain

To investigate subsets of microglia and other hematopoietic cells in the developing brain, CD45^+^ cells from all brain regions and ages were selected by 2D gating (**Extended Data Fig. 9a**), partitioned by Leiden clustering to identify cell subtypes, and visualized with a 3D UMAP layout to improve visualization of cell clusters (**Fig. 7a**). Microglia were identified as CD45^mid^CD11b^high^ cells, but our ability to further specify microglial subtypes was restricted by our antibody panel including only four microglia-specific markers, as opposed to the microglia-focused panels used in previous mass cytometry studies^38–40, 42, 45, 130–132^. However, while lacking in microglia-subtype specificity, our antibody panel enables detection and quantification of putative phagocytic cargoes, allowing us to divide microglia into two functional subtypes: “unladen” without cargo, and “cargo-laden” containing neuronal or glial markers (**Fig. 7a**). Comparison of the microglial marker CD11b with key markers of neuronal cells (Tbr2, NeuroD1, and TuJ1) and various glial populations (Sox2, BLBP, Olig2, and OligoO4) revealed discrete patterns across cargo-laden microglial clusters, whereas these markers were generally absent in all unladen microglia clusters (**Fig. 7b**). This observation is similar to that of presumptive cargos in immature satellite glial cells by mass cytometry^46^, and identification of mRNAs for MBP and GFAP in P7 microglia by scRNA-seq^130^.

**Fig. 7.**
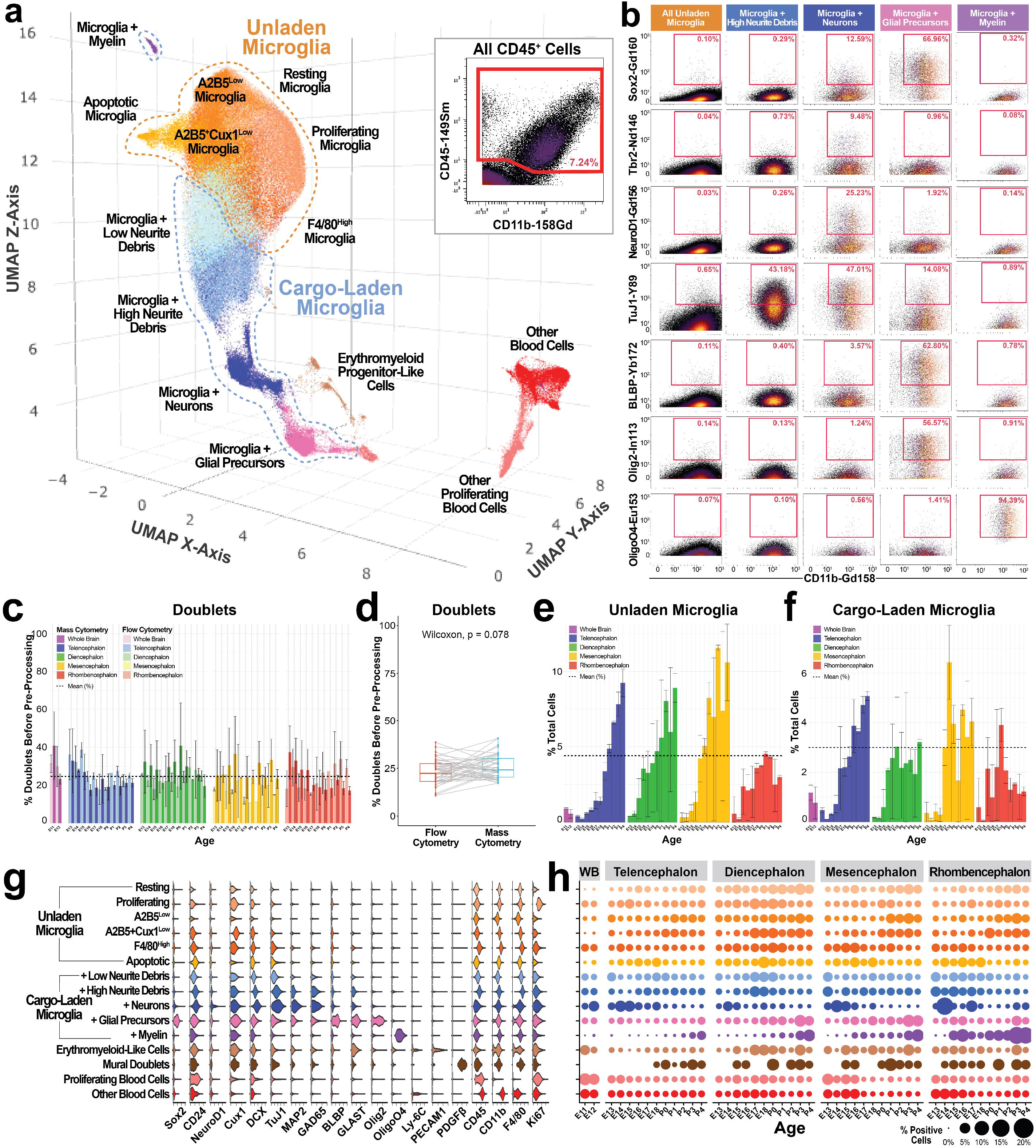
Identification of functionally distinct microglia subpopulations in the developing mouse brain. **a**, Three-dimensional UMAP of CD45+ cells colored according to Leiden cluster identity. Dashed lines indicate grouping of microglia into non-interacting (orange) and interacting (blue) clusters. Inset shows gating of CD45+cells using CytoBank. Cells inside the red box were included in analysis. **b,** Biaxial plots showing relative abundances of cells positive for select neural proteins. **c,** Relative percentages of doublets identified in each sample by flow cytometry and mass cytometry. Error bars indicate standard deviation. **d,** Wilcoxon test for significant difference in relative percentages of doublets identified for each sample by flow cytometry and mass cytometry. **e,f** Relative percentages of unladen (e) and cargo-laden **(f)** microglia identified by mass cytometry. **g,** Expression of select proteins key for distinguishing functional microglial subpopulations in the developing mouse brain. **h,** Dot plot showing relative abundances of CD45+ clusters in each brain region from El 1.5-P4. n = 2-5 litters per age/brain region for a total 1,758,016 cells from 112 samples.

Although we initially attempted to remove cargo-laden microglia from our analysis as artifactual cell doublets or aggregates (a bright-field image of cells directly prior to analysis on the mass cytometer is shown in **Extended Data Fig. 9b**), they were retained as cell singlets by our gating parameters (**Extended Data Fig. 2c–h**). To further investigate whether these observations represent single-cell events or cell aggregates, we quantified the extent of these behaviors in our tissue samples by two methods: flow cytometry with DAPI to identify cell events with greater than 4n DNA content (**Extended Data Fig. 9c**), and the 6-choose-3 doublet-filtering scheme used for mass cytometry barcoding (**Extended Data Fig. 9d**), where any cell event with greater than three palladium metals must contain cells from more than one barcoded sample^58^. As measured by both methods, the frequency of cell doublets or aggregates in samples (before pre-processing) did not exhibit any noticeable trend across age or tissues (**Fig. 7c,d**), in contrast to the observed frequencies of unladen and cargo-laden microglia (**Fig. 7e,f**). Unladen microglia generally increased in abundance with age in the forebrain and mesencephalon, although their numbers plateaued in the rhombencephalon (**Fig. 7e**). Cargo-laden microglia increased with age in the telencephalon, plateaued early in the diencephalon, displayed two waves centered around birth in the mesencephalon, and apexed before birth in the rhombencephalon (**Fig. 7f**). Notably, relative abundances of both unladen and cargo-laden microglia were lower in the rhombencephalon compared with the other brain regions. These differences in frequency from known doublets and aggregates, coupled with the established propensity of microglia for phagocytosis during brain development^92^, suggest these observations represent *bona fide* phagocytic events for microglia in the developing brain.

Microglia with positive but low DCX, TuJ1, and MAP2 levels indicate engulfment of neurite debris, while microglia positive for these neurofilaments plus neuronal-related transcription factors (e.g. Tbr2, NeuroD1, Tbr1, and/or Ctip2) indicate engulfment of entire neurons (**Fig. 7b,g; Extended Data Fig. 9e**). Similarly, microglia positive for glial and OPC markers indicate phagocytosis of these cell types, as previously reported^130, 133^. Microglia positive for OligoO4, but not other markers of OPCs/oligodendrocytes or neurites, likely represent myelin-engulfing microglia^134, 135^, which displayed caudal-to-rostral expansion mirroring myelin basic protein (MBP) expression in the developing mouse brain^93^.

Among unladen microglia, we observed Ki67^high^ proliferating microglia, cleaved-caspase 3^high^ apoptotic microglia, and several states without further defining characteristics other than varying expression levels of F4/80, A2B5, and Cux1. Although relative abundances of unladen microglia generally increased as brain development progressed, we observed more distinct trends for phagocytic microglia: a consistent increase after E15.5 in the telencephalon, a plateau after E17.5 in the diencephalon, two waves of expansion before and after birth in the mesencephalon, and one wave of expansion centered around birth in the rhombencephalon (**Fig. 7h**). While the antibody panel used in this study was not designed to probe microglial subtypes, the ability to quantify cargoes within these cells provides a novel method to monitor and characterize how microglia sculpt the developing brain with their phagocytic activity.

Concerning the origins of microglia in the brain, the prevailing theory is that microglia exclusively arise from yolk sac-derived primitive macrophages that migrate into the brain around E9.5^136, 137^, while endothelial lineages in the brain vasculature arise exclusively from angiogenic sprouting of endothelial cells in the perineural vascular plexus^124^. However, as mentioned above, we observed a rare population of CD45^low^PECAM^high^ cells, consistent with previous reports of cells resembling EMPs capable of producing erythroid, myeloid, and endothelial lineages in the embryonic mouse brain^91, 125, 126^. These cells generally increased in proportion from E11.5 to P4 (**Fig. 7h**) and mimicked the expansion of microglia, endothelial cells, and other nonneural lineages (**Fig. 2b**).

## DISCUSSION

In this study, we adapted mass cytometry for single-cell profiling of brain tissues to produce a protein-based cell atlas of the developing mouse telencephalon, diencephalon, mesencephalon, and rhombencephalon. Using sample replicates acquired daily across embryonic and postnatal development, we identified molecularly distinct cell populations, quantified their variability, and characterized their differentiation trajectories. As a resource, the companion dataset to this manuscript recapitulates decades of neural development research, characterizing the molecular profile and timing of appearance of virtually every major cell type, from progenitor cells, neurons, astrocytes, and oligodendrocytes, to microglia, vascular cells, and other nonneural cell types (**Fig. 8a**). In addition to recapitulating previous studies, our survey of the developing mouse brain provides novel insight into 1) the abundance and timing of rare neuronal progenitor and iGC populations, 2) the discrepancy between protein and mRNA levels during brain development, 3) the expression of Sox2 and nestin across a wide variety of cell types, including endothelial cells, 4) two distinct differentiation trajectories for generation of OPCs (direct-differentiating and iGC-derived), 5) the molecular dynamics underlying specification of glutamatergic neurons in the telencephalon, 6) CD45^low^PECAM^high^ EMP-like cells that may serve as an alternative source of microglia, and 7) cargo-laden microglia in the brain.

**Fig. 8.**
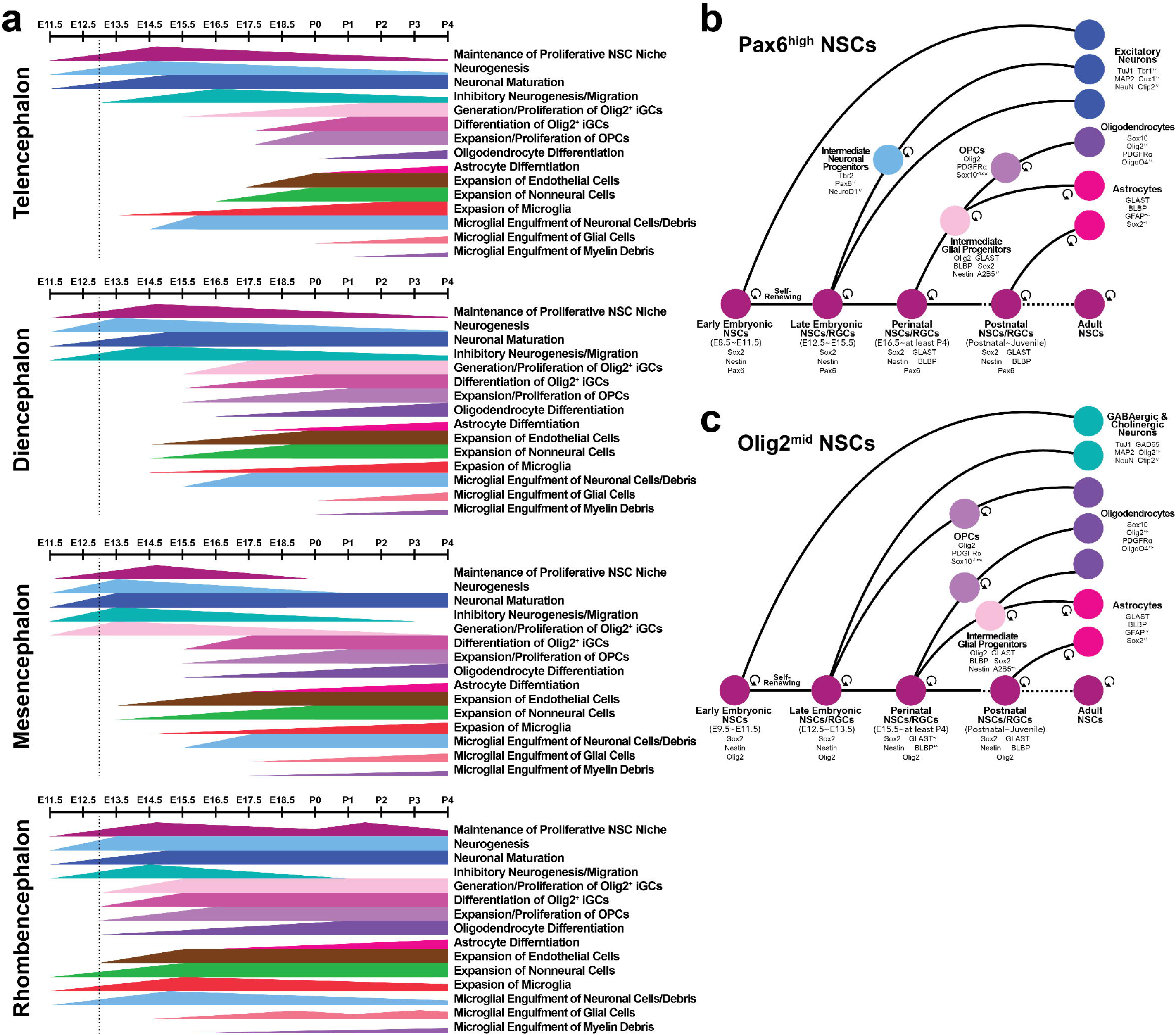
Overview of key processes in mouse brain development. **a**, Schematic of major processes involved in development of the mouse telencephalon, diencephalon, mesencephalon, and rhombencephalon from E11.5-P4. **b** and **c,** Overview of how the differentiation potential of Pax6high NSCs **(b)** and Olig2mid NSCs (**c)** changes over the course of mouse brain development.

Consistent with previous reports^89, 123, 138, 139^, our analyses support a model of neural cell differentiation in which RGCs primarily generate neurons and INPs until E15.5/16.5, increasingly produce OPCs and iGCs perinatally, and primarily generate glial cells or become quiescent postnatally (**Fig. 8b**). However, our findings also emphasize the dynamic and mosaic nature of neural stem cell niches across different regions of the developing brain. For example, we observed cells committed to direct and indirect OPC trajectories from E11.5 to P4 (**Fig. 5**), suggesting that both of these differentiation pathways can occur during all three waves of OPC generation^89^. We hypothesize that the Olig2^mid^ NSC populations initially responsible for generating specific neuronal subtypes (e.g. cholinergic and GABAergic neurons in the ventral forebrain^81, 140, 141^, and Purkinje cells in the cerebellum^69^) are primed to subsequently generate OPCs by a molecular pathway that does not involve a bipotent BLBP^high^GLAST^high^ RGC/iGC intermediate capable of generating astrocytes and OPCs (**Fig. 8c**). This notion is supported by the presence of Olig2-expressing NSCs in the ventricular zone of the medial ganglionic eminence (MGE) as early as E9.5^82^ and cerebellum at E10.5^142^, Olig2^+^PDGFRɑ^+^ OPCs in the subventricular zone of the MGE^143^, and the sparse number of oligodendrocytes identified in a *Glast* lineage-tracing study^144^. Moreover, a recent scRNA-seq study identified iGC-dependent and -independent differentiation pathways for OPCs in the neonatal mouse cortex and human glioma^86^.

The ability to measure millions of cells per hour with mass cytometry enables quantification of rare cell populations. Here, we observed several infrequent cell populations, such as Pax6^high^GAD65^high^ neuronal progenitors, Ctip2^high^NeuroD1^high^ neuronal progenitors, and PDGFRɑ^neg^ OPCs. But perhaps the most notable rare cell population identified was a small population of CD45^low^PECAM^high^ EMP-like cells. Previous reports identifying cells co-expressing CD45 and endothelial markers as early as E9.5 in the mouse head (prior to embryonic circulation) propose a local and autonomous source of hemogenic endothelial cells (i.e. hematopoietic stem cells)^125, 126, 145^, in contrast to prevailing theories of endothelial cell sprouting^124^ and migration of primitive macrophages from the yolk sac into the brain^136, 137^. Accordingly, the identities and molecular trajectories of progenitor cells involved in cephalic vasculogenesis and angiogenesis remain unclear, and there exists the potential for a *de novo* source of hematopoietic and endothelial cells in the embryonic brain. Beyond supporting the existence of EMP-like cells in embryonic mouse brain as early as E11.5, our results show that CD45^high^PECAM^high^ cells persist thereafter into the perinatal period at relative abundances that mimicked expansion of endothelial and hematopoietic cells in each brain region. These results are reminiscent of recent findings showing that a lymphatic-derived microglial population seeds the zebrafish brain prior to colonization by yolk sac-derived microglia and persists thereafter until at least the juvenile stage^146^. Accordingly, the extent to which this putative stem cell pool contributes to erythroid, myeloid, and/or endothelial progeny during mouse brain development is a promising area of inquiry for future studies.

Reminiscent of our previous observation of putative phagocytosis by satellite glial precursors in the dorsal root ganglia^46^, a fortuitous result of this study was the observation of putative phagocytic cargoes in microglia. Phagocytosis by microglia plays an important role in brain development^134, 147, 148^, homeostasis^130^, injury^149^, and disease^150^. Notably, our findings indicating phagocytosis of myelin by microglia at embryonic ages represent the earliest description of this developmental phenomenon, corroborating recent findings in zebrafish, P10 mouse optic nerve, and P14 mouse brain^134, 135^. Clean-up gating and doublet analysis indicate that it is unlikely the putative phagocytic cargoes observed here are due to cell doublets or larger aggregates. One remaining caveat is the possibility that these observations are due to neuronal and glial cell fragments sticking to the outside of microglia. Performing confocal immunofluorescence microscopy to observe the complete engulfment of neuronal and glial cargoes would provide further evidence that we are observing *bona fide* phagocytic events. If these putative phagocytic events are corroborated, future studies could provide a more detailed view of microglial subtypes and their phagocytic behavior by employing a hybrid antibody panel including additional markers to define microglial subpopulations^38, 39, 43–45, 47, 130, 151^, as well as markers of phagocytic activity and cargoes. With additional validation and characterization, this approach to quantify phagocytic cargoes could be extended beyond microglia to macrophages and “non-professional” phagocytes like neural crest cells^152^.

As mentioned above for mural cells and putative phagocytosing microglia, the loss of cell processes (e.g. neuronal axons and dendrites, astrocytic end feet, and myelin sheaths) during single-cell dissociation is an important caveat for analyses of neural tissues by mass cytometry and scRNA-seq. In particular, single-cell dissociation hinders analysis of axonal and dendritic proteins in neurons, although many of these are synthesized in the cell body and can be detected there at lower levels. Postsynaptic proteins are less prone than presynaptic proteins to marker loss in this manner, because they can be localized in synapses on the cell body as well as in dendrites. Cell dissociation and sample processing may also cause fragments of cells or sheared processes to stick to the surface of other cells, resulting in cells appearing positive for markers of two discrepant cell types – as appears to be the case for the mural cells observed.

Another caveat for single-cell analysis of neural tissues is the potential for systematic bias in the quantification of relative cell-type abundances, which could be caused by depletion of specific cell populations during sample processing. While over 85% of the cell events we measured were viable (**Supplementary Table 1**), it is possible that particularly fragile cells lyse during sample dissociation and are therefore lost from analysis. As an example, the relatively low numbers of cleaved-caspase 3^+^ cells in our study may reflect the fragility of cells undergoing apoptosis, rather than their low abundance in the developing brain. Another potential source of systematic cell-type depletion is that cells resistant to dissociation will be filtered out as clumps or cell aggregates, either physically by the 40-micron strainer, or computationally by the doublet-filtering barcode scheme (**Methods**). As above, immunofluorescence microscopy can be applied on tissue slices to corroborate cell abundances measured by mass cytometry, although single slices may not be representative of the entire tissue. For tissue imaging with enhanced capacity for molecular profiling, the metal isotope-labeled antibodies used in this study can also be applied for mass spectrometry-based imaging, either with imaging mass cytometry^153^ or multiplexed ion beam imaging^154^.

While the cell populations we identified by protein-based measurements show general agreement with those previously identified by scRNA-seq^9–11, 13–15, 16, 17–24, 25, 49, 50, 51–55^, the magnitude of discrepancies between specific protein-mRNA pairs was unexpected, and demonstrates the value of protein measurements to characterize and quantify functional cell states. The wide variety of relationships we observed between mRNA and protein abundance likely arises from a combination of many factors, such as varying rates of mRNA processing and degradation, as well as varying rates of protein translation, maturation, trafficking, and degradation, all of which can differ between specific protein-mRNA pairs, across cell types, and across developmental stages within a single cell type. For example, translational control, protein degradation, and incomplete trafficking or internalization of surface proteins may result in high levels of mRNA but no protein present. Conversely, mRNA degradation may result in high levels of long-lived proteins, but no mRNA present. Investigating these mechanisms is beyond the scope of this study, but could be investigated further by split sample measurements with mass cytometry and scRNA-seq, or with CITE-seq^155^ to simultaneously detect protein and RNA abundances in single cells.

The neural mass cytometry approach developed in this manuscript will enable future mechanistic studies on mouse neurological disease models, enhance characterization of genetic and pharmacological perturbations of brain development, and can also be used to study *in vitro* models of neural differentiation employing embryonic or induced pluripotent stem cells. The high sample throughput and relatively low cost of mass cytometry make it particularly attractive for larger scale studies, where increased sample numbers allow for statistical comparisons between conditions, and increased cell numbers allow for the characterization of rare, low-frequency cell populations. Indeed, the breadth of findings presented in this manuscript are derived from a single experiment run over 3 days with barcode-multiplexed samples, demonstrating the power of mass cytometry as a high-throughput platform for single-cell analyses. Potential for further optimization includes adapting the antibody panel to measure human brain cells; at present, 27 out of 40 antibodies in our mouse CNS staining panel are cross-reactive for their respective human homologs (**Extended Data Table 1**). The antibody panel could also be modified to focus on specific brain regions, and tissue dissociation methods could be optimized for analysis of adult tissues. Collectively, our mass cytometry analyses represent a region-specific roadmap of cell specification and maturation in the mouse brain, and provide a high-throughput, multiplexed platform to investigate fundamental mechanisms of development at the protein level.

## METHODS

### Animals

All animal husbandry and experiments were carried out in accordance with guidelines of the Association for Assessment of Laboratory Animal Care and approved by the University of Virginia Animal Care and Use Committee (Deppmann Protocol No. 3795). Mice were harvested from C57/BL6 females (Jackson Labs, 000664) bred in house from embryonic day 11.5 (E11.5) to postnatal day 4 (P4). For timed pregnancies, animals were mated overnight and removed the following day once a week. Animals were housed with a 12-h light/dark cycle with food and water *ad libitum*.

### Dissection and single-cell dissociation of brain tissue

Following decapitation of embryonic and postnatal mice, dissociation of viable single cells from brain tissue was optimized similar to a previously reported strategy^156^. Whole brains were removed from mice aged E11.5 to P4 and placed in 35-mm Petri dishes containing Dulbecco’s phosphate-buffered saline (PBS; Thermo Fisher Scientific, 14190) on ice. After dissection of whole brains (E11.5/E12/5) or microdissection into cortex, diencephalon, midbrain, and cerebellum/hindbrain (E13.5–P4) (**Fig. 1a; Extended Data Fig. 5a**), tissues were separated, meninges were removed, and samples from whole litters were pooled. A P1000 micropipette was used to add 1 mL of Dulbecco’s Modified Eagle’s Medium containing 4.6 mg/mL dispase II (Sigma-Aldrich, D4693), 1 mg/mL collagenase type IV (Worthington, LS004186), 0.2 mg/mL DNAse-I (Sigma-Aldrich, 11284932001), and 0.2 mg/mL hyaluronidase (Sigma-Aldrich, H3884) to each dish. Tissue was immediately mechanically dissociated by mincing with forceps and gently pipetting up and down five times with a P1000 micropipette, before transferring the resulting cell slurry to a microcentrifuge tube. After incubation at 37°C in a water bath for 20 min, the cell suspension was passed through a 75-µm sieve and 45-µm sieve (Thermo Fisher Scientific, 50871316 and 50871319) with a P1000 micropipette. Tubes were then centrifuged at 300 × g for 3 min at 4°C, the supernatant was discarded, and cells were washed by adding 1.5 mL of PBS containing 0.5% bovine serum albumin (BSA; Sigma-Aldrich, A9418) and gently pipetting. After centrifugation at 300 × g for 3 min at 4°C, cells were resuspended in 100 µL of PBS.

### Cisplatin staining and fixation of single-cell suspensions

Resuspended cells were mixed with 100 µL of 2× cisplatin solution (10 µM in PBS; Sigma Aldrich, P4394) with a P1000 micropipette, incubated at room temperature for 30 sec, and then quenched with 1.3 mL of PBS containing 0.5% BSA. Following centrifugation at 300 × g for 3 min at 4°C, supernatants were removed, and the resulting cell pellets were washed once with PBS containing 0.5% BSA. Subsequently, cells were fixed for 10 min at room temperature in 1 mL of 1.6% paraformaldehyde solution (Electron Microscopy Services, CAS 30525-89-4) in PBS. Following fixation, the cell suspension was centrifuged at 600 × g for 3 min at 4°C, washed once with PBS, centrifuged again, and resuspended in 1 mL of cell staining medium (CSM; 0.5% BSA, 0.02% NaN3 in PBS). A 100-µL aliquot of each sample was placed in a separate tube (for flow cytometry analysis) before storing samples and aliquots at -80°C until use for analysis.

### Cell counts and visual inspection by light microscopy

To confirm the quality of dissociation, fixed cells were visually inspected in bright field mode at 4X, 10X, and 20X using an EVOS AMF4300 microscope (Thermo Fisher Scientific, Waltham, MA). Samples exhibiting a preponderance of single cells with low levels of debris and cell clumps were counted using a Bio-Rad TC20 Automated Cell Counter (Hercules, CA).

### Flow cytometry

To validate the quality of dissociated cells, samples that passed visual inspection were subjected to flow cytometry with the fluorescent nuclear stain DRAQ7^TM^ (Biolegend, San Diego, CA) and an Alexa Fluor^TM^ 488-conjugated antibody against the neuronal intermediate filament beta-tubulin III (TuJ1). Briefly, aliquots of each cell sample were thawed on ice, pelleted by centrifugation at 600 × g for 3 min at 4°C, and the supernatant was discarded. Next, cells were permeabilized with ice-cold 100% methanol and incubated on ice for 10 min with vortexing every 2 min. After centrifugation at 600 × g for 3 min at 4°C, the supernatant was discarded and cells were washed once with CSM. Cells were incubated with TuJ1 antibody (1:1000 in CSM; Biolegend) for 1 h at room temperature on a shaker. After incubation, cells were centrifuged at 600 × g for 3 min at 4°C, the supernatant was discarded, and DRAQ7 (1:5000 in PBS) was added for 5 min. Samples were immediately measured on an Attune^TM^ NxT flow cytometer (Thermo Fisher Scientific) and analyzed using CytoBank (community.cytobank.org).

### Metal conjugation of antibodies

Purified antibodies (lyophilized or in buffer free of BSA and gelatin) were conjugated to isotopically pure metals (listed in Table 1) for mass cytometry analysis using Maxpar® antibody conjugation kits (Fluidigm) according to the manufacturer’s instructions. Immediately following conjugation, stock solutions were prepared for long-term storage at 4°C by diluting conjugated antibodies at least two-fold with Candor PBS Antibody Stabilization Solution (Candor Bioscience GmbH). Final concentrations of antibodies in stock solutions ranged from 0.05–0.4 mg/mL.

### Validation of antibodies

Following metal conjugation, each antibody was titrated to determine the optimal concentration for mass cytometry analysis. To define the antibody concentration providing the highest signal-to-noise ratio, we employed positive and negative counterstains (e.g., TuJ1 for neurons, BLBP for glial cells, CD45 for hematopoietic cells) to evaluate cell samples from mouse brain, a mouse embryonic stem cell line (E14Tg2a, ATCC, CRL-1821), two mouse neuroblastoma cell lines [N1E-115 (ATCC, CRL-2263) and Neuro-2a (ATCC, CCl-131)], one mouse glioma cell line (GL261, National Cancer Institute Division of Cancer Treatment and Diagnosis Tumor Repository, Glioma 261), two mouse OPC cell lines (OPC-1052 and OPC-8173, gift from Prof. Hui Zong, University of Virginia), and a human embryonic kidney cell line (293T, ATCC, CRL-3216). The optimal concentration of each metal-conjugated antibody preparation was defined as the concentration providing the highest signal-to-noise ratio between appropriate positive and negative controls (**Extended Data** **Fig. 1a**). Antibodies were considered specific and reliable if they produced signal in DNA intercalator-positive cells exhibiting one or more positive counterstains, and were absent in cells exhibiting a negative counterstain (**Extended Data** **Fig. 1b,c**).

### Sample barcoding, staining, and intercalation for mass cytometry

To prepare samples for mass cytometry, frozen cells were thawed and pelleted by centrifugation at 600 × g for 3 min at 4°C. After removing the supernatant, cells were washed once with CSM and resuspended in 0.5 mL of cold saponin solution (0.02% in PBS) containing one of the twenty 6-choose-3 combinations of 1 mM isothiocyanobenzyl EDTA-chelated palladium metals, as previously described^58, 60^. Following incubation on a shaker at 800 rpm for 15 min at room temperature, samples were centrifuged at 600 × g for 3 min at 4°C and the supernatant was discarded. The resulting cell pellet was washed three times with CSM and then pooled with other samples into a total of seven barcoded sets for antibody staining (**Supplementary Table 1**).

Prior to staining of surface epitopes, each barcoded set was blocked in CSM containing 10% (v/v) normal donkey serum (Millipore, S30-100ML) for 30 min at room temperature. After blocking, antibodies indicated as “Surface” in **Extended Data Table 1** were diluted in CSM and added to cells (100 uL staining volume per 1 × 10^6^ cells). Following incubation at room temperature on a shaker at 800 rpm for 30 min, samples were centrifuged at 600 × g for 3 minutes at 4°C and the supernatant was discarded. After washing the cell pellet three times with CSM, cells were permeabilized for intracellular staining by filling the sample tube with ice-cold 100% methanol and incubating on ice for 10 min with vortexing every 2 min. Next, samples were centrifuged at 600 × g for 3 min at 4°C, the supernatant was discarded, and cells were washed once with CSM. Samples were then incubated with primary antibodies listed as “Intracellular” in **Extended Data Table 1** (diluted in CSM) for 1 h at room temperature on a shaker at 800 rpm. After incubation, samples were centrifuged at 600 × g for 3 min at 4°C, the supernatant was discarded, and cells were washed three times with CSM.

After primary antibody staining, cells were stained with 0.1 uM Cell-ID^TM^ Intercalator-Ir (201192, Fluidigm) in 1.6% PFA containing for 15 min overnight at 4°C. After intercalation, cells were washed once with CSM, once with water, once with 0.05% Tween-20 (in water), and again with water. Finally, cells were pelleted by centrifugation at 600 × g for 3 min at 4°C and then samples were kept on ice until run on the mass cytometer.

### Mass cytometry

Immediately before analysis, cells were resuspended in Maxpar® Cell Acquisition Solution (approximately 1 mL per 1 × 10^6^ cells, Fluidigm) containing 1:20 EQ^TM^ Four Element Calibration Beads (Fluidigm) and pipetted through a 40-µm nylon mesh filter. Cells were analyzed in multiple runs on a Helios^TM^ CyTOF® 2 System (Fluidigm Corporation) at a rate of 500 cells per second or less.

### Normalization and debarcoding

To control for variations in instrument signal sensitivity across individual mass cytometry runs, raw .fcs data files were normalized using EQ Four Element Calibration Beads as described in Finck et al. 2013 (https://github.com/nolanlab/bead-normalization) (**Extended Data Fig. 2a**). The resulting normalized .fcs files from each run were concatenated for each sample set and then debarcoded using software described in Fread et al., 2017 (https://github.com/zunderlab/single-cell-debarcoder) to deconvolute palladium metal expression on single cells, thus permitting identification of individual samples according to a 6-choose-3 combinatorial system^58^ (**Extended Data** **Fig. 2b**). The modified version adds a new parameter for barcode negativity (bc_neg) that is the sum of the counts for the three barcode metals expected to equal zero based on the barcode deconvolution assignment. High values of this parameter indicate events likely containing two or more cells.

### Isolation of single-cell events

To isolate single cells from debris and clumps of multiple cells, .fcs files processed as described above were uploaded to CytoBank (community.cytobank.org) and gated according to the strategy illustrated in **Extended Data** **Figure 2c–h**. First, an additional debarcoding process was performed by gating out events with a low barcode separation distance, high Mahalanobis distance, and/or high signal for non-barcode metals (**Extended Data Fig. 2c**). Subsequently, singlets were isolated by comparing the center of events with their length and width (**Extended Data Fig. 2d**).

To remove non-viable cells, events with high Pt195 signal (indicating high cisplatin uptake before fixation) were removed (**Extended Data Fig. 2e**). Next, events with a high cerium (Ce140) signal, indicating potential failure to remove a calibration bead during normalization, were removed (**Extended Data Fig. 2f**). Finally, gating was applied to remove events exhibiting high spectral overlap between metal isotopes (**Extended Data Fig. 2g**) and cellular debris (**Extended Data Fig. 2h**). Isolated single-cell events for individual timepoints were exported to .fcs files for further analysis. Raw.fcs files for mass cytometry data presented within this manuscript accessible at Flow Repository (https://flowrepository.org/) [ID: FR-FCM-Z5LD]. In addition, FCS files for samples and clean-up gating used for sample pre-processing are available on CytoBank at https://community.cytobank.org/cytobank/experiments/ (Experiment IDs: 105280–105284).

### Batch correction

To account for changes in antibody signals between barcode sets and remove the potentially negative influence of aberrant signals from debris or cell clumps on normalization of signals from individual markers, batch correction was performed after isolating single-cell events as described above. To correct for differences in signal intensities of individual markers across the seven barcoded sets (batch effects), debarcoded .fcs files were processed as described in Schuyler et al. 2019 (https://github.com/CUHIMSR/CytofBatchAdjust). Briefly, each barcoded set contained a universal sample (mixture of all ages and brain regions examined). Antibodies that produced Gaussian distributions and mean signals with variance greater than 1% for this universal sample were corrected at the 50^th^ percentile, including: BLBP, CD24, Cux1, GAD65, GLAST, MAP2, N-cadherin, nestin, NeuN, NeuroD1, p75NTR, Pax6, PSA-NCAM, Sox2, Sox10, and TuJ1 (**Extended Data Fig. 2i**). Mean signals for ALDH1A1, CD11b, and SSEA-1 had 2%–3% variance and normal distributions with truncated lower tails. Batch correction of these markers at 80^th^ and 95^th^ percentiles was determined to be ineffective at reducing the variance of mean signal, while correction at the 50^th^ percentile resulted in overcorrection (ALDH1A1 and SSEA-1) or a non-zero mean (CD11b); therefore, these markers were not batch corrected. The remaining markers were not batch corrected because the variance of their mean signal was less than 1%.

### ASinH Scaling

To minimize background signal levels and provide the greatest signal-to-noise ratio, ArcSinH values were manually scaled for each antibody using CytoBank. Default (ArcSinH = 5) and final ArcSinH transformation values are shown in **Extended Data Fig. 2j**. The default value was used for markers not shown.

### Clustering of high-dimensional data

To reduce the effect of differences in tissue and sample size on clustering, 5 × 10^5^ cells were randomly selected from each age/region for the global analyses shown in **Figures 1–3**, yielding a total of 5.75 × 10^6^ (∼24%) cells from the original dataset (2.43 × 10^7^ cells). For analyses shown in **Figures 4 and 5**, clustering was performed on all Sox2^+^Nestin^+^ cells (defined by the manual gating strategy in **Fig. 4a**), yielding a total of 3,253,641 cells (∼13%) from the original dataset. For analyses of the telencephalon shown in **Fig. 6**, clustering was performed on 1.25 × 10^5^ cells randomly sampled from each replicate (n = 2 per age), yielding a total of 3.25 × 10^6^ cells (∼47%) from the 6,855,672 telencephalon cells in the original dataset. For analyses shown in **Fig. 7**, clustering was performed on all CD45+ cells (defined by the manual gating strategy shown in the inset in **Fig. 7a** and **Extended Data Fig. 9a**), yielding a total of 1,345,841 cells (∼5.5%) of the original dataset.

To identify discrete cell populations within the developing mouse brain, isolated single cells were subjected to two rounds of high-dimensional analysis. For both rounds of clustering, all markers were included for generation of UMAP layouts to distinguish major classes (i.e. NSCs, neuronal progenitors, glial progenitors, neurons, astrocytes, oligodendrocyte progenitors, hematopoietic cells, and mural cells).

### Assignment of cluster identities

The cell-type specificity of each antibody used in the panel is outlined in **Extended Data Table 1**. Accordingly, cells were organized into major classes by their molecular profile as follows: 1. <u>Neural stem cells</u> (NSCs): positive for Sox2 and nestin with or without Pax6, Sox1, Olig2, CD133, and CD24; negative for mature neural markers. 2. <u>Intermediate neuronal progenitors</u>: positive for Tbr2. 3. <u>Neurons</u>: positive for doublecortin (DCX), β-tubulin 3 (TuJ1), and/or microtubule-associated protein 2 (MAP2), with low levels of the postmitotic neuronal marker NeuN. 4. <u>Inhibitory neurons</u>: positive for glutamic acid decarboxylase (GAD65). 5. <u>Radial glial cells/glial</u> <u>precursors</u>: high expression glutamate aspartate transporter 1 (GLAST) and brain lipid-binding protein 7 (BLBP), with or without glial fibrillary acidic protein (GFAP). 6. <u>OPCs</u>: variable levels of Olig2, platelet-derived growth factor alpha (PDGFRɑ), and Sox10; two clusters also expressed low levels of oligodendrocyte marker O4 (OligoO4), indicating differentiation into early oligodendrocytes. 7. <u>Neuronal progenitors</u>: combined expression of markers associated with NSCs and mature neurons, negative for glial markers. 8. <u>Glial progenitors</u>: combined expression of markers associated with NSCs and RGCs/astroglial cells, with or without Olig2; negative for neuronal markers. 9. <u>Endothelial cells</u>: positive for platelet endothelial cell adhesion molecule-1

(PECAM1) with or without Ly6C. 10. <u>Mural cells</u>: positive for melanoma cell adhesion molecule (MCAM) or PDGFR-ꞵ; notably, mural cell clusters also contain several neural markers, but do not have elevated DNA-intercalator levels, indicating that these are not cell doublets or aggregates. Instead, the most likely explanation is that neural cell debris is sticking to their cell surfaces. 11. <u>Non-neural cells</u>: negative for virtually all neural cell markers, although the presence of markers such as PDGFR⍺, P75 neurotrophic receptor (P75NTR), Cux1, CD24, and VCAM indicated the presence of putative fibroblasts among these clusters. 12. <u>Microglia</u>: low expression of canonical markers CD45 and CD11b, as well as positive expression of F4/80. 13. <u>Other hematopoietic cells</u>: high expression of CD45 and CD11b. 14. <u>Neural crest-derived cells</u>: high expression of Sox10 and negative for Olig2. Two putative non-brain populations appeared to contaminate early embryonic dissections, a cluster exhibiting low expression of P75NTR, BLBP, and some stem cell markers presumably representing developing cranial ganglia; and a cluster exhibiting ALDH1A1 expression presumably representing a population of developing glial cells. 15. “<u>Apoptotic” cells</u>: high expression of cleaved caspase 3; notably, cells with low to moderate expression were observed in numerous clusters (especially glial progenitors). 16. <u>Low-complexity cells:</u> one small cluster of cells was negative for expression of all panel proteins, representing 2.16% of total cells analyzed.

### Visualization of high-dimensional data with UMAP

Data was visualized by uniform manifold approximation and projection (UMAP, https://github.com/lmcinnes/umap)^63^ using the following parameters: nearest neighbors = 15, metric = euclidean, local connectivity = 1, N components layout = 2 (3 for UMAP in **Fig. 7a**), N components cluster = 2, N epochs = 1000.

### Leiden clustering

Community detection with the Leiden algorithm^62^ was performed to partition cells into ‘clusters’ according to molecular profile similarity using Python. To improve computational speed and scalability, the hnswlib package (https://github.com/nmslib/hnswlib) was incorporated into this process (https://github.com/zunderlab/VanDeusen-et-al.-CNS-Development-Manuscript/blob/main/02_UMAP_and_Leiden_Clustering/03_Leiden.py) using the following parameters:

Hierarchical Navigable Small World Graphs (HNSWG) space = 12, HNSWG EFConstruction = 200, HNSWG M = 16, HNSWG EFSet = 20.

### Comparison of protein and RNA expression profiles

To compare protein and RNA expression levels, we chose two of the most comprehensive data sets available, *in situ* hybridization data from the Allen Brain Atlas^93^ and the largest scRNA-seq data set published to data^53^. Microdissected regions were aligned between all three datasets (**Extended Data Fig. 5a,b**). ‘Structure Unionized’ data was downloaded from the Allen Brain Atlas for each aligned brain region for the 25 available genes corresponding to markers in our mass cytometry antibody panel, and the values for “sum_expressing_pixel_intensity”, “sum_expressing_pixels”, and “sum_pixels” were extracted and used to calculate the percentage of cells expressing a gene and the mean expression level in those expressing cells. The expression counts matrix for the scRNA-seq data was downloaded from mousebrain.org and preprocessed according to the original publication^53^. To calculate percentages of expressing cells for scRNA-seq and mass cytometry data, cells in scRNA-seq data were considered ‘expressing’ if their value was above zero. For mass cytometry data, which typically exhibits low background for individual markers compared with any antibody-based technique, thresholds for labeling cells as ‘expressing’ were manually assessed as the 99^th^ percentile of expression in low-complexity cells (**Extended Data Fig. 5c**). Mean expression was then calculated for the population of identified expressing cells for each gene/marker per each brain region. For all three datasets, the calculated mean expression values were finally per-feature range-normalized to fall between 0 and 1. Percentages of cells expressing each marker and normalized mean expression values were then examined for each RNA-protein pair in each brain region and plotted using R. These summarized scRNA-seq and mass cytometry data were then used to perform dynamic time warp (DTW) analysis using the dtw package (v1.22-3) in R to examine the timing alignment of relative expression changes between the two datasets^94^. All time points for both datasets were used in DTW analysis. To quantify the lead/lag between these two time series, cross-correlation analysis was performed using the base R ccf function. To generate the input data for this analysis, matching time points were extracted from the summarized scRNA-seq and mass cytometry datasets and differencing was applied for each dataset across these matched timepoints.

### Identification of developmental cell trajectories with URD

To examine fate decisions involved in developmental cell trajectories, the continuous diffusion-approximation R package URD^56^ was adapted for use with mass cytometry data. “Root” and “tip” cells were manually chosen as the beginning and end points for construction of a map based on diffusion of protein expression levels (**Fig. 5a** and **Fig. 6b**). All markers were used for analysis with the following URD parameters: floodPseudotime n= 500, minimum.cells.flooded = 2, max.frac.NA = 40, knn = 15. Because of computational limitations, both the Sox2^+^Nestin^+^ cell dataset and telencephalon dataset were randomly and proportionally (with regard to sample age and cluster, respectively) sampled to a total 61,000 cells. The URD algorithm assigned each cell a pseudotime value based on its distance from the root, indicating its relative position along the differentiation trajectory from root to tip. By identifying intersecting paths from thousands of random walks from each tip back to the root, URD constructs a dendrogram from which molecular trajectories and branchpoints can be derived. Pax6^+^ NSCs present at E11.5 were chosen as the root for both URD analyses, while clusters chosen as tips were selected because their molecular profile indicated a mature differentiation status (i.e. relatively high expression of mature neural markers and low expression of stem/immature markers compared with subclusters of similar identity). To ensure proper representation of neurogenic Sox2^+^Nestin^+^ subpopulations that became sparse postnatally, cells in tip subclusters from P0–P4 samples were included in this analysis and selected irrespective of their origin in the brain

### Quantification of cell doublets/aggregates by flow cytometry and mass cytometry

As described above in the Flow Cytometry section, cells were stained with DRAQ7 (a DNA intercalator) and TuJ1 conjugated to Alexa Fluor 488. Positive cells exhibiting increased ratios of forward- or side-scatter area to width were considered doublets/cell aggregates, as verified by increased DRAQ7 positivity. To approximate this approach using mass cytometry, proportions of cells exhibiting high positivity for non-barcode-specific palladium isotopes, low barcode separation distance (indicating cell aggregates/doublets occurring because of immunocytochemical processing), and/or off-center signals (indicating abnormally dense metal ion contents of potential doublets/aggregates occurring in the original “single-cell” suspension) were quantified.

To test the robustness of our method to identify (and exclude) cell doublets and aggregates by mass cytometry, differences between relative abundances of doublets quantified by flow and mass cytometry were compared for ages and brain regions using Student’s t-test. Of the 42 ages/brain regions analyzed (four were excluded because of a lack of paired samples), none exhibited a significant difference (p > 0.09) in proportions of doublets identified by mass or flow cytometry. Moreover, Wilcoxon test of all paired samples revealed no significant difference between proportions of doublets/cell aggregates identified by flow cytometry and mass cytometry (p = 0.078, **Fig. 7h**).

## Supporting information

Extended Data Figures Legends and Bibliography

Souce Data Figure 1

## Author Contributions

A.L.V., C.D.D., and E.R.Z. planned all experiments. A.L.V., I.C. and A.B.K. collected tissue and performed single-cell dissociations. A.L.V. performed all antibody conjugations. A.L.V. validated all antibodies by performing titration experiments on positive and negative control cell samples. A.L.V. and E.R.Z. performed the mass cytometry measurements. A.L.V., S.M.G., C.M.W, A.B.K., K.I.F., and E.R.Z. wrote scripts for data analysis. A.L.V., S.M.G., and E.R.Z. performed data analysis. A.J.S. contributed protein chemistry expertise and resources. E.R.Z. and C.D.D. conceived and supervised all aspects of the project. A.L.V., S.M.G. and E.R.Z. prepared figures. A.L.V., E.R.Z., and C.D.D. wrote the manuscript, with input from all authors.

## Acknowledgments

Research reported in this publication was supported by the National Institute of Neurological Disorders and Stroke of the National Institutes of Health under award number R01NS111220 to E.R.Z. and C.D.D. The content is solely the responsibility of the authors and does not necessarily represent the official views of the National Institutes of Health. Further support was provided by the 3 Cavaliers Pilot research program to C.D.D. and E.R.Z. We thank Hui Zong for providing cell lines to validate antibodies, and for providing feedback on the manuscript. We thank Sarah Kucenas for providing the Sox10 antibody and feedback on the manuscript. We thank Thomas Müller and Carmen Birchmeier for providing the BLBP antibody. We thank Yipkin Calhan, Barry Condron, Ali Güler, Micah Hunter-Chang, Sushanth Kumar, Amrita Pathak, Jonathan Sewell, and Ekaterina Stepanova for providing feedback on the manuscript. We thank Ashley Hirt for assistance with coding and data analysis, Katrina Warner for assistance with antibody validation, and Lucy Jin for assistance with mouse dissections. We thank the University of Virginia Flow Cytometry Core, RRID: SCR_017829 for technical assistance with the CyTOF mass cytometer instrument. The authors acknowledge Research Computing at The University of Virginia for providing computational resources and technical support that have contributed to the results reported within this publication. URL: https://rc.virginia.edu.

## Declaration of Interests

The authors declare no competing interests.

## Data availability

Requests for reagents and other resources will be fulfilled by Eli Zunder. Raw and processed single-cell mass cytometry datasets will be deposited at Flow Repository (https://flowrepository.org/) upon publication. FCS files for each individual sample (debarcoded, normalized, and batch corrected) and clean-up gating used for sample pre-processing are available on CytoBank at https://community.cytobank.org/cytobank/experiments/ [Experiment IDs: 105280–105284].

## Code availability

Code used to perform analysis of mass cytometry data (as detailed in the Methods section) was adapted from standard R and Python packages, including UMAP, LeidenAlg, Seurat, dtw, URD, and ggplot, and is available on GitHub at https://github.com/zunderlab/VanDeusen-et-al.-CNS-Development-Manuscript. More detailed information is available upon request.

## Contact for reagent and resource sharing

Resources, reagents, and further information will be provided by Eli Zunder upon reasonable request.

## Notes

https://flowrepository.org/id/FR-FCM-Z5LD

https://community.cytobank.org/cytobank/experiments/105280/

https://community.cytobank.org/cytobank/experiments/105281/

https://community.cytobank.org/cytobank/experiments/105282/

https://community.cytobank.org/cytobank/experiments/105283/

https://community.cytobank.org/cytobank/experiments/105284/

### Competing Interest Statement

The authors have declared no competing interest.

