## Extended Data Table 1 for "A developmental atlas of the mouse brain by single-cell mass cytometry"

**Extended Data Table 1.** Antibodies used for mass cytometry.

| Metal | Antibody/Reagent | Full Name(s) | Cell Types in the Brain | Original Identification for Cell-Type Specificity | Vendor | Catalog No. | Host | Clone | [CyTOF] | Staining Protocol | Reactivity |
|---|---|---|---|---|---|---|---|---|---|---|---|
| Y89 | TuJ1 | Beta 3-tubulin | Neurons | Caccamo, 1989 | Covance | MMS-435P-250 | Mouse | TuJ1 | 1000 ng/mL | Intracellular | Hu, Ms, Rt |
| Pd102 | Barcode 1 | – | – | – | Trace Sciences International | Contact vendor | – | – | 50 nM–1 µM | Barcode | – |
| Pd104 | Barcode 2 | – | – | – | Trace Sciences International | Contact vendor | – | – | 50 nM–1 µM | Barcode | – |
| Pd105 | Barcode 3 | – | – | – | Trace Sciences International | Contact vendor | – | – | 50 nM–1 µM | Barcode | – |
| Pd106 | Barcode 4 | – | – | – | Trace Sciences International | Contact vendor | – | – | 50 nM–1 µM | Barcode | – |
| Pd108 | Barcode 5 | – | – | – | Trace Sciences International | Contact vendor | – | – | 50 nM–1 µM | Barcode | – |
| Pd110 | Barcode 6 | – | – | – | Trace Sciences International | Contact vendor | – | – | 50 nM–1 µM | Barcode | – |
| In113 | Olig2 | Oligodendrocyte transcription factor 2 | OPCs, Cholinergic Neuronal Progenitors, Interneuron Progenitors | Lu, 2000; Zhou, 2000 | Millipore | MABN50 | Mouse | 211F1.1 | 1000 ng/mL | Intracellular | Hu, Ms, Rt |
| In115 | A2B5 | – | O2A Progenitors, Type 2 Astrocytes | Eisenbarth, 1979; Raff, 1983 | Biolegend | 150702 | Mouse | 105/A2B5 | 300 ng/mL | Surface | Hu, Ms |
| La139 | PSA-NCAM | Polysialylated-neural cell adhesion molecule | Neuronal Progenitors | Theiry, 1979 | eBioscience | 14-9118-82 | Mouse | 12E3 | 300 ng/mL | Surface | Hu, Ms, Rt |
| Pr141 | CD140b | Platelet-derived growth factor beta | OPCs, Fibroblasts | Richardson, 1988; Noble, 1988 | eBioscience | 14-1402-82 | Rat | APB5 | 10 ng/mL | Surface | Hu, Ms, Fish |
| Nd142 | VCAM1 | CD106, Vascular cell adhesion molecule | NSCs, Glial Progenitors, and Neurovascular Cells | Osborn, 1989; Kokovay, 2012 | Biolegend | 105702 | Mouse | 429 (MVCAM.A) | 100 ng/mL | Surface | Ms |
| Nd143 | CD31 | PECAM-1, Endothelial cell adhesion molecule | Endothelial cells | Newman, 1990; Vasudevan, 2008 | Biolegend | 102425 | Rat | 390 | 50 ng/mL | Surface | Ms |
| Nd144 | Nestin | – | NSCs, Glial Progenitors, and Neurovascular Cells | Dahlstrand, 1992 | R&D Systems | MAB2736 | Mouse | 307501 | 30 ng/mL | Intracellular | Ms, Rt |
| Nd145 | Sox1 | SRY-box transcription factor 1 | Neural stem cells (especially neurogenic striatal progenitors) | Gubbay, 1990; Pevny, 1998 | R&D Systems | AF3369 | Goat | Polyclonal | 300 ng/mL | Intracellular | Hu, Ms, Rt |
| Nd146 | Tbr2 | T-box brain gene 2, EOMES, eomesdermin | Intermediate neuronal progenitors | Ross, 2000 | Thermo Fisher | 14-4875-82 | Mouse | Dan11mag | 1000 ng/mL | Intracellular | Ms |
| Sm147 | CD140a | Platelet-derived growth factor alpha | OPCs and brain fibroblasts | Pringle, 1989 | Biolegend | 135902 | Rat | APA5 | 30 ng/mL | Surface | Ms |
| Nd148 | CD133 | Prominin | Ependymal cells; apical neural progenitors | Weigmann, 1997 | Biolegend | 141202 | Rat | 315-2C11 | 100 ng/mL | Surface | Ms |
| Sm149 | CD45 | Tyrosine phosphatase receptor type C | Microglia and other leukocytes | Akiyama, 1988 | Fluidigm | 3089005B | Mouse | 30-F11 | 10 ng/mL | Surface | Ms |
| Nd150 | NeuN | Neuronal Nuclei | Neurons | Mullen, 1992 | Novus | NBP1-92693 | Mouse | 1B7 | 100 ng/mL | Intracellular | Hu, Ms, Rt |
| Eu151 | Sox10 | Sex-determining region Y box 10 | OPCs | Hu, 2009 | Gift (S. Kucenas) | – | Rabbit | Monoclonal | 3000 ng/mL | Intracellular | Ms, Rt, Fish |
| Sm152 | Ki67 | MKI67, Marker of proliferation Ki-67 | Proliferative cells | Gerdes, 1983 | BD Biosciences | 550609 | Mouse | B56 | 1000 ng/mL | Intracellular | Hu, Ms |
| Eu153 | Oligo O4 | Oligodendrocyte marker O4 | Oligodendrocytes | Yokoyama, 2003 | R&D Systems | MAB1326 | Mouse | O4 | 300 ng/mL | Surface | Hu, Ms, Rt, Ck |
| Sm154 | Pax6 | Paired box 6 | Neural stem cells, dorsal endothelial progenitors, dorsal white matter inhibitory neurons | Walther, 1991; Vasudevan, 2008; Riccio, 2011 | BD Biosciences | 561462 | Mouse | O18-1330 | 50 ng/mL | Intracellular | Hu, Ms |
| Gd155 | MCAM | CD146, Melanoma cell adhesion molecule | Neurovascular cells (endothelial cells and pericytes) | Schwarz, 1998 | eBioscience | 14-1469-80 | Mouse | P1H12 | 30 ng/mL | Surface | Hu, Ms, Rb |
| Gd156 | NeuroD1 | Neurogenic differentiation 1 | Differentiating neurons | Lee, 1998 | R&D Systems | AF2746 | Goat | Polyclonal | 1000 ng/mL | Intracellular | Hu, Ms |
| Gd157 | SSEA-1 | CD15, Stage-specific embryonic antigen, Sialyl LewisX | Stem cells | Knowles, 1978 | eBioscience | 14-8813-80 | Mouse | MC-480 | 10 ng/mL | Surface | Hu, Ms |
| Gd158 | CD11b | Integrin aN, Mac-1 | Microglia and monocytes | Perry, 1985 | Biolegend | 101249 | Rat | M1/70 | 100 ng/mL | Surface | Hu, Ms |
| Tb159 | CD24 | Heat stable antigen | Postmitotic neural cells | Shirasawa, 1993 | BD Biosciences | 557436 | Rat | m1/69 | 30 ng/mL | Surface | Ms |
| Gd160 | Sox2 | Sex-determining region Y box 2 | Neural stem cells, early embryonic endothelial progenitors | Uwanogho, 1995; Boström, 2018 | R&D Systems | MAB2018 | Mouse | 245610 | 1000 ng/mL | Intracellular | Hu, Ms, Rt |
| Dy161 | CD325 | N-cadherin | Most neural cells | Hatta, 1985 | Biolegend | 844702 | Mouse | 13A9 | 100 ng/mL | Surface | Hu, Ms, Rt |
| Dy162 | GAD65 | Glutamic acid decarboxylase 65-kD, glutamate decarboxylase 2 | Interneurons | Kaufman, 1991 | Biolegend | 844502 | Mouse | N-GAD65 | 300 ng/mL | Intracellular | Hu, Ms, Rt |
| Dy163 | DCX | Doublecortin | Newborn neurons | des Portes, 1998 | Thermo Fisher | 481200 | Rabbit | Polyclonal | 500 ng/mL | Intracellular | Hu, Ms, Rt |
| Dy164 | MAP2 | Microtubule-associated protein 2 | Differentiated neurons, preoligodendrocytes | Izant and McIntosh, 1980; Vouyiouklis, 1995 | Novus | NBP2-25156 | Mouse | 4H5 | 500 ng/mL | Intracellular | Hu, Ms |
| Ho165 | ALDH1A1 | Aldehyde dehydrogenase 1A1 | Dopaminergic neurons, astrocyte progenitors | Galter, 2003; Adam, 2012 | R&D Systems | AF5869 | Goat | Polyclonal | 100 ng/mL | Intracellular | Hu, Ms |
| Er166 | Ly-6C | Lymphocyte antigen 6 complex, locus C | T cells | LeClair, 1989 | Biolegend | 128002 | Rat | HK1.4 | 30 ng/mL | Surface | Ms |
| Er167 | GLAST | Glutamate asparate transporter | Glial cells | Storck, 1992 | Novus | NB100-1869 | Rabbit | Polyclonal | 1000 ng/mL | Surface | Hu, Ms, Rt |
| Er168 | Ctip2 | COUP-TF-interacting protein 2, Bcl11b | Developing neurons | Yamamato, 1999 | Abcam | ab18465 | Rat | 25B6 | 200 ng/mL | Intracellular | Hu, Ms |
| Tm169 | GFAP | Glial fibrillary acidic protein | Astrocytes | Uyeda, 1972 | BD Biosciences | 556330 | Mouse | 102 | 100 ng/mL | Intracellular | Hu, Ms, Rt |
| Er170 | Cux1 | Cut-like homeobox 1 | Neural cells | Quaggin, 1996 | Abcam | ab54583 | Mouse | 2A10 | 10 ng/mL | Intracellular | Hu, Ms |
| Yb171 | Tbr1 | T-box brain gene 1 | Neurons (predominantly cortical and cerebellar) | Bulfone, 1995 | Abcam | ab31940 | Rabbit | Polyclonal | 1000 ng/mL | Intracellular | Hu, Ms, Rt |
| Yb172 | BLBP | Brain lipid-binding protein, fatty acid binding protein 7 | Radial glial cells, astrocytes | Feng, 1994; Arnold, 1994 | Gift (C. Birchmeier and T. Meuller; Kurtz et al., 1994) | – | Rabbit | Polyclonal | 1000 ng/mL | Intracellular | Hu, Ms, Rt, Fish |
| Yb173 | Cl. Casp-3 | Caspase 3 (Cleaved Form) | Apoptotic Cells | Fernandes-Alnemri, 1994 | BD Biosciences | 559565 | Rabbit | C92-605 | 500 ng/mL | Intracellular | Hu, Ms |
| Yb174 | TrkB | Neurotrophic tyrosine kinase receptor type 2 | Developing neural cells | Klein, 1989 | Thermo Fisher | AF1494 | Goat | Polyclonal | 100 ng/mL | Surface | Ms |
| Yb175 | F4/80 | EMR1, Ly-71 | Macrophages | Austyn, 1981 | Biolegend | 123101 | Rat | BM8 | 100 ng/mL | Surface | Ms |
| Yb176 | p75NTR | P75 neurotrophic receptor, TNF receptor 16 | Neural cells | Herrup, 1973 | R&D Systems | AF1157 | Goat | Polyclonal | 300 ng/mL | Surface | Ms |
| Ir191/193 | DNA Intercalator | Cell-ID Intercalator-Ir | Live cell membrane-impermeable cationic nucleic acid intercalator | – | Fluidigm | 201192A | - | - | 1:5000 dilution | Post-Stain | Eukaryotic cells |
| Pt195/198 | Cisplatin | Cisplatin | Dead cells | Fienberg, 2012 | Sigma Aldrich | | - | - | 5 µM | Pre-Fix | Non-viable cells |

Ck, chicken; Hu, human; Ms, mouse; Rt, rat
