## Extended Data Figures for "A developmental atlas of the mouse brain by single-cell mass cytometry"

#### Extended Data Fig. 1

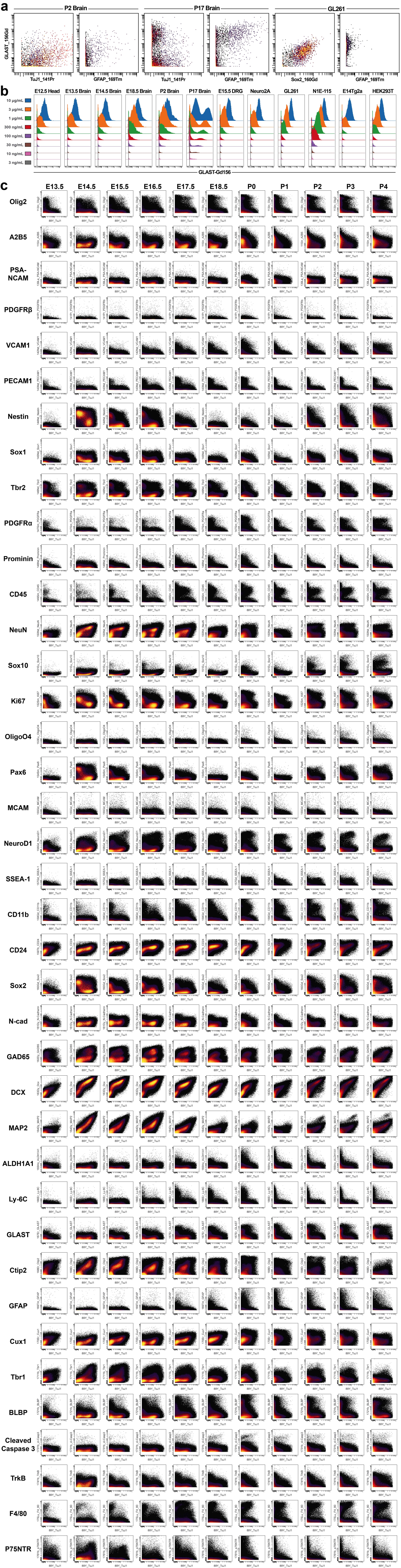

#### Extended Data Fig. 2

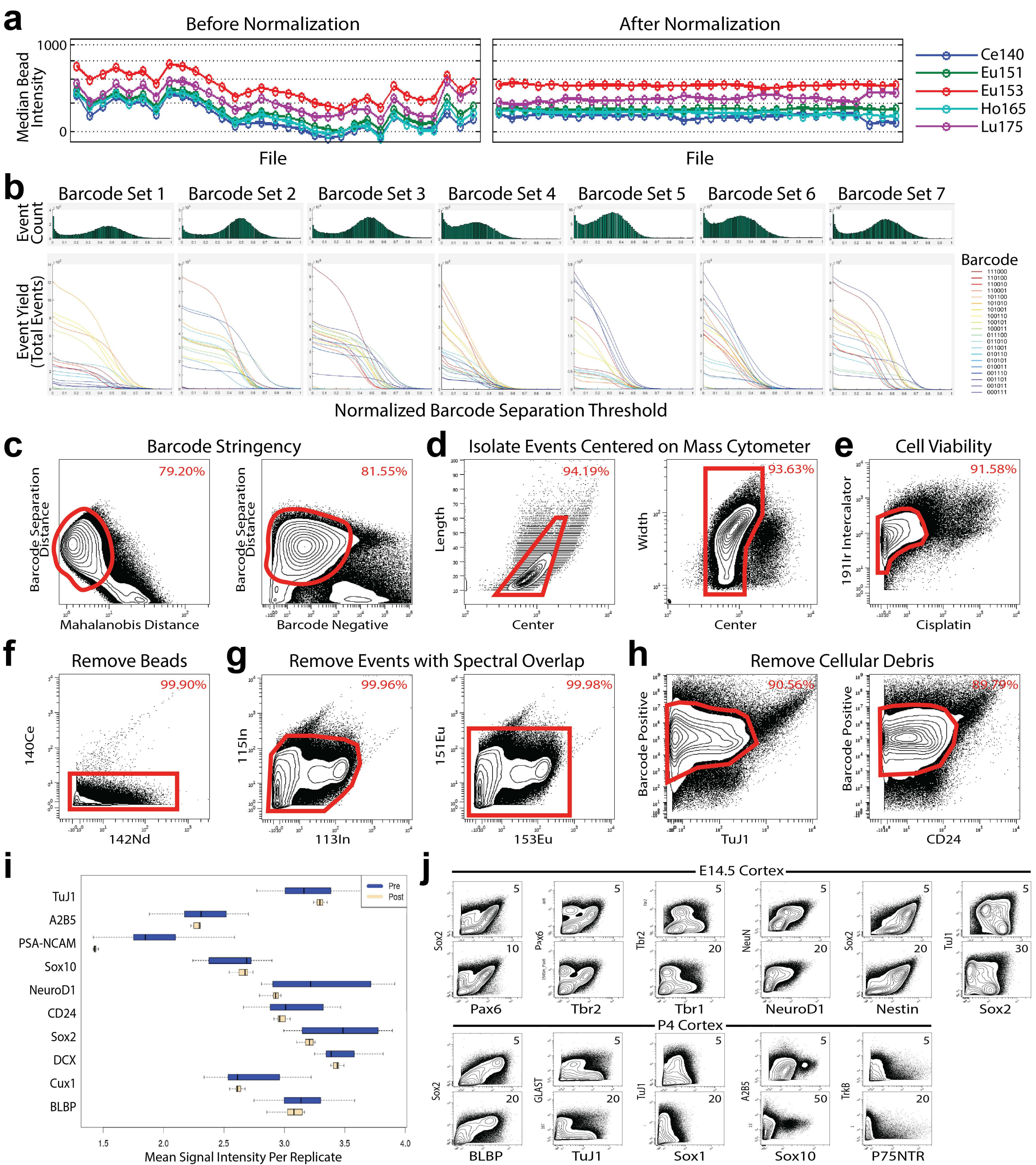

### Extended Data Fig. 3

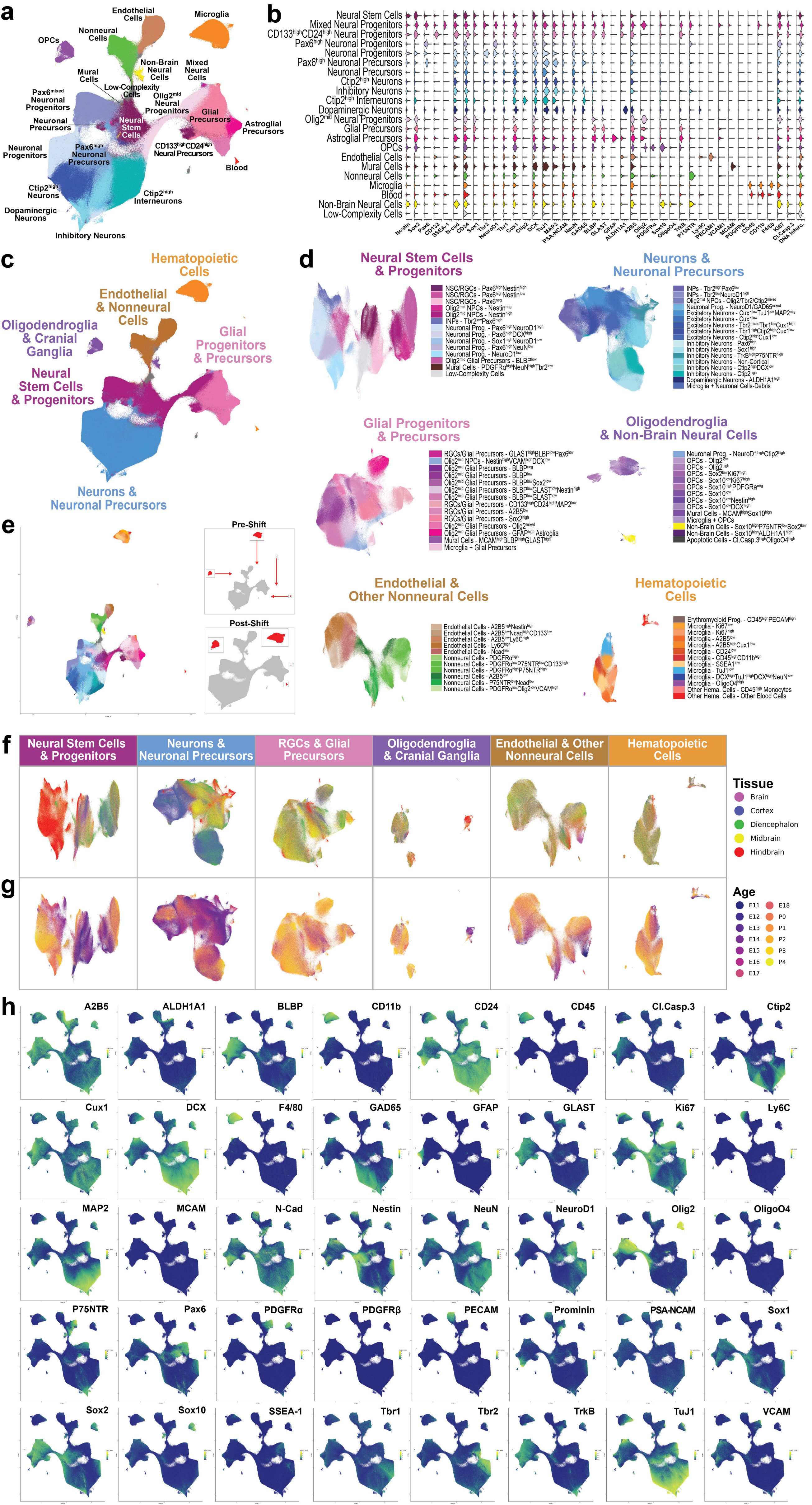

**Extended Data Fig. 4**

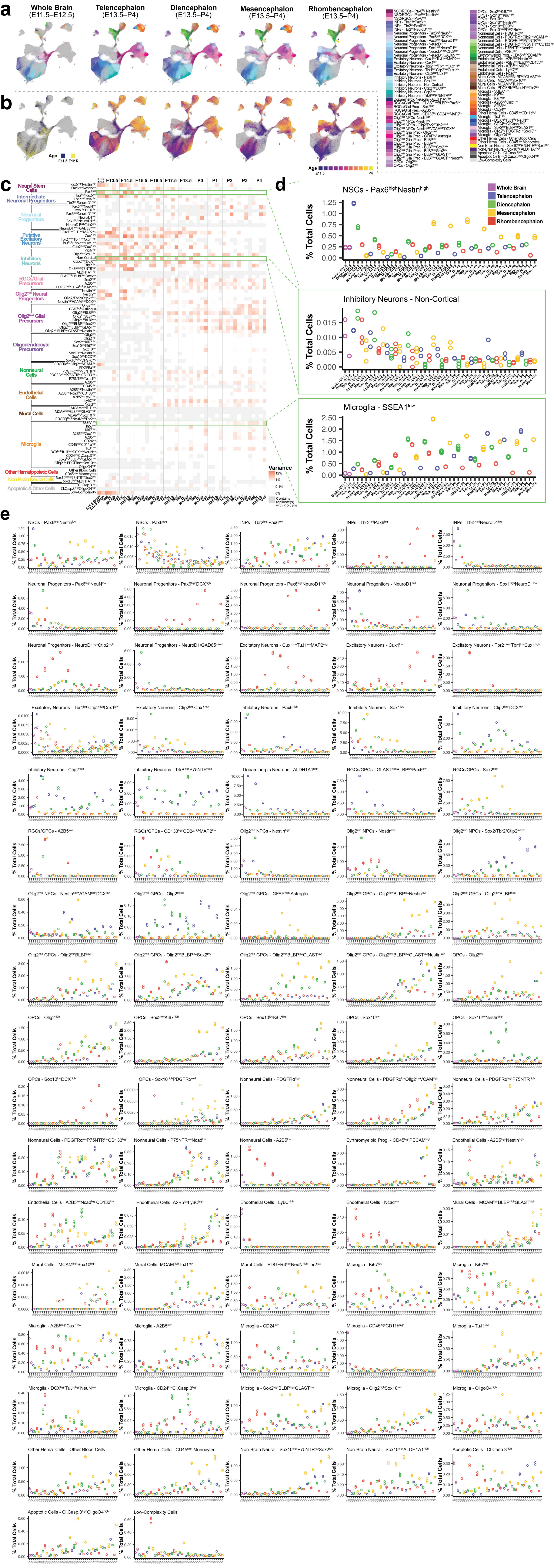

### Extended Data Fig. 5

**a**

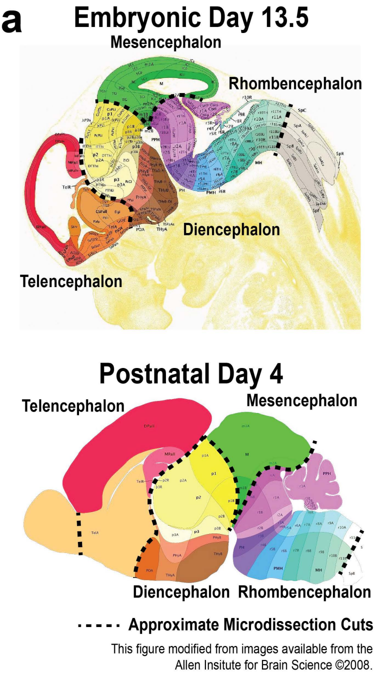


**b**

| Age | Tissue in Figures | Tissues in La Manno, 2021 | Ages in La Manno, 2021 |
|---|---|---|---|
| **E11.5** | Whole Brain | Forebrain, Midbrain, Hindbrain | E11.0 |
| **E12.5** | Whole Brain | Forebrain (Dorsal), Forebrain (Ventral), Midbrain, Hindbrain | E12.0, E12.5 |
| **E13.5** | Telencephalon | Forebrain (Dorsal) | E13.0, E13.5 |
| **E14.5** | Telencephalon | Forebrain (Dorsal) | E14.0, E14.5 |
| **E15.5** | Telencephalon | Forebrain (Dorsal) | E15.0, E15.5 |
| **E16.5** | Telencephalon | Forebrain (Dorsal) | E16.0, E16.25, E16.5 |
| **E17.5** | Telencephalon | Forebrain (Dorsal) | E17.0, E17.5 |
| **E18.5** | Telencephalon | Forebrain (Dorsal) | E18.0 |
| **E13.5** | Diencephalon | Forebrain (Ventral) | E13.0, E13.5 |
| **E14.5** | Diencephalon | Forebrain (Ventral) | E14.0, E14.5 |
| **E15.5** | Diencephalon | Forebrain (Ventral) | E15.0, E15.5 |
| **E16.5** | Diencephalon | Forebrain (Ventrothalamic), Forebrain (Ventrolateral) | E16.0, E16.5 |
| **E17.5** | Diencephalon | Forebrain (Ventrothalamic), Forebrain (Ventrolateral) | E17.5 |
| **E18.5** | Diencephalon | Forebrain (Ventrothalamic), Forebrain (Ventrolateral) | E18.0 |
| **E13.5** | Mesencephalon | Midbrain | E13.0, E13.5 |
| **E14.5** | Mesencephalon | Midbrain, Midbrain (Dorsal), Midbrain (Ventral) | E14.0, E14.5 |
| **E15.5** | Mesencephalon | Midbrain, Midbrain (Dorsal), Midbrain (Ventral) | E15.0, E15.5 |
| **E16.5** | Mesencephalon | Midbrain | E16.0, E16.5 |
| **E17.5** | Mesencephalon | Midbrain | E17.5 |
| **E18.5** | Mesencephalon | Midbrain | E18.0 |
| **E13.5** | Rhombencephalon | Hindbrain | E13.0, E13.5 |
| **E14.5** | Rhombencephalon | Hindbrain | E14.0, E14.5 |
| **E15.5** | Rhombencephalon | Hindbrain | E15.0, E15.5 |
| **E16.5** | Rhombencephalon | Hindbrain | E16.0, E16.5 |
| **E17.5** | Rhombencephalon | Hindbrain | E17.0, E17.5 |
| **E18.5** | Rhombencephalon | Hindbrain | E18.0 |

**c**

| Marker | % Positive Threshold for Mass Cytometry | Protein vs. RNA Class |
|---|---|---|
| **Sox2** | 1.67 | Transcription Factor |
| **Pax6** | 0.8 | Transcription Factor |
| **Tbr2** | 0.8 | Transcription Factor |
| **NeuroD1** | 1.5 | Transcription Factor |
| **Tbr1** | 0.75 | Transcription Factor |
| **Ctip2** | 0.4 | Transcription Factor |
| **Cux1** | 3 | Transcription Factor |
| **Olig2** | 1.15 | Transcription Factor |
| **Sox10** | 2.7 | Transcription Factor |
| **Sox1** | 0.6 | Transcription Factor |
| **Nestin** | 0.45 | Neurofilament |
| **DCX** | 1.3 | Neurofilament |
| **TuJ1** | 3.3 | Neurofilament |
| **MAP2** | 0.7 | Neurofilament |
| **GFAP** | 2 | Neurofilament |
| **PDGFRa** | 0.6 | Neurotrophic Receptor |
| **PDGFRb** | 0.7 | Neurotrophic Receptor |
| **TrkB** | 1 | Neurotrophic Receptor |
| **P75NTR** | 1.3 | Neurotrophic Receptor |
| **Prominin** | 0.5 | Cell Surface Molecule |
| **GLAST** | 2.8 | Cell Surface Molecule |
| **CD45** | 1.5 | Cell Surface Molecule |
| **F4/80** | 2 | Cell Surface Molecule |
| **N-Cad** | 1 | Cell Adhesion Molecule |
| **CD11b** | 2 | Cell Adhesion Molecule |
| **VCAM1** | 1.2 | Cell Adhesion Molecule |
| **MCAM** | 4 | Cell Adhesion Molecule |
| **PECAM1** | 2 | Cell Adhesion Molecule |
| **GAD65** | 0.5 | Enzyme |
| **ALDH1A1** | 2 | Enzyme |
| **SSEA-1** | 2 | Enzyme |
| **NeuN** | 1.5 | Accessory Protein |
| **BLBP** | 1 | Accessory Protein |
| **Ki67** | 4 | Accessory Protein |
| **A2B5** | – | RNA Data Unavailable |
| **CD24** | – | RNA Data Unavailable |
| **Cl.Casp.3** | – | RNA Data Unavailable |
| **Ly-6C** | – | RNA Data Unavailable |
| **OligoO4** | – | RNA Data Unavailable |
| **PSA-NCAM** | – | RNA Data Unavailable |

**d**

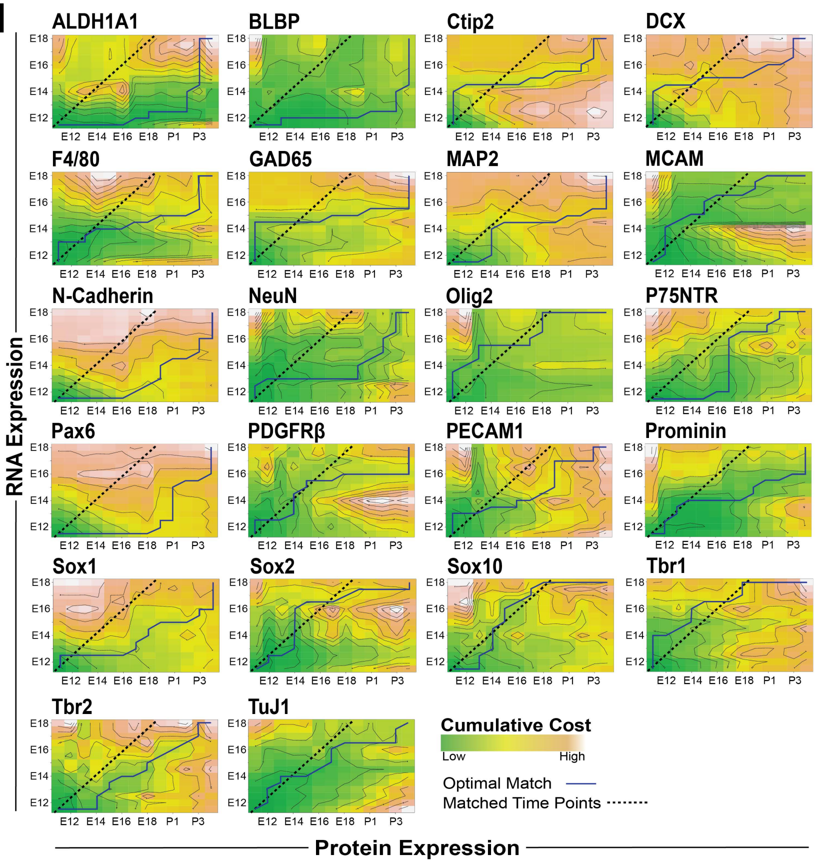

### Extended Data Fig. 6

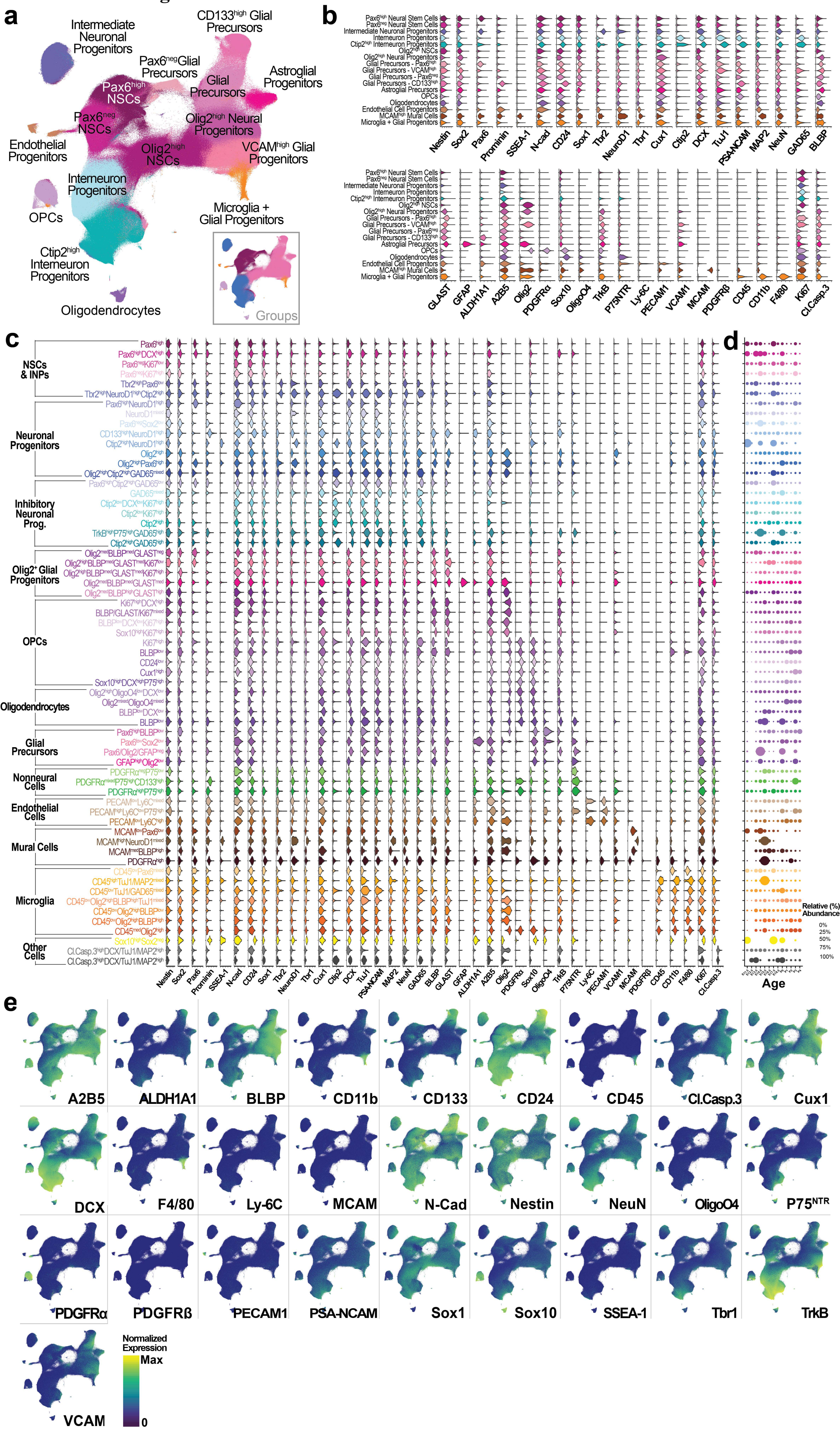

### Extended Data Fig. 7

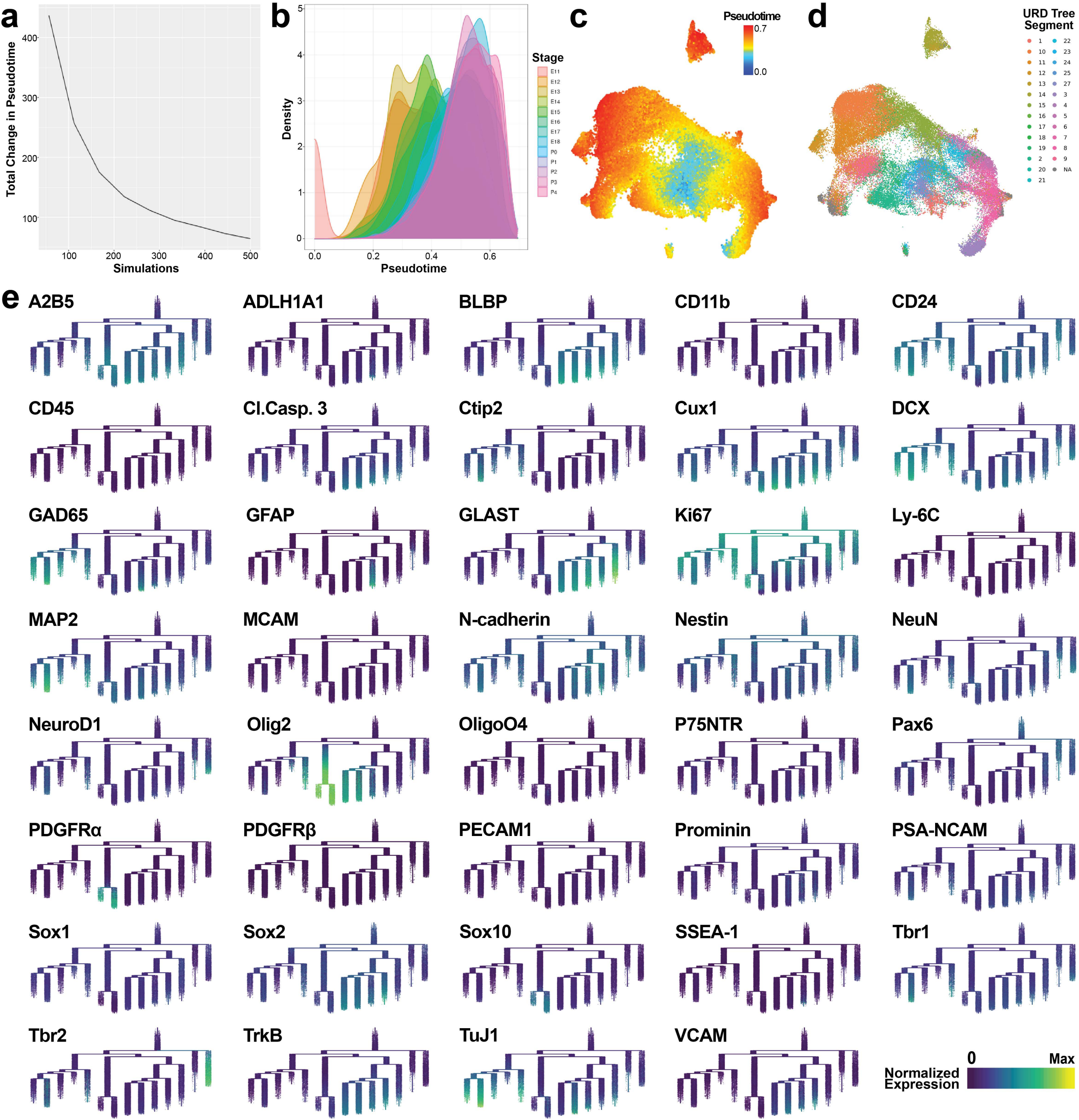

**Extended Data Fig. 8**

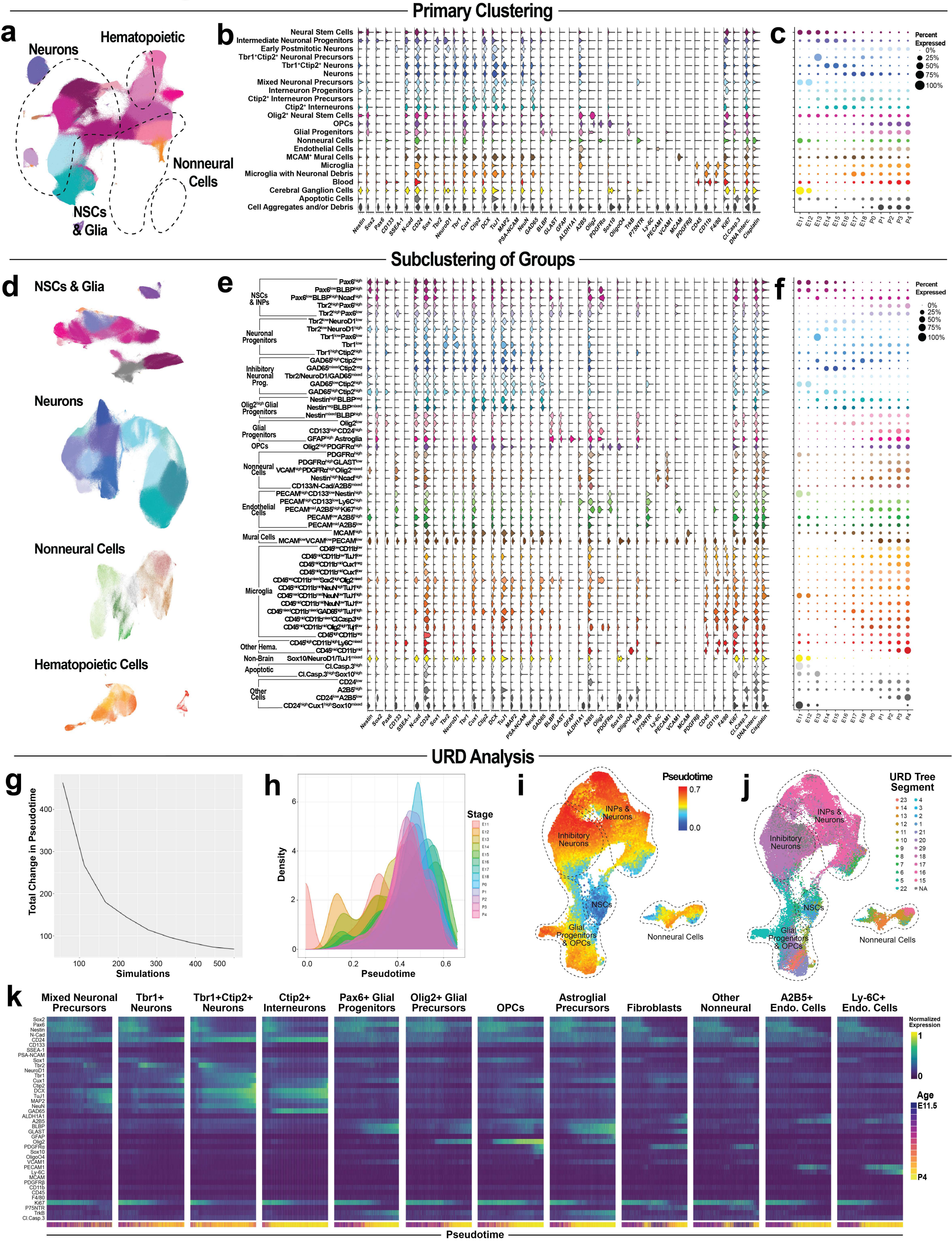

### Extended Data Fig. 9

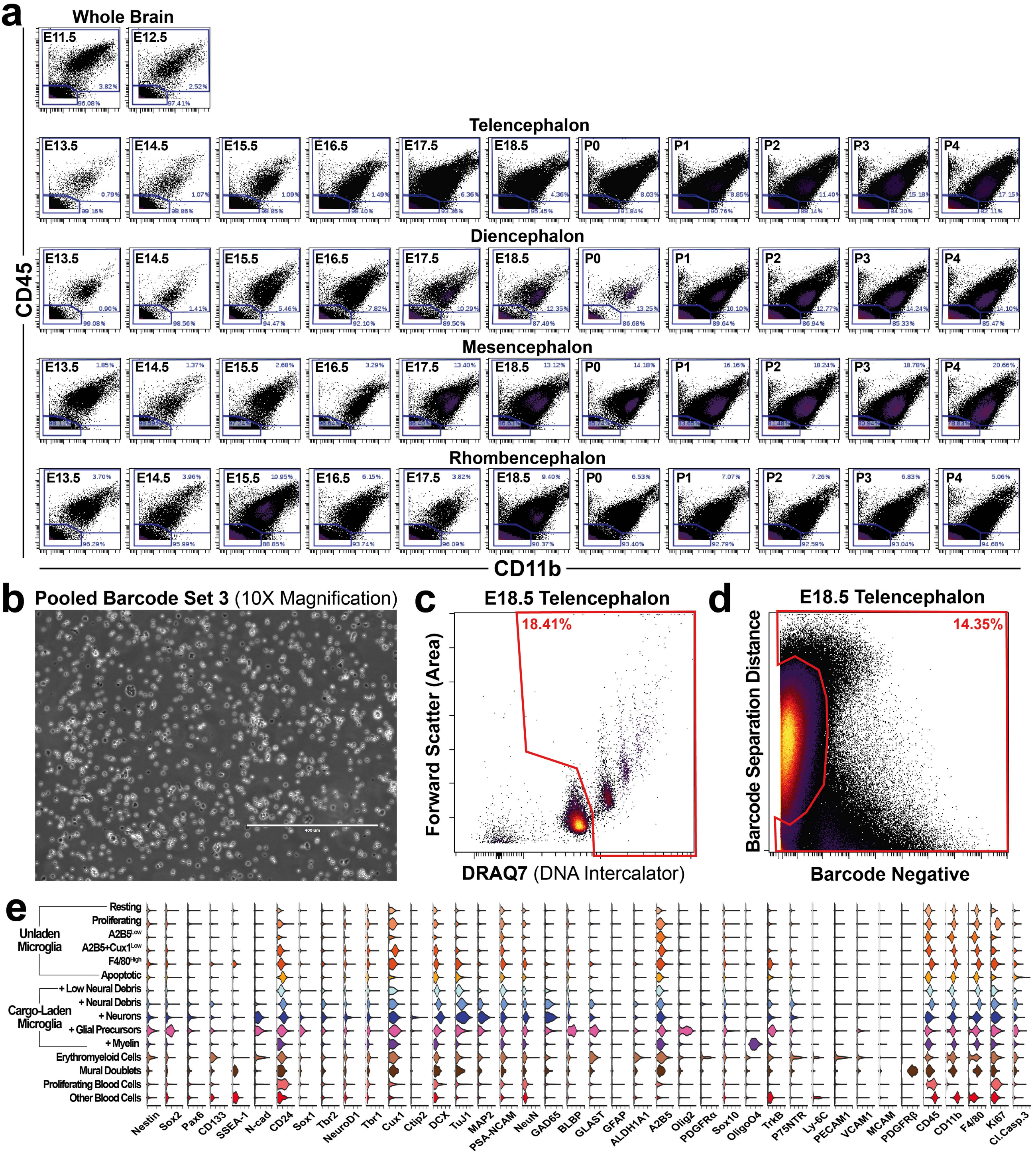
