## Extended Data Figures Legends and Bibliography for "A developmental atlas of the mouse brain by single-cell mass cytometry"

**Extended Data Table 1.** Antibodies used for mass cytometry.

**Extended Data Fig. 1. Titration and validation of antibodies for mass cytometry. a**, Biaxial plots (CytoBank) showing 1 μg/mL GLAST-Gd156 antibody versus known-negative (TuJ1) and known-positive (GFAP and Sox2) markers in primary cells harvested from P2 and P17 brain, and the GL261 mouse glioma cell line. **b**, Results for titration of GLAST-Gd156 antibody in primary cells harvested from mice [E12.5 head, E13.5/E14.5/E18.5/P2/P17 brain, E15.5 dorsal root ganglia (DRG)], mouse cell lines [Neuro2a (neuroblastoma), GL261 (glioma), N1E-115 (neuroblastoma), E14Tg2a (embryonic stem cells)], and a human cell line [HEK293T (embryonic kidney)]. **c**, Biaxial plots (CytoBank) showing protein expression of the neuronal marker TuJ1 versus every antibody included in the panel in a single replicate of each age from the first mass cytometry run (barcode sets 1–3) for the telencephalon.

**Extended Data Fig. 2. Pre-processing of mass cytometry data for the developing mouse brain. a**, Normalization of raw data (.fcs files) with calibration beads using the Matlab software described in Finck, et al. *Cytometry A*, 2013. **b**, Debarcoding of normalized .fcs files using the R package described in Fread, et al. *Pac Symp Biocomput*, 2017. Bar graphs on top show numbers of events with each normalized barcode separation threshold value for all samples. Line graphs on bottom show yields of events for each debarcoded sample. **c–h**, Biaxial plots (CytoBank) showing gating (indicated in red) used to isolate single cells from developing mouse brains. Percentages in the top right corner of each plot indicate the ratio of ungated cells within the gate. **i**, Mean signal intensity and variance in pre- and post-batch corrected debarcoded .fcs files, as processed with the R package described in Schuyler, et al. *Front Immunol*, 2019. Markers not shown had a mean signal variance < 0.01 and were not corrected. **j**, Biaxial plots (CytoBank) comparing default (ASinH = 5) and manually optimized ASinH values in E14.5 or P4 cortex. ASinH values for markers on the x-axis are indicated at the top right corner of each panel. Markers not shown were transformed using the default value.

**Extended Data Fig. 3. Classification of cells in the developing mouse brain by mass cytometry. a**, UMAP colored and labeled according to Round 1 Leiden clustering. **b**, Violin plot showing expression of all 40 protein markers in Round 1 Leiden clusters. **c**, UMAP colored according to the six groups of Round 1 clusters that were subjected to a second round of Leiden clustering. Groups were manually assigned according to molecular expression profile and UMAP layout. **d**, UMAPs showing Round 2 Leiden clustering of each group. **e**, Original UMAP layout (corresponding to Fig. 1C–E) and shifts used for compression into the final UMAP layout. **f**, Round 2 UMAPs for each group colored according to the brain region from which each cell originated. **g**, Round 2 UMAPs for each group colored according to the developmental age (E11.5–P4) of samples. n = 2–5 litters per age for a total of 5,750,000 cells from 112 samples.

**Extended Data Fig. 4. Profiling cell abundances in the developing mouse telencephalon, diencephalon, mesencephalon, and rhombencephalon. a**, UMAPs colored according to Round 2 Leiden clustering assignment for each microdissected brain region; cells in other brain regions are colored grey. **b**, UMAPs colored according to the developmental age (E11.5–P4) of each cell. **c**, Heatmap of cluster variance for each sample type (e.g. E11.5 whole brain, P4 telencephalon). Low-abundance sample types (one or more replicate with < 5 cells) were excluded from variance analysis and are colored grey. **d** and **e**, Scatter plots showing relative abundances of each cluster for each sample type for corresponding clusters in (**c**). Three clusters are highlighted in **d**, while the remaining clusters are shown in **e**. n = 2–5 litters per age for a total of 5,750,000 cells from 112 samples.

**Extended Data Fig. 5. Comparison of protein and mRNA expression profiles in the developing mouse brain. a**, Adapted images from the Allen Brain Atlas of the Developing Mouse Brain showing approximate microdissection of E13.5–P4 mouse brain samples. Calculations of voxel intensity (as a measure of mRNA expression) in Fig. 3a–g for Allen Brain Atlas *in situ* hybridization (ISH) data were grouped for each brain region according to the dashed lines. The olfactory bulb was discarded and not included in analyses. **b**, Overview of how results from the present study were compared sample-by-sample with the scRNA-seq results of La Manno, et al. *Nature*, 2021. **c**, Overview of genes included in the antibody panel and, if used in comparative analyses, the threshold value applied to calculate relative abundances of positive and negative cells for ISH, scRNA-seq, and mass cytometry datasets. **d**, Dynamic time-warp analysis of lag between relative RNA and protein expression levels in positively expressing cells for markers not shown in Fig. 3h. Highly similar relative expression levels are indicated in green, while low-similarity levels are indicated in orange. The solid blue line indicates the optimal match calculated by the dynamic time-warp algorithm, while the black dashed line indicates the theoretical perfect correlation between relative RNA and protein expression levels at each overlapping time point (i.e., lag = 0).

**Extended Data Fig. 6. Survey of cells with stem-like properties in the developing mouse brain. a**, UMAP colored and labeled according to R1 Leiden clustering assignment. Inset shows UMAP colored according to the five groups of Round 1 clusters that were subjected to a second round of Leiden clustering. **b**, Violin plot showing expression of all 40 protein markers in Round 1 Leiden clusters. **c**, Violin plot showing expression of all 40 protein markers in Round 2 Leiden clusters. **d**, Dot plot showing relative abundances of clusters at each developmental age for all brain regions together. **e**, UMAPs colored according to expression levels of markers not included in Figure 4e. n = 2–5 litters per age for a total of 3,253,024 cells in 112 samples.

**Extended Data Fig. 7. Exploring neural cell fates in the developing mouse brain with pseudotime-based analysis of molecular trajectories. a**, Change in total stability of URD pseudotime calculations with number of simulations. **b**, Histogram showing relative cell density for each developmental age (E11.5–P4) for each URD pseudotime bin. **c**, UMAP generated by URD algorithm [only 6.1 x $10^4$ cells (1.9% of total $Sox2^+Nestin^+$ cells) included in URD analysis] colored according to pseudotime. **d**, UMAP generated by URD algorithm colored according to URD dendrogram tree segment. **e**, URD dendrograms colored according to protein expression levels of the indicated markers. n = 2–5 litters per age for a total of 60,955 cells from 112 samples.

**Extended Data Fig. 8. Identifying molecular trajectories underlying cell specification in the developing mouse telencephalon. a**, UMAP colored according to Round 1 Leiden clustering assignment. Dashed lines indicate four groups of Round 1 clusters that were subjected to a second round of Leiden clustering. **b**, Violin plot showing expression of all 40 protein markers in Round 1 Leiden clusters. **c**, Dot plot showing relative abundances of Round 1 Leiden clusters at each developmental age. **d**, UMAPs showing Round 2 Leiden clustering of each group. **e**, Violin plot showing expression of all 40 protein markers in Round 2 Leiden clusters. **f**, Dot plot showing relative abundances of Round 2 Leiden clusters at each developmental age. **g**, Change in total stability of URD pseudotime calculations with number of simulations. **h**, Histogram showing relative cell density for each developmental age (E11.5–P4) for each URD pseudotime bin. **i**, UMAP generated by the URD algorithm (only includes $6.1 \times 10^4$ cells included in URD analysis) colored according to pseudotime. **j**, UMAP generated by the URD algorithm colored according to the URD dendrogram tree segment. **k**, Heatmaps colored by protein expression of all 40 markers for select trajectories identified by URD analysis. For **a–f**, n = 2 litters per age for a total of 3,048,448 cells from 26 samples. For **g–k**, n = 2 litters per age for a total of 61,006 cells from 26 samples.

**Extended Data Fig. 9. Identifying functionally distinct microglia subpopulations in the developing mouse brain.** **a**, Biaxial plots showing gating of all single-cell events from one replicate of each sample type. Cells in upper gate were retained as $CD45^+$ cells. **b**, Violin plot showing expression of all 40 protein markers in Leiden clusters. n = 2–5 litters per age/brain region for a total 1,758,016 cells from 112 samples. **c**, Bright field image of pooled Barcode Set 3 (containing P1–P4 samples) prior to mass cytometry analysis (10× magnification, scale bar = 400 μm).

**Extended Data Bibliography**


Adam, S. Alexandra, Oliver Schnell, Julia Pöschl, Sabina Eigenbrod, Hans A. Kretzschmar, Jörg-Christian Tonn, and Ulrich Schüller. "ALDH1A1 Is a Marker of Astrocytic Differentiation during Brain Development and Correlates with Better Survival in Glioblastoma Patients." *Brain Pathology (Zurich, Switzerland)* 22, no. 6 (November 2012): 788–97. https://doi.org/10.1111/j.1750-3639.2012.00592.x.

Akiyama, H., S. Itagaki, and P. L. McGeer. "Major Histocompatibility Complex Antigen Expression on Rat Microglia Following Epidural Kainic Acid Lesions." *Journal of Neuroscience Research* 20, no. 2 (1988): 147–57. https://doi.org/10.1002/jnr.490200202.

Arnold, D., L. Feng, J. Kim, and N. Heintz. "A Strategy for the Analysis of Gene Expression during Neural Development." *Proceedings of the National Academy of Sciences of the United States of America* 91, no. 21 (October 11, 1994): 9970–74. https://doi.org/10.1073/pnas.91.21.9970.

Austyn, J. M., and S. Gordon. "F4/80, a Monoclonal Antibody Directed Specifically against the Mouse Macrophage." *European Journal of Immunology* 11, no. 10 (October 1981): 805–15. https://doi.org/10.1002/eji.1830111013.

Boström, Kristina I., Jiayi Yao, Xiuju Wu, and Yucheng Yao. "Endothelial Cells May Have Tissue-Specific Origins." *Journal of Cell Biology and Histology* 1, no. 1 (June 2018): 104.

Bulfone, A., S. M. Smiga, K. Shimamura, A. Peterson, L. Puelles, and J. L. Rubenstein. "T-Brain-1: A Homolog of Brachyury Whose Expression Defines Molecularly Distinct Domains within the Cerebral Cortex." *Neuron* 15, no. 1 (July 1995): 63–78. https://doi.org/10.1016/0896-6273(95)90065-9.

Caccamo, D. V., M. M. Herman, A. Frankfurter, C. D. Katsetos, V. P. Collins, and L. J. Rubinstein. "An Immunohistochemical Study of Neuropeptides and Neuronal Cytoskeletal Proteins in the Neuroepithelial Component of a Spontaneous Murine Ovarian Teratoma. Primitive Neuroepithelium Displays Immunoreactivity for Neuropeptides and Neuron-Associated Beta-Tubulin Isotype." *The American Journal of Pathology* 135, no. 5 (November 1989): 801–13.

"Characterization of the Human Nestin Gene Reveals a Close Evolutionary Relationship to Neurofilaments | Journal of Cell Science | The Company of Biologists." Accessed July 4, 2022. https://journals.biologists.com/jcs/article/103/2/589/23533/Characterization-of-the-human-nestin-gene-reveals.

"Conversion of Xenopus Ectoderm into Neurons by NeuroD, a Basic Helix-Loop-Helix Protein." Accessed July 4, 2022. https://www.science.org/doi/abs/10.1126/science.7754368.

Eisenbarth, G S, F S Walsh, and M Nirenberg. "Monoclonal Antibody to a Plasma Membrane Antigen of Neurons." *Proceedings of the National Academy of Sciences of the United States of America* 76, no. 10 (October 1979): 4913–17.

Feng, L., M. E. Hatten, and N. Heintz. "Brain Lipid-Binding Protein (BLBP): A Novel Signaling System in the Developing Mammalian CNS." *Neuron* 12, no. 4 (April 1994): 895–908. https://doi.org/10.1016/0896-6273(94)90341-7.

Fernandes-Alnemri, T, G Litwack, and E S Alnemri. "CPP32, a Novel Human Apoptotic Protein with Homology to Caenorhabditis Elegans Cell Death Protein Ced-3 and Mammalian Interleukin-1 Beta-Converting Enzyme." *Journal of Biological Chemistry* 269, no. 49 (December 1994): 30761–64. https://doi.org/10.1016/S0021-9258(18)47344-9.

Fienberg, Harris G., Erin F. Simonds, Wendy J. Fantl, Garry P. Nolan, and Bernd Bodenmiller. "A Platinum-Based Covalent Viability Reagent for Single-Cell Mass Cytometry." *Cytometry. Part*

*A: The Journal of the International Society for Analytical Cytology* 81, no. 6 (June 2012): 467–75. https://doi.org/10.1002/cyto.a.22067.
Galter, Dagmar, Silvia Buervenich, Andrea Carmine, Maria Anvret, and Lars Olson. "ALDH1 MRNA: Presence in Human Dopamine Neurons and Decreases in Substantia Nigra in Parkinson's Disease and in the Ventral Tegmental Area in Schizophrenia." *Neurobiology of Disease* 14, no. 3 (December 2003): 637–47. https://doi.org/10.1016/j.nbd.2003.09.001.
Gerdes, J., U. Schwab, H. Lemke, and H. Stein. "Production of a Mouse Monoclonal Antibody Reactive with a Human Nuclear Antigen Associated with Cell Proliferation." *International Journal of Cancer* 31, no. 1 (January 15, 1983): 13–20. https://doi.org/10.1002/ijc.2910310104.
Gubbay, John, Jérôme Collignon, Peter Koopman, Blanche Capel, Androulla Economou, Andrea Münsterberg, Nigel Vivian, Peter Goodfellow, and Robin Lovell-Badge. "A Gene Mapping to the Sex-Determining Region of the Mouse Y Chromosome Is a Member of a Novel Family of Embryonically Expressed Genes." *Nature* 346, no. 6281 (July 1990): 245–50. https://doi.org/10.1038/346245a0.
Hatta, K, T S Okada, and M Takeichi. "A Monoclonal Antibody Disrupting Calcium-Dependent Cell-Cell Adhesion of Brain Tissues: Possible Role of Its Target Antigen in Animal Pattern Formation." *Proceedings of the National Academy of Sciences of the United States of America* 82, no. 9 (May 1985): 2789–93.
Herrup, K., and E. M. Shooter. "Properties of the Beta Nerve Growth Factor Receptor of Avian Dorsal Root Ganglia." *Proceedings of the National Academy of Sciences of the United States of America* 70, no. 12 (December 1973): 3884–88. https://doi.org/10.1073/pnas.70.12.3884.
Hu, Bao-Yang, Zhong-Wei Du, Xue-Jun Li, Melvin Ayala, and Su-Chun Zhang. "Human Oligodendrocytes from Embryonic Stem Cells: Conserved SHH Signaling Networks and Divergent FGF Effects." *Development (Cambridge, England)* 136, no. 9 (May 2009): 1443–52. https://doi.org/10.1242/dev.029447.
Izant, J G, and J R McIntosh. "Microtubule-Associated Proteins: A Monoclonal Antibody to MAP2 Binds to Differentiated Neurons." *Proceedings of the National Academy of Sciences of the United States of America* 77, no. 8 (August 1980): 4741–45.
Kaufman, D. L., C. R. Houser, and A. J. Tobin. "Two Forms of the Gamma-Aminobutyric Acid Synthetic Enzyme Glutamate Decarboxylase Have Distinct Intraneuronal Distributions and Cofactor Interactions." *Journal of Neurochemistry* 56, no. 2 (February 1991): 720–23. https://doi.org/10.1111/j.1471-4159.1991.tb08211.x.
Klein, R, L F Parada, F Coulier, and M Barbacid. "TrkB, a Novel Tyrosine Protein Kinase Receptor Expressed during Mouse Neural Development." *The EMBO Journal* 8, no. 12 (December 1, 1989): 3701–9.
Kokovay, Erzsebet, Yue Wang, Gretchen Kusek, Rachel Wurster, Patty Lederman, Natalia Lowry, Qin Shen, and Sally Temple. "VCAM1 Is Essential to Maintain the Structure of the SVZ Niche and Acts as an Environmental Sensor to Regulate SVZ Lineage Progression." *Cell Stem Cell* 11, no. 2 (August 3, 2012): 220–30. https://doi.org/10.1016/j.stem.2012.06.016.
LeClair, K. P., M. M. Bridgett, F. J. Dumont, R. G. Palfree, U. Hämmerling, and A. L. Bothwell. "Kinetic Analysis of Ly-6 Gene Induction in a T Lymphoma by Interferons and Interleukin 1, and Demonstration of Ly-6 Inducibility in Diverse Cell Types." *European Journal of Immunology* 19, no. 7 (July 1989): 1233–39. https://doi.org/10.1002/eji.1830190713.
Lu, Q. R., D. Yuk, J. A. Alberta, Z. Zhu, I. Pawlitzky, J. Chan, A. P. McMahon, C. D. Stiles, and D. H. Rowitch. "Sonic Hedgehog--Regulated Oligodendrocyte Lineage Genes Encoding BHLH

Proteins in the Mammalian Central Nervous System." *Neuron* 25, no. 2 (February 2000): 317–29. https://doi.org/10.1016/s0896-6273(00)80897-1.

Mullen, R. J., C. R. Buck, and A. M. Smith. "NeuN, a Neuronal Specific Nuclear Protein in Vertebrates." *Development (Cambridge, England)* 116, no. 1 (September 1992): 201–11. https://doi.org/10.1242/dev.116.1.201.

"New Pool of Cortical Interneuron Precursors in the Early Postnatal Dorsal White Matter | Cerebral Cortex | Oxford Academic." Accessed July 4, 2022. https://academic.oup.com/cercor/article/22/1/86/364887?login=true.

Noble, M., K. Murray, P. Stroobant, M. D. Waterfield, and P. Riddle. "Platelet-Derived Growth Factor Promotes Division and Motility and Inhibits Premature Differentiation of the Oligodendrocyte/Type-2 Astrocyte Progenitor Cell." *Nature* 333, no. 6173 (June 9, 1988): 560–62. https://doi.org/10.1038/333560a0.

Osborn, Laurelee, Catherine Hession, Richard Tizard, Cornelia Vassallo, Stefan Luhowskyj, Gloria Chi-Rosso, and Roy Lobb. "Direct Expression Cloning of Vascular Cell Adhesion Molecule 1, a Cytokine-Induced Endothelial Protein That Binds to Lymphocytes." *Cell* 59, no. 6 (December 22, 1989): 1203–11. https://doi.org/10.1016/0092-8674(89)90775-7.

"PECAM-1 (CD31) Cloning and Relation to Adhesion Molecules of the Immunoglobulin Gene Superfamily." Accessed July 4, 2022. https://www.science.org/doi/abs/10.1126/science.1690453.

Perry, V. H., D. A. Hume, and S. Gordon. "Immunohistochemical Localization of Macrophages and Microglia in the Adult and Developing Mouse Brain." *Neuroscience* 15, no. 2 (June 1985): 313–26. https://doi.org/10.1016/0306-4522(85)90215-5.

Pevny, L.H., S. Sockanathan, M. Placzek, and R. Lovell-Badge. "A Role for SOX1 in Neural Determination." *Development* 125, no. 10 (May 15, 1998): 1967–78. https://doi.org/10.1242/dev.125.10.1967.

Portes, V. des, J. M. Pinard, P. Billuart, M. C. Vinet, A. Koulakoff, A. Carrié, A. Gelot, et al. "A Novel CNS Gene Required for Neuronal Migration and Involved in X-Linked Subcortical Laminar Heterotopia and Lissencephaly Syndrome." *Cell* 92, no. 1 (January 9, 1998): 51–61. https://doi.org/10.1016/s0092-8674(00)80898-3.

Pringle, N., E. J. Collarini, M. J. Mosley, C. H. Heldin, B. Westermark, and W. D. Richardson. "PDGF A Chain Homodimers Drive Proliferation of Bipotential (O-2A) Glial Progenitor Cells in the Developing Rat Optic Nerve." *The EMBO Journal* 8, no. 4 (April 1989): 1049–56. https://doi.org/10.1002/j.1460-2075.1989.tb03472.x.

Quaggin, S. E., G. B. Heuvel, K. Golden, R. Bodmer, and P. Igarashi. "Primary Structure, Neural-Specific Expression, and Chromosomal Localization of Cux-2, a Second Murine Homeobox Gene Related to Drosophila Cut." *The Journal of Biological Chemistry* 271, no. 37 (September 13, 1996): 22624–34. https://doi.org/10.1074/jbc.271.37.22624.

Raff, MC, ER Abney, J Cohen, R Lindsay, and M Noble. "Two Types of Astrocytes in Cultures of Developing Rat White Matter: Differences in Morphology, Surface Gangliosides, and Growth Characteristics." *The Journal of Neuroscience* 3, no. 6 (June 1, 1983): 1289–1300. https://doi.org/10.1523/JNEUROSCI.03-06-01289.1983.

Richardson, W. D., N. Pringle, M. J. Mosley, B. Westermark, and M. Dubois-Dalcq. "A Role for Platelet-Derived Growth Factor in Normal Gliogenesis in the Central Nervous System." *Cell* 53, no. 2 (April 22, 1988): 309–19. https://doi.org/10.1016/0092-8674(88)90392-3.

Russ, A. P., S. Wattler, W. H. Colledge, S. A. Aparicio, M. B. Carlton, J. J. Pearce, S. C. Barton, et al. "Eomesodermin Is Required for Mouse Trophoblast Development and Mesoderm Formation." *Nature* 404, no. 6773 (March 2, 2000): 95–99. https://doi.org/10.1038/35003601.

Schwarz, M. J., N. Müller, D. Körschenhausen, K. H. Kirsch, R. Penning, M. Ackenheil, J. P. Johnson, and H. Hampel. “Melanoma-Associated Adhesion Molecule MUC18/MCAM (CD146) and Transcriptional Regulator Mader in Normal Human CNS.” *Neuroimmunomodulation* 5, no. 5 (October 1998): 270–76. https://doi.org/10.1159/000026347.

Shirasawa, T., T. Akashi, K. Sakamoto, H. Takahashi, N. Maruyama, and K. Hirokawa. “Gene Expression of CD24 Core Peptide Molecule in Developing Brain and Developing Non-Neural Tissues.” *Developmental Dynamics: An Official Publication of the American Association of Anatomists* 198, no. 1 (September 1993): 1–13. https://doi.org/10.1002/aja.1001980102.

Solter, D., and B. B. Knowles. “Monoclonal Antibody Defining a Stage-Specific Mouse Embryonic Antigen (SSEA-1).” *Proceedings of the National Academy of Sciences of the United States of America* 75, no. 11 (November 1978): 5565–69. https://doi.org/10.1073/pnas.75.11.5565.

Storck, T., S. Schulte, K. Hofmann, and W. Stoffel. “Structure, Expression, and Functional Analysis of a Na(+)-Dependent Glutamate/Aspartate Transporter from Rat Brain.” *Proceedings of the National Academy of Sciences of the United States of America* 89, no. 22 (November 15, 1992): 10955–59. https://doi.org/10.1073/pnas.89.22.10955.

Thiery, J P, R Brackenbury, U Rutishauser, and G M Edelman. “Adhesion among Neural Cells of the Chick Embryo. II. Purification and Characterization of a Cell Adhesion Molecule from Neural Retina.” *Journal of Biological Chemistry* 252, no. 19 (October 10, 1977): 6841–45. https://doi.org/10.1016/S0021-9258(17)39926-X.

Uwanogho, D., M. Rex, E. J. Cartwright, G. Pearl, C. Healy, P. J. Scotting, and P. T. Sharpe. “Embryonic Expression of the Chicken Sox2, Sox3 and Sox11 Genes Suggests an Interactive Role in Neuronal Development.” *Mechanisms of Development* 49, no. 1–2 (January 1995): 23–36. https://doi.org/10.1016/0925-4773(94)00299-3.

Uyeda, C. T., L. F. Eng, and A. Bignami. “Immunological Study of the Glial Fibrillary Acidic Protein.” *Brain Research* 37, no. 1 (February 11, 1972): 81–89. https://doi.org/10.1016/0006-8993(72)90347-2.

Vasudevan, Anju, Jason E. Long, James E. Crandall, John L. R. Rubenstein, and Pradeep G. Bhide. “Compartment-Specific Transcription Factors Orchestrate Angiogenesis Gradients in the Embryonic Brain.” *Nature Neuroscience* 11, no. 4 (April 2008): 429–39. https://doi.org/10.1038/nn2074.

Vouyiouklis, D. A., and P. J. Brophy. “Microtubule-Associated Proteins in Developing Oligodendrocytes: Transient Expression of a MAP2c Isoform in Oligodendrocyte Precursors.” *Journal of Neuroscience Research* 42, no. 6 (1995): 803–17. https://doi.org/10.1002/jnr.490420609.

Walther, C., and P. Gruss. “Pax-6, a Murine Paired Box Gene, Is Expressed in the Developing CNS.” *Development (Cambridge, England)* 113, no. 4 (December 1991): 1435–49. https://doi.org/10.1242/dev.113.4.1435.

Weigmann, A., D. Corbeil, A. Hellwig, and W. B. Huttner. “Prominin, a Novel Microvilli-Specific Polytopic Membrane Protein of the Apical Surface of Epithelial Cells, Is Targeted to Plasmalemmal Protrusions of Non-Epithelial Cells.” *Proceedings of the National Academy of Sciences of the United States of America* 94, no. 23 (November 11, 1997): 12425–30. https://doi.org/10.1073/pnas.94.23.12425.

Yamamoto, M., M. Fujinuma, M. Tanaka, U. C. Dräger, and P. McCaffery. “Sagittal Band Expression of COUP-TF2 Gene in the Developing Cerebellum.” *Mechanisms of Development* 84, no. 1–2 (June 1999): 143–46. https://doi.org/10.1016/s0925-4773(99)00054-4.

Yokoyama, Akiko, Lihua Yang, Suzuka Itoh, Kohji Mori, and Junya Tanaka. “Microglia, a Potential Source of Neurons, Astrocytes, and Oligodendrocytes.” *Glia* 45, no. 1 (2004): 96–104. https://doi.org/10.1002/glia.10306.

Zhou, Q., S. Wang, and D. J. Anderson. “Identification of a Novel Family of Oligodendrocyte Lineage-Specific Basic Helix-Loop-Helix Transcription Factors.” *Neuron* 25, no. 2 (February 2000): 331–43. https://doi.org/10.1016/s0896-6273(00)80898-3.
