## Supplementary Table 1 for "A developmental atlas of the mouse brain by single-cell mass cytometry"

**Supplementary Table 1.** Sample metadata.

| FCS File Name | Run | BC Set | Age | Tissue | Litter ID | No. Pooled Animals | Ungated | Export | Yield (%) | CD45+ Cells | CD45+ Cells (%) | Sox2/ Nestin+ Cells | Sox2/ Nestin+ Cells (%) |
|---|---|---|---|---|---|---|---|---|---|---|---|---|---|
| BCSet1_E11_Brain.fcs | 1 | 1 | E11.5 | Whole Brain | E11-20180205-1 | 5 | 356990 | 220031 | 61.64% | 8401 | 3.82% | 43471 | 19.76% |
| BCSet1_E12_Brain.fcs | 1 | 1 | E12.5 | Whole Brain | E12-20180109-1 | 6 | 274834 | 162068 | 58.97% | 4081 | 2.52% | 62170 | 38.36% |
| BCSet1_E13_Cortex.fcs | 1 | 1 | E13.5 | Telencephalon | E13-20180821-1 | 4 | 282560 | 177349 | 62.77% | 1470 | 0.83% | 316 | 0.18% |
| BCSet1_E13_Diencephalon.fcs | 1 | 1 | E13.5 | Diencephalon | E13-20180821-1 | 4 | 343286 | 237347 | 69.14% | 2144 | 0.90% | 7194 | 3.03% |
| BCSet1_E14_Cortex.fcs | 1 | 1 | E14.5 | Telencephalon | E14-20180216-1 | 2 | 323720 | 198057 | 61.18% | 2210 | 1.12% | 117827 | 59.49% |
| BCSet1_E14_Diencephalon.fcs | 1 | 1 | E14.5 | Diencephalon | E14-20180216-1 | 2 | 174637 | 113995 | 65.28% | 1609 | 1.41% | 47927 | 42.04% |
| BCSet1_E15_Cortex.fcs | 1 | 1 | E15.5 | Telencephalon | E15-20180221-1 | 4 | 751626 | 511330 | 68.03% | 5743 | 1.12% | 104877 | 20.51% |
| BCSet1_E15_Diencephalon.fcs | 1 | 1 | E15.5 | Diencephalon | E15-20180221-1 | 4 | 208665 | 137360 | 65.83% | 7503 | 5.46% | 33029 | 24.05% |
| BCSet1_E16_Cortex.fcs | 1 | 1 | E16.5 | Telencephalon | E16-20180222-2 | 8 | 1210839 | 862822 | 71.26% | 13290 | 1.54% | 193987 | 22.48% |
| BCSet1_E16_Diencephalon.fcs | 1 | 1 | E16.5 | Diencephalon | E16-20180222-2 | 8 | 250630 | 165487 | 66.03% | 12937 | 7.82% | 33103 | 20.00% |
| BCSet1_E17_Cortex.fcs | 1 | 1 | E17.5 | Telencephalon | E17-20180219-1 | 7 | 818591 | 570018 | 69.63% | 37038 | 6.50% | 14726 | 2.58% |
| BCSet1_E17_Diencephalon.fcs | 1 | 1 | E17.5 | Diencephalon | E17-20180219-1 | 7 | 120081 | 61996 | 51.63% | 6381 | 10.29% | 8966 | 14.46% |
| BCSet1_E18_Cortex.fcs | 1 | 1 | E18.5 | Telencephalon | E18-20180220-1 | 11 | 857450 | 591345 | 68.97% | 26330 | 4.45% | 514 | 0.09% |
| BCSet1_E18_Diencephalon.fcs | 1 | 1 | E18.5 | Diencephalon | E18-20180220-1 | 11 | 89535 | 46221 | 51.62% | 5710 | 12.35% | 5637 | 12.20% |
| BCSet1_P0_Cortex.fcs | 1 | 1 | P0 | Telencephalon | P0-20180829-1 | 8 | 674707 | 466050 | 69.07% | 37723 | 8.09% | 1791 | 0.38% |
| BCSet1_P0_Diencephalon.fcs | 1 | 1 | P0 | Diencephalon | P0-20180829-1 | 8 | 68755 | 13688 | 19.91% | 1813 | 13.25% | 693 | 5.06% |
| BCSet2_E13_Hindbrain.fcs | 1 | 2 | E13.5 | Rhombencephalon | E13-20180821-1 | 4 | 374839 | 211327 | 56.38% | 7811 | 3.70% | 11079 | 5.24% |
| BCSet2_E13_Midbrain.fcs | 1 | 2 | E13.5 | Mesencephalon | E13-20180821-1 | 4 | 806721 | 597751 | 74.10% | 11060 | 1.85% | 116801 | 19.54% |
| BCSet2_E14_Hindbrain.fcs | 1 | 2 | E14.5 | Rhombencephalon | E14-20180216-1 | 2 | 300040 | 169719 | 56.57% | 6723 | 3.96% | 40936 | 24.12% |
| BCSet2_E14_Midbrain.fcs | 1 | 2 | E14.5 | Mesencephalon | E14-20180216-1 | 2 | 362876 | 211906 | 58.40% | 2908 | 1.37% | 39891 | 18.82% |
| BCSet2_E15_Hindbrain.fcs | 1 | 2 | E15.5 | Rhombencephalon | E15-20180221-1 | 4 | 456327 | 318170 | 69.72% | 34851 | 10.95% | 58871 | 18.50% |
| BCSet2_E15_Midbrain.fcs | 1 | 2 | E15.5 | Mesencephalon | E15-20180221-1 | 4 | 480185 | 325244 | 67.73% | 8722 | 2.68% | 33474 | 10.29% |
| BCSet2_E16_Hindbrain.fcs | 1 | 2 | E16.5 | Rhombencephalon | E16-20180222-2 | 8 | 376559 | 273434 | 72.61% | 16824 | 6.15% | 48424 | 17.71% |
| BCSet2_E16_Midbrain.fcs | 1 | 2 | E16.5 | Mesencephalon | E16-20180222-2 | 8 | 368925 | 232622 | 63.05% | 7657 | 3.29% | 28128 | 12.09% |
| BCSet2_E17_Hindbrain.fcs | 1 | 2 | E17.5 | Rhombencephalon | E17-20180219-1 | 7 | 221931 | 133163 | 60.00% | 5084 | 3.82% | 6203 | 4.66% |
| BCSet2_E17_Midbrain.fcs | 1 | 2 | E17.5 | Mesencephalon | E17-20180219-1 | 7 | 243946 | 151528 | 62.12% | 20305 | 13.40% | 17500 | 11.55% |
| BCSet2_E18_Hindbrain.fcs | 1 | 2 | E18.5 | Rhombencephalon | E18-20180220-1 | 11 | 597326 | 448823 | 75.14% | 42170 | 9.40% | 1603 | 0.36% |
| BCSet2_E18_Midbrain.fcs | 1 | 2 | E18.5 | Mesencephalon | E18-20180220-1 | 11 | 371257 | 233669 | 62.94% | 30662 | 13.12% | 12961 | 5.55% |
| BCSet2_P0_Hindbrain.fcs | 1 | 2 | P0 | Rhombencephalon | P0-20180220-1 | 6 | 585780 | 427537 | 72.99% | 27917 | 6.53% | 111 | 0.03% |
| BCSet2_P0_Midbrain.fcs | 1 | 2 | P0 | Mesencephalon | P0-20180220-1 | 6 | 310088 | 193306 | 62.34% | 27404 | 14.18% | 24638 | 12.75% |
| BCSet3_P1_Cortex.fcs | 1 | 3 | P1 | Telencephalon | P1-20180807-1 | 4 | 963487 | 609358 | 63.25% | 55312 | 9.08% | 402 | 0.07% |
| BCSet3_P1_Diencephalon.fcs | 1 | 3 | P1 | Diencephalon | P1-20180807-1 | 4 | 544880 | 373186 | 68.49% | 37690 | 10.10% | 66815 | 17.90% |
| BCSet3_P1_Hindbrain.fcs | 1 | 3 | P1 | Rhombencephalon | P1-20180807-1 | 4 | 379166 | 293705 | 77.46% | 20771 | 7.07% | 22598 | 7.69% |
| BCSet3_P1_Midbrain.fcs | 1 | 3 | P1 | Mesencephalon | P1-20180807-1 | 4 | 482847 | 341430 | 70.71% | 55177 | 16.16% | 1928 | 0.56% |
| BCSet3_P2_Cortex.fcs | 1 | 3 | P2 | Telencephalon | P2-20180808-1 | 5 | 469807 | 318403 | 67.77% | 37040 | 11.63% | 34549 | 10.85% |
| BCSet3_P2_Diencephalon.fcs | 1 | 3 | P2 | Diencephalon | P2-20180808-1 | 5 | 469694 | 328469 | 69.93% | 41940 | 12.77% | 49678 | 15.12% |
| BCSet3_P2_Hindbrain.fcs | 1 | 3 | P2 | Rhombencephalon | P2-20180808-1 | 5 | 340095 | 241581 | 71.03% | 17542 | 7.26% | 18859 | 7.81% |
| BCSet3_P2_Midbrain.fcs | 1 | 3 | P2 | Mesencephalon | P2-20180808-1 | 5 | 530548 | 367259 | 69.22% | 66978 | 18.24% | 34236 | 9.32% |
| BCSet3_P3_Cortex.fcs | 1 | 3 | P3 | Telencephalon | P3-20180809-1 | 3 | 430002 | 275983 | 64.18% | 42719 | 15.48% | 48820 | 17.69% |
| BCSet3_P3_Diencephalon.fcs | 1 | 3 | P3 | Diencephalon | P3-20180809-1 | 3 | 412967 | 291234 | 70.52% | 41483 | 14.24% | 62347 | 21.41% |
| BCSet3_P3_Hindbrain.fcs | 1 | 3 | P3 | Rhombencephalon | P3-20180809-1 | 3 | 409400 | 317626 | 77.58% | 21706 | 6.83% | 2007 | 0.63% |
| BCSet3_P3_Midbrain.fcs | 1 | 3 | P3 | Mesencephalon | P3-20180809-1 | 3 | 433281 | 295148 | 68.12% | 55435 | 18.78% | 39907 | 13.52% |
| BCSet3_P4_Cortex.fcs | 1 | 3 | P4 | Telencephalon | P4-20180225-1 | 6 | 382818 | 241213 | 63.01% | 42321 | 17.55% | 62251 | 25.81% |
| BCSet3_P4_Diencephalon.fcs | 1 | 3 | P4 | Diencephalon | P4-20180225-1 | 6 | 474334 | 338974 | 71.46% | 47809 | 14.10% | 51436 | 15.17% |
| BCSet3_P4_Hindbrain.fcs | 1 | 3 | P4 | Rhombencephalon | P4-20180225-1 | 6 | 542755 | 423797 | 78.08% | 21443 | 5.06% | 10383 | 2.45% |
| BCSet3_P4_Midbrain.fcs | 1 | 3 | P4 | Mesencephalon | P4-20180225-1 | 6 | 336269 | 228522 | 67.96% | 47206 | 20.66% | 37319 | 16.33% |
| BCSet4_E11_Brain.fcs | 2 | 4 | E11.5 | Whole Brain | E11-20190129-1 | 4 | 166854 | 62449 | 37.43% | 2337 | 3.74% | 28089 | 44.98% |
| BCSet4_E12_Brain.fcs | 2 | 4 | E12.5 | Whole Brain | E12-20180221-2 | 8 | 488763 | 273420 | 55.94% | 8823 | 3.23% | 99530 | 36.40% |
| BCSet4_E13_Cortex.fcs | 2 | 4 | E13.5 | Telencephalon | E13-20180131-1 | 8 | 142454 | 57147 | 40.12% | 1063 | 1.86% | 21593 | 37.79% |
| BCSet4_E13_Diencephalon.fcs | 2 | 4 | E13.5 | Diencephalon | E13-20180131-1 | 8 | 124852 | 54790 | 43.88% | 520 | 0.95% | 462 | 0.84% |
| BCSet4_E14_Cortex.fcs | 2 | 4 | E14.5 | Telencephalon | E14-20180208-1 | 7 | 278051 | 146472 | 52.68% | 853 | 0.58% | 76179 | 52.01% |
| BCSet4_E14_Diencephalon.fcs | 2 | 4 | E14.5 | Diencephalon | E14-20180208-1 | 7 | 104789 | 54875 | 52.37% | 637 | 1.16% | 17959 | 32.73% |
| BCSet4_E15_Cortex.fcs | 2 | 4 | E15.5 | Telencephalon | E15-20180119-1 | 9 | 458206 | 222149 | 48.48% | 2416 | 1.09% | 91499 | 41.19% |
| BCSet4_E15_Diencephalon.fcs | 2 | 4 | E15.5 | Diencephalon | E15-20180119-1 | 9 | 75874 | 33313 | 43.91% | 938 | 2.82% | 7502 | 22.52% |
| BCSet4_E16_Cortex.fcs | 2 | 4 | E16.5 | Telencephalon | E16-20181006-1 | 5 | 517763 | 311349 | 60.13% | 6881 | 2.21% | 52950 | 17.01% |
| BCSet4_E16_Diencephalon.fcs | 2 | 4 | E16.5 | Diencephalon | E16-20181006-1 | 5 | 193964 | 124259 | 64.06% | 7730 | 6.22% | 19212 | 15.46% |
| BCSet4_E17_Cortex.fcs | 2 | 4 | E17.5 | Telencephalon | E17-20181021-1 | 10 | 334333 | 186925 | 55.91% | 2923 | 1.56% | 28407 | 15.20% |
| BCSet4_E17_Diencephalon_2.fcs | 2 | 4 | E17.5 | Diencephalon | E17-20181007-1 | 9 | 188959 | 111841 | 59.19% | 11796 | 7.32% | 18081 | 16.17% |
| BCSet4_E17_Diencephalon.fcs | 2 | 4 | E17.5 | Diencephalon | E17-20181021-1 | 10 | 181663 | 101245 | 55.73% | 8182 | 11.65% | 16458 | 16.26% |
| BCSet4_E18_Cortex.fcs | 2 | 4 | E18.5 | Telencephalon | E18-20180220-2 | 7 | 290774 | 132769 | 45.66% | 6249 | 4.71% | 8026 | 6.05% |
| BCSet4_E18_Diencephalon_2.fcs | 2 | 4 | E18.5 | Diencephalon | E18-20181015-1 | 5 | 103487 | 44723 | 43.22% | 4257 | 15.41% | 1282 | 2.87% |
| BCSet4_E18_Diencephalon.fcs | 2 | 4 | E18.5 | Diencephalon | E18-20180220-2 | 7 | 114631 | 58567 | 51.09% | 6894 | 7.27% | 5880 | 10.04% |
| BCSet4_P0_Cortex.fcs | 2 | 4 | P0 | Telencephalon | P0-20181015-1 | 6 | 329225 | 173746 | 52.77% | 10068 | 5.79% | 23936 | 13.78% |
| BCSet4_P0_Diencephalon_2.fcs | 2 | 4 | P0 | Diencephalon | P0-20180220-1 | 6 | 58941 | 24879 | 42.21% | 3062 | 29.62% | 4908 | 19.73% |
| BCSet4_P0_Diencephalon.fcs | 2 | 4 | P0 | Diencephalon | P0-20181015-1 | 6 | 167232 | 100779 | 60.26% | 7370 | 3.04% | 20490 | 20.33% |
| BCSet5_E13_Hindbrain.fcs | 2 | 5 | E13.5 | Rhombencephalon | E13-20180131-1 | 8 | 47942 | 18794 | 39.20% | 1479 | 7.87% | 4361 | 23.20% |
| BCSet5_E13_Midbrain.fcs | 2 | 5 | E13.5 | Mesencephalon | E13-20180131-1 | 8 | 46860 | 22495 | 48.00% | 673 | 2.99% | 4106 | 18.25% |
| BCSet5_E14_Hindbrain.fcs | 2 | 5 | E14.5 | Rhombencephalon | E14-20180208-1 | 7 | 38852 | 16030 | 41.26% | 1491 | 9.30% | 4392 | 27.40% |
| BCSet5_E14_Midbrain.fcs | 2 | 5 | E14.5 | Mesencephalon | E14-20180208-1 | 7 | 204360 | 115796 | 56.66% | 1210 | 1.04% | 29527 | 25.50% |
| BCSet5_E15_Hindbrain.fcs | 2 | 5 | E15.5 | Rhombencephalon | E15-20180119-1 | 9 | 54881 | 25574 | 46.60% | 3522 | 13.77% | 7692 | 30.08% |
| BCSet5_E15_Midbrain.fcs | 2 | 5 | E15.5 | Mesencephalon | E15-20180119-1 | 9 | 31837 | 11116 | 34.92% | 299 | 2.69% | 1367 | 12.30% |
| BCSet5_E16_Hindbrain.fcs | 2 | 5 | E16.5 | Rhombencephalon | E16-20181006-1 | 5 | 56471 | 31098 | 55.07% | 3305 | 10.63% | 3323 | 10.69% |
| BCSet5_E16_Midbrain.fcs | 2 | 5 | E16.5 | Mesencephalon | E16-20181006-1 | 5 | 50818 | 15863 | 31.22% | 644 | 4.06% | 1821 | 11.48% |
| BCSet5_E17_Hindbrain_2.fcs | 2 | 5 | E17.5 | Rhombencephalon | E17-20181007-1 | 9 | 274385 | 183543 | 66.89% | 18237 | 6.74% | 15748 | 8.58% |
| BCSet5_E17_Hindbrain.fcs | 2 | 5 | E17.5 | Rhombencephalon | E17-20181021-1 | 10 | 314879 | 212628 | 67.53% | 12372 | 8.58% | 14146 | 6.65% |
| BCSet5_E17_Midbrain.fcs | 2 | 5 | E17.5 | Mesencephalon | E17-20181021-1 | 10 | 191524 | 106580 | 55.65% | 6482 | 6.08% | 10886 | 10.21% |
| BCSet5_E18_Hindbrain.fcs | 2 | 5 | E18.5 | Rhombencephalon | E18-20180220-2 | 7 | 328607 | 190663 | 58.02% | 18775 | 9.85% | 2675 | 1.40% |
| BCSet5_E18_Midbrain.fcs | 2 | 5 | E18.5 | Mesencephalon | E18-20180220-2 | 7 | 115466 | 60205 | 52.14% | 10237 | 17.00% | 7023 | 11.67% |
| BCSet5_P0_Hindbrain.fcs | 2 | 5 | P0 | Rhombencephalon | P0-20181015-1 | 6 | 304472 | 210280 | 69.06% | 16961 | 8.07% | 11129 | 5.29% |
| BCSet5_P0_Midbrain.fcs | 2 | 5 | P0 | Mesencephalon | P0-20181015-1 | 6 | 167655 | 92753 | 55.32% | 13148 | 14.18% | 10601 | 11.43% |
| BCSet6_P1_Cortex.fcs | 2 | 6 | P1 | Telencephalon | P1-20180807-2 | 7 | 512111 | 283669 | 55.39% | 31685 | 11.17% | 57731 | 20.35% |
| BCSet6_P1_Diencephalon.fcs | 2 | 6 | P1 | Diencephalon | P1-20180807-2 | 7 | 440601 | 255431 | 57.97% | 44195 | 17.30% | 43135 | 16.89% |
| BCSet6_P1_Hindbrain.fcs | 2 | 6 | P1 | Rhombencephalon | P1-20180807-2 | 7 | 483670 | 307038 | 63.48% | 23973 | 7.81% | 29132 | 9.49% |
| BCSet6_P1_Midbrain.fcs | 2 | 6 | P1 | Mesencephalon | P1-20180807-2 | 7 | 140621 | 64796 | 46.08% | 12970 | 20.02% | 11916 | 18.39% |
| BCSet6_P2_Cortex.fcs | 2 | 6 | P2 | Telencephalon | P2-20180808-2 | 3 | 313253 | 174849 | 55.82% | 21520 | 12.31% | 32374 | 18.52% |
| BCSet6_P2_Diencephalon.fcs | 2 | 6 | P2 | Diencephalon | P2-20180808-2 | 3 | 403087 | 254374 | 63.11% | 27746 | 10.91% | 34834 | 13.69% |
| BCSet6_P2_Hindbrain.fcs | 2 | 6 | P2 | Rhombencephalon | P2-20180808-2 | 3 | 367793 | 244641 | 66.52% | 17342 | 7.09% | 14832 | 6.06% |
| BCSet6_P2_Midbrain.fcs | 2 | 6 | P2 | Mesencephalon | P2-20180808-2 | 3 | 183554 | 103644 | 56.47% | 19513 | 18.83% | 15940 | 15.38% |
| BCSet6_P3_Cortex.fcs | 2 | 6 | P3 | Telencephalon | P3-20180809-2 | 3 | 332533 | 177979 | 53.52% | 24608 | 13.83% | 17671 | 9.93% |
| BCSet6_P3_Diencephalon.fcs | 2 | 6 | P3 | Diencephalon | P3-20180809-2 | 3 | 280885 | 165460 | 58.91% | 16597 | 10.03% | 7467 | 4.51% |
| BCSet6_P3_Hindbrain.fcs | 2 | 6 | P3 | Rhombencephalon | P3-20180809-2 | 3 | 523719 | 333766 | 63.73% | 18727 | 5.61% | 704 | 0.21% |
| BCSet6_P3_Midbrain.fcs | 2 | 6 | P3 | Mesencephalon | P3-20180809-2 | 3 | 87504 | 36631 | 41.86% | 6417 | 17.52% | 1956 | 5.34% |
| BCSet6_P4_Cortex.fcs | 2 | 6 | P4 | Telencephalon | P4-20181008-1 | 5 | 307179 | 166690 | 54.26% | 26713 | 16.03% | 36170 | 21.70% |
| BCSet6_P4_Diencephalon.fcs | 2 | 6 | P4 | Diencephalon | P4-20181008-1 | 5 | 357155 | 230665 | 64.58% | 29732 | 12.89% | 38325 | 16.62% |
| BCSet6_P4_Hindbrain.fcs | 2 | 6 | P4 | Rhombencephalon | P4-20181008-1 | 5 | 653168 | 435469 | 66.67% | 22276 | 5.12% | 9873 | 2.27% |
| BCSet6_P4_Midbrain.fcs | 2 | 6 | P4 | Mesencephalon | P4-20181008-1 | 5 | 161400 | 94010 | 58.25% | 13456 | 14.31% | 11762 | 12.51% |
| BCSet7_E13_Hindbrain.fcs | 3 | 7 | E13.5 | Rhombencephalon | E13-20180821-2 | 7 | 201984 | 121752 | 60.28% | 6964 | 5.72% | 16649 | 13.67% |
| BCSet7_E13_Midbrain.fcs | 3 | 7 | E13.5 | Mesencephalon | E13-20180821-2 | 7 | 439565 | 210562 | 47.90% | 6174 | 2.93% | 20793 | 9.88% |
| BCSet7_E14_Hindbrain.fcs | 3 | 7 | E14.5 | Rhombencephalon | E14-20180208-1 | 7 | 272845 | 168231 | 61.66% | 18542 | 11.02% | 37088 | 22.05% |
| BCSet7_E15_Diencephalon.fcs | 3 | 7 | E15.5 | Diencephalon | E15-20180216-1 | 5 | 607576 | 446974 | 73.57% | 13172 | 2.95% | 116474 | 26.06% |
| BCSet7_E15_Hindbrain.fcs | 3 | 7 | E15.5 | Rhombencephalon | E15-20180216-1 | 5 | 320838 | 236699 | 73.78% | 22336 | 9.44% | 41088 | 17.36% |
| BCSet7_E15_Midbrain.fcs | 3 | 7 | E15.5 | Mesencephalon | E15-20180216-1 | 5 | 456778 | 299600 | 65.59% | 11470 | 3.83% | 34616 | 11.55% |
| BCSet7_E16_Hindbrain.fcs | 3 | 7 | E16.5 | Rhombencephalon | E16-20181006-2 | 9 | 232090 | 147612 | 63.60% | 13422 | 9.09% | 25898 | 17.54% |
| BCSet7_E16_Midbrain.fcs | 3 | 7 | E16.5 | Mesencephalon | E16-20181006-2 | 9 | 390998 | 253297 | 64.78% | 10969 | 4.33% | 28989 | 11.44% |
| BCSet7_E17_Diencephalon.fcs | 3 | 7 | E17.5 | Diencephalon | E17-20180219-2 | 10 | 258698 | 147048 | 56.84% | 12096 | 8.23% | 28725 | 19.53% |
| BCSet7_E18_Diencephalon.fcs | 3 | 7 | E18.5 | Diencephalon | E18-20180220-1 | 11 | 451887 | 321729 | 71.20% | 24787 | 7.70% | 46118 | 14.33% |
| BCSet7_P0_Diencephalon.fcs | 3 | 7 | P0 | Diencephalon | P0-20190123-1 | 4 | 404667 | 278098 | 68.72% | 19609 | 7.05% | 62902 | 22.62% |
| BCSet7_P1_Diencephalon.fcs | 3 | 7 | P1 | Diencephalon | P1-20171227-1 | 3 | 259809 | 143354 | 55.18% | 11670 | 8.14% | 24439 | 17.05% |
| BCSet7_P1_Midbrain.fcs | 3 | 7 | P1 | Mesencephalon | P1-20171227-1 | 3 | 133971 | 70057 | 52.29% | 14402 | 20.56% | 11479 | 16.39% |
| BCSet7_P3_Diencephalon.fcs | 3 | 7 | P3 | Diencephalon | P3-20180809-1 | 3 | 366790 | 240052 | 65.45% | 33566 | 13.98% | 41929 | 17.47% |
| BCSet7_P3_Hindbrain.fcs | 3 | 7 | P3 | Rhombencephalon | P3-20180809-2 | 3 | 633149 | 475487 | 75.10% | 29064 | 6.11% | 16614 | 3.49% |
| BCSet7_P3_Midbrain.fcs | 3 | 7 | P3 | Mesencephalon | P3-20180809-1 | 3 | 117667 | 63048 | 53.58% | 12603 | 19.99% | 4395 | 6.97% |
